# Distinguishing dopamine and calcium responses using XNA-nanotube sensors for improved neurochemical sensing

**DOI:** 10.1101/2021.02.20.428669

**Authors:** Alice J. Gillen, Alessandra Antonucci, Melania Reggente, Daniel Morales, Ardemis A. Boghossian

## Abstract

To date, the engineering of single-stranded DNA-SWCNT (DNA-SWCNT) optical biosensors have largely focused on creating sensors for new applications with little focus on optimising existing sensors for *in vitro* and *in vivo* conditions. Recent studies have shown that nanotube fluorescence can be severely impacted by changes in local cation concentrations. This is particularly problematic for neurotransmitter sensing applications as spatial and temporal fluctuations in the concentration of cations, such as Na^+^, K^+^, or Ca^2+^, play a central role in neuromodulation. This can lead to inaccuracies in the determination of neurotransmitter concentrations using DNA-SWCNT sensors, which limits their use for detecting and treating neurological diseases.

Herein, we present new approaches using locked nucleic acid (LNA) to engineer SWCNT sensors with improved stability towards cation-induced fluorescence changes. By incorporating LNA bases into the (GT)_15_-DNA sequence, we create sensors that are not only more resistant towards undesirable fluorescence modulation in the presence of Ca^2+^ but that also retain their capabilities for the label-free detection of dopamine. The synthetic biology approach presented in this work therefore serves as a complementary means for enhancing nanotube optoelectronic behavior, unlocking previously unexplored possibilities for developing nano-bioengineered sensors with augmented capabilities.

## Introduction

Neurotransmitters play a central role in a variety of biological functions that rely on chemical communication^1,2^. Imbalances or signalling problems of key neurotransmitters, such as dopamine, serotonin, and γ-aminobutyric acid (GABA)^2–5^, are often linked to the pathology of many neurological diseases, including Alzheimer’s and Parkinson’s disease. The prevalence of these diseases is unfortunately expected to increase as the global population ages^6^, contributing to not only greater mortality, but also a severely compromised quality of life. State-of-the-art treatments could be improved by real-time measurements of neurotransmitter concentrations, however the current lack of selective, responsive, and minimally invasive technologies limits personalized treatment options and patient comfort.

Several of these limitations have been overcome with the development of optical biosensors based on single-walled carbon nanotubes (SWCNTs). SWCNTs can be conceptualised as cylindrically rolled sheets of graphene whose optoelectronic properties vary with diameter and chirality. The semiconducting chiralities show distinct intrinsic fluorescence emissions that enable multimodal sensing in mixed chirality samples. Furthermore, these emissions span the second near-infrared optical window, which overlaps with the optical transparency window for biological tissue. This optical overlap, along with the SWCNT’s resistance to photobleaching, makes SWCNTs suitable for deep-tissue, long-term, and continuous *in vitro* and *in vivo* imaging^7^. SWCNTs have also shown high spatial and temporal resolution, which is particularly important for the real-time, spatiotemporal monitoring of neurotransmitters^8,9^. The selectivity preferences of these SWCNT-based optical sensors toward neurotransmitters, such as dopamine^9–12^ and serotonin^13,14^ can be modulated by non-covalently functionalizing the SWCNT surface with specific sequences of single-stranded DNA (DNA). Following either direct sonication or dialysis exchange, the DNA has been shown to self-assemble onto the SWCNT surface through *π-π* stacking of the bases (**Figure 1A**). In addition to solubilizing the nanotubes, the DNA can alter the nanotube fluorescence response towards specific analytes in a sequence- and length-dependent manner, thus enabling preferential selectivity towards analytes of interest. Such optical SWCNT sensors have been used for *in vivo* brain imaging and *in vitro* spatiotemporal dopamine mapping of neurons^9,11,15–17^.

**Figure 1:**
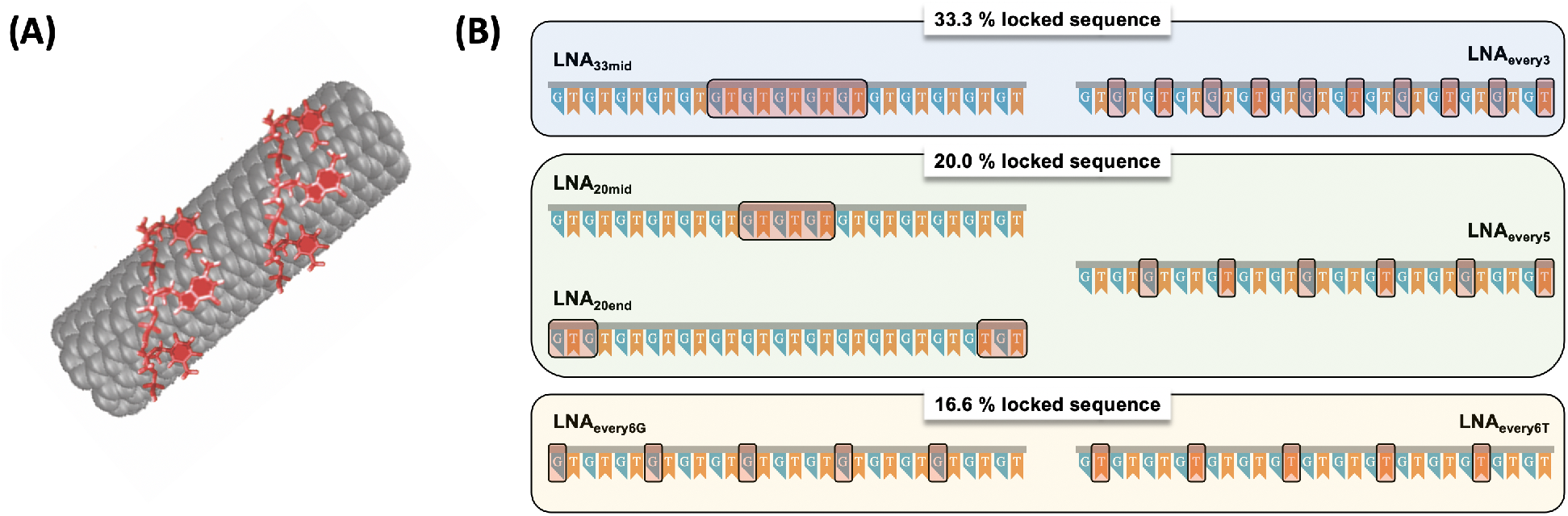
**(A)** Schematic of oligonucleotide wrapping on SWCNTs surface via π-π stacking. **(B)** Schematic of all single-stranded LNA(GT)_15_ sequences tested in this study. Locked bases are indicated by a red-shaded box.

However, recent studies have shown that nanotube fluorescence can be severely impacted by changes in local cation concentrations^18,19^. This sensitivity to fluctuating cation concentrations is particularly detrimental to neurotransmitter sensing applications, where cells communicate via changes in concentrations of neurotransmitters and neuromodulators that are often triggered by fluxes of cations such as Ca^2+^^1,20–23^. In fact, the competitive responsivity of many existing DNA-SWCNT dopamine sensors to such cations limits the application of these sensors in efforts to untangle the complex signalling dynamics of the brain^11,18,19^. Consequently, the real-time, selective detection of physiological concentrations of dopamine in the absence of ionic or salt interference remains an ongoing challenge.

Herein, we present a SWCNT-based sensor capable of detecting dopamine with suppressed cation responsivity. We bioengineer a DNA coating, (GT)_15_, which has previously been shown to respond to both dopamine^10^ and various cations^19^, by incorporating locked nucleic acid (LNA) bases at various positions throughout the sequence. We systematically characterized the different LNA wrappings, examining their suspension quality, morphologies, and optoelectronic effects on sensor response to both cations and dopamine. The LNA serves to not only suppress the sensor’s competitive response to cations, but also sustains the sensor’s selective dopamine responsivity, allowing us to demonstrate dopamine detection in the presence of multiple cationic species. Furthermore, via concomitant monitoring of multiple chiralities, we showed that certain LNA sequences enable, for the first time, the selective detection of both Ca^2+^ and dopamine simultaneously.

## Results and Discussion

We compared the solubilisation and fluorescence properties of the original DNA, (GT)_15_, to those of seven distinct single-stranded LNA-based derivatives (**Figure 1B**). As summarized in **Figure 1B**, the designed LNA sequences differed in terms of the percentage of locking, the type of base locked (either T or G), and the location and distribution of locking (detailed sequence information is provided in **Table S1**). All tested LNA sequences were able to suspend the SWCNTs to varying extents, as evidenced by the dark solutions obtained post dialysis (**Figure SI.1**) as well as from the bright fluorescence peaks and distinct bands in the absorption spectrum for each of the solutions (**Figure 2 (A)** and **(B)**, **SI.2**).

**Figure 2:**
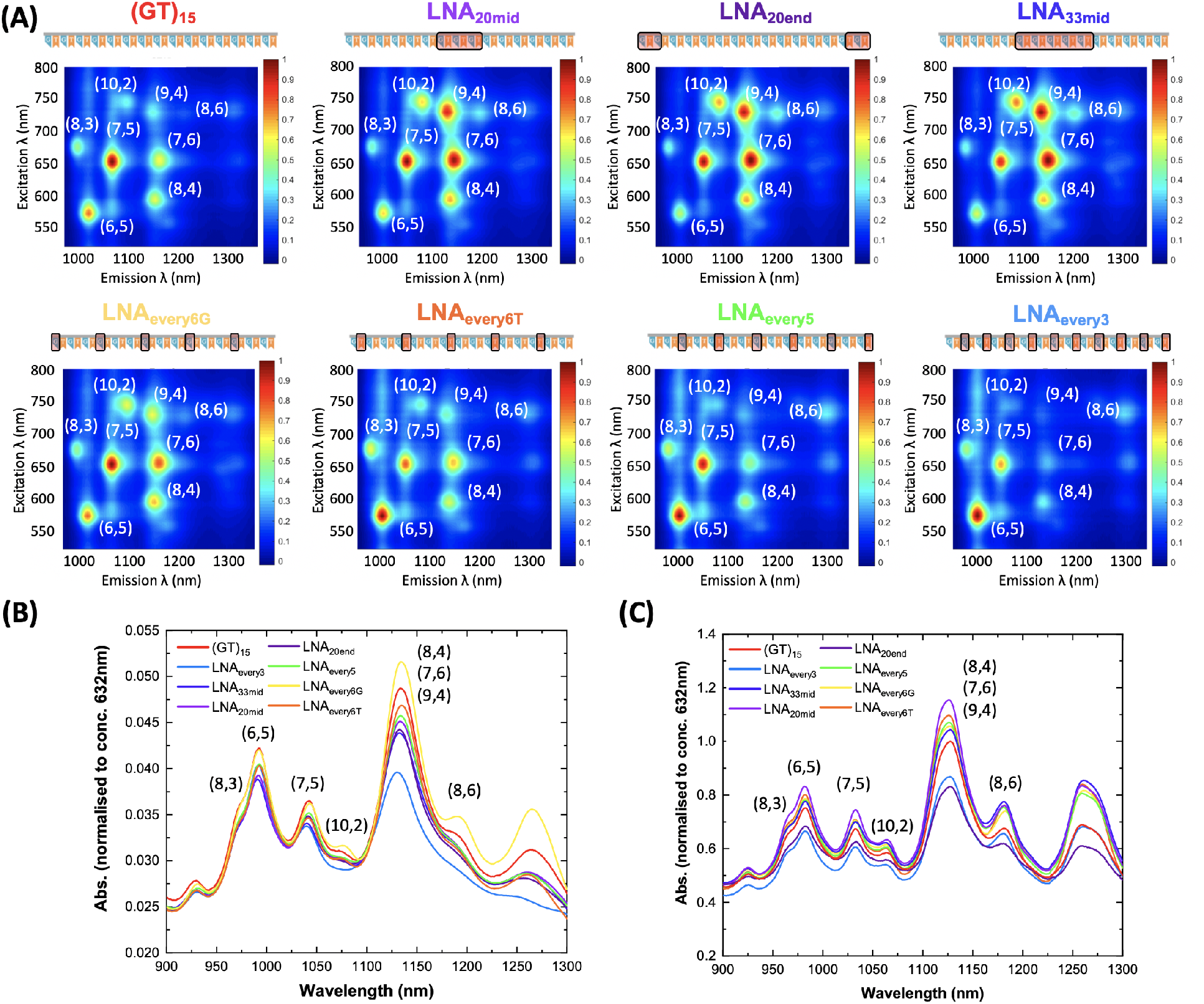
Suspension quality and chirality specificity of LNA sequences. **(A)** PLE maps of the original (GT)_15_- and all LNA-SWCNT solutions. **(B)** Absorbance spectra for all DNA- and LNA-SWCNT samples examined in this study. All spectra are normalized to concentration as determined using an extinction coefficient of 0.036 mg/L at Abs632nm. **(C)** Absorbance spectra of all sequences following SDOC replacement of original solutions collected following 16 h of incubation. Spectra were normalized to nanotube concentration as determined using an extinction coefficient of 0.036 mg/L at Abs_632nm_ and subsequently normalized to the (GT)_15_ spectrum (**red**) for comparison.

As shown in the photoluminescence excitation/emission (PLE) maps (**Figure 2 (A)**), the LNA-SWCNTs showed preferential, chirality-specific fluorescence emissions, depending on the locking positions of the sequence. The (GT)_15_ SWCNTs showed highest fluorescence emissions for the (7,5) nanotube chirality at 660 nm excitation, as expected for HiPco SWCNT samples^19^. In contrast, LNA sequences that contained blocks of consecutively locked bases (LNA_20mid_, LNA_20end_ and LNA_33mid_) showed higher fluorescence emissions from the larger diameter chiralities that appear at longer excitation and emissions wavelengths. On the other hand, the fluorescence was more pronounced for small diameter nanotubes, such as the (6,5) chirality, for sequences with an increased locking periodicity (LNA_every6T_, LNA_every5_ and LNA_every3_).

To determine whether the chirality-specific differences in fluorescence emissions were due to selective increases in quantum yield or chirality enrichment, we compared the chirality distribution of the nanotubes using absorption spectroscopy (**Figure 2 (B)**). Although all LNA-SWCNT suspensions were synthesized from the same initial distribution of SWCNT chiralities, differences in the absorption peak ratios for the different LNA sequences suggests that that the LNA preferentially enriches certain chiralities. Chirality enrichment was further confirmed by displacing all LNA and DNA wrappings with the same wrapping, sodium deoxycholate (SDOC)^24^. Following equilibration with the preferentially adsorbed SDOC (**Figure SI.3, SI.4, SI.5**), we observed that none of the SDOC-suspended SWCNTs had overlapping absorbance spectra (**Figure 2 (C)**). These observations confirm that the LNA is not simply altering the quantum yield in a chiral-dependent manner but rather that the different LNA sequences exhibit preferences for suspending certain nanotube chiralities.

The wrapping morphology of the different oligonucleotide sequences was subsequently characterized using atomic force microscopy (AFM) (**Figure 3**, **SI.7 – SI.10**). Because of the increased vertical (compared to lateral) resolution of AFM measurements, we examined the differences between the wrappings by comparing coated (peaks, **Figure 3** (**center**)) and uncoated (valleys, **Figure 3** (**right**)) regions along the nanotube. This enabled us to examine the wrapping homogeneity along the nanotube for the different sequences by comparing the variance in the height distribution of the peaks (**Figure 3** (**left**)). We could also qualitatively assess the periodicity of the wrappings by counting the number of peaks observed per unit length along the nanotube.

**Figure 3:**
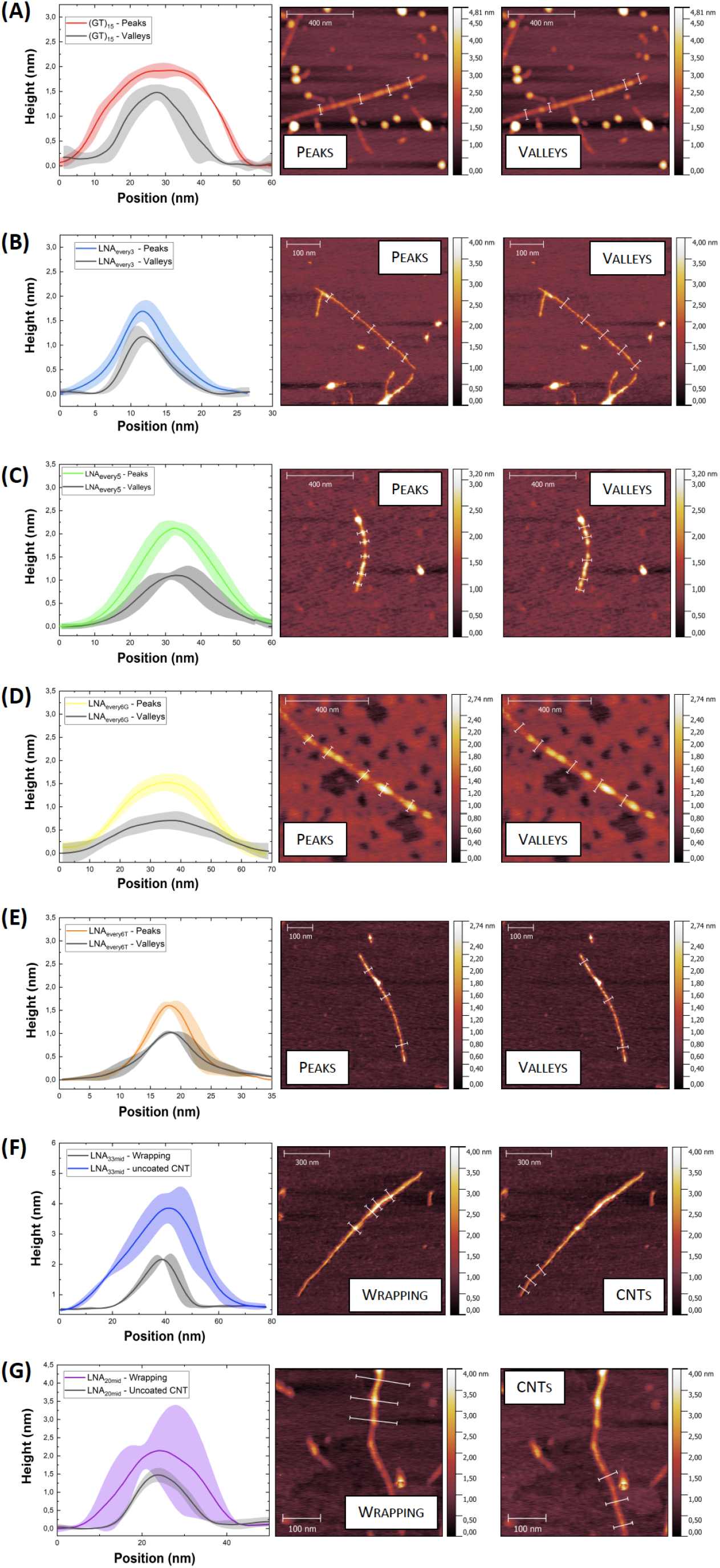
AFM characterization was performed to examine differences in the wrapping behavior of **(A)** (GT)_15_, **(B)** LNA_every3_, **(C)** LNA_every5_, **(D)** LNA_every6G_, **(E)** LNA_every6T_, **(F)** LNA_33mid_, and **(G)** LNA_20mid_. Average height profiles were extracted from coated (peaks) and uncoated (valleys) cross sections along the nanotube. A minimum of three positions were selected along the nanotube to extract the height profiles (indicated by the white lines in AFM height images). Solid lines in the height profiles represent the average height and shaded regions represent 1σ standard deviation (n = 3 – 5 profiles). Due to the presence of an additional layer of material on the substrate of the LNA_every6G_-SWCNTs, the heights estimated by the cross-section analysis shown are slightly underestimated. Additional measurements, taken across the nanotubes into exposed parts of the substrate show the true height of the nanotubes, which is used for comparison to the other wrappings **Figure SI.10**

The (GT)_15_ exhibited an overall regularized wrapping along the nanotube length (**Figure 3 (A)**). This consistency is in agreement with previous reports that have suggested helical wrapping of DNA onto the SWCNT surface^25,26^. Similarly, all LNA sequences with periodic lockings (LNA_every5_, LNA_every6T_, LNA_every6G_, and LNA_every3_) displayed regular wrapping heights along the surface of the nanotube (**Figure 3 (B – E)**). In addition, we note that the SWCNTs suspended using the periodic LNA sequences were smaller in height (LNA_every3_ – 1.17 ± 0.12 nm, LNA_every5_ – 1.09 ± 0.09 nm, LNA_every6T_ – 1.02 ± 0.25 nm, LNA_every6G_ – 0.82 ± 0.22 nm) compared to (GT)_15_-SWCNTs (1.48 ± 0.16 nm). Conversely, SWCNTs suspended using LNA sequences with blocks of locked bases (**Figure 3 (F – G)**) appeared to preferentially suspend larger diameter SWCNTs (LNA_33mid_ – 2.16 ± 0.03 nm, LNA_20mid_ – 1.47 ± 0.20 nm).

We attribute these differences to the relative increase of certain chiralities (such as the (6,5) for periodic lockings and the (10,2) or (9,4) for blocks of locked bases) in the LNA suspensions, in agreement with our observations from the PLE maps and absorbance spectra (**Figure 2**).

Examining the regularity of the wrapping for the different sequences, we observed a distinct difference between periodically locked sequences and the sequences with a block of continuous locking. Whereas the former showed frequent peaks regularly dispersed along the length of the nanotube, similar to (GT)_15_, LNA_33mid_ and LNA_20mid_ exhibited a more sporadic coverage. Rather than periodic peaks, LNA_33mid_ showed longer coated sections along the nanotubes. In addition, there were larger spacings between wrapped regions (**Figure SI.8**). Similar trends were observed for the LNA_20mid_ (**Figure SI.9**).

Subsequent analysis of the coated regions of LNA_33mid_ qualitatively demonstrated the consistency of the wrapping over these sections (**Figure SI.8, SI.9**). In addition, we observed that the wrapping heights of LNA with blocks of locked bases were much greater than both the periodically locked sequences and (GT)_15_. We attribute the distinct wrapping pattern of LNA_33mid_ and LNA_20mid_ to the bundled locking of bases in the middle of the sequence. The presence of locked bases limits the ability of this sequence to reconfigure on the nanotube surface, which may, in turn, limit the ability of these sequences to tightly wrap around the nanotube resulting in the higher wrapping heights observed.

While the free rotation of natural DNA enables a majority of bases in a wrapping sequence to π-stack onto the nanotube^25,26^, steric restrictions of LNA sequences at locked positions into the 3’-endo north conformation (**Figure SI.11**) may result in only a fraction of these bases being able to stack onto the SWCNT surface. Other bases may either self-stack onto each other or protrude from the nanotube surface disrupting any helical wrapping^26,27^ and lead to differences in the wrapping heights. The variability in wrapping behavior can also affect the amount of DNA/LNA that can stack onto the nanotube surface per unit area.

To further investigate differences in the wrapping behavior, we characterized all SWCNT solutions using *ζ*-potential (ZP) (**Figure 4 (A)**), which measures a nanoparticle’s surface charge. Due to a lack of salt in the DNA- and LNA-SWCNT solutions, we assume that only a single layer of DNA or LNA can adsorb onto the nanotube surface as the electrostatic repulsion between the phosphate groups along the backbone should prevent multilayer adsorption. Hence, we can use ZP values as a proxy for the relative amount of negatively charged oligonucleotides adsorbed onto the surface^26,29^. The ZP measurements showed only slight variations for the majority of LNA-SWCNTs compared to (GT)_15_-SWCNTs. Two exceptions were LNA_33mid_, which showed a notably lower ZP, and LNA_every3_, which showed the most negative ZP of all complexes. Both of these sequences had the same percentage of locking, however as was shown in the AFM images (**Figure 3, SI.7, SI.8**) due to differences in the locking periodicity, these sequences adopted very different wrapping structures on the nanotube surface. While LNA_33mid_ had certain areas with a dense coverage, a majority of the nanotube surface remained uncoated (indicated by the lower height profile extracted from the AFM images). We hypothesize that these areas of uncoated SWCNT reduce the overall charge density of the complex, resulting in a lower than expected ZP value. Conversely, LNA_every3_ had a very dense wrapping, with almost the entirety of the nanotube surface covered by frequent and tightly packed peaks, which could result in a higher charge density and hence the more negative ZP value observed.

**Figure 4:**
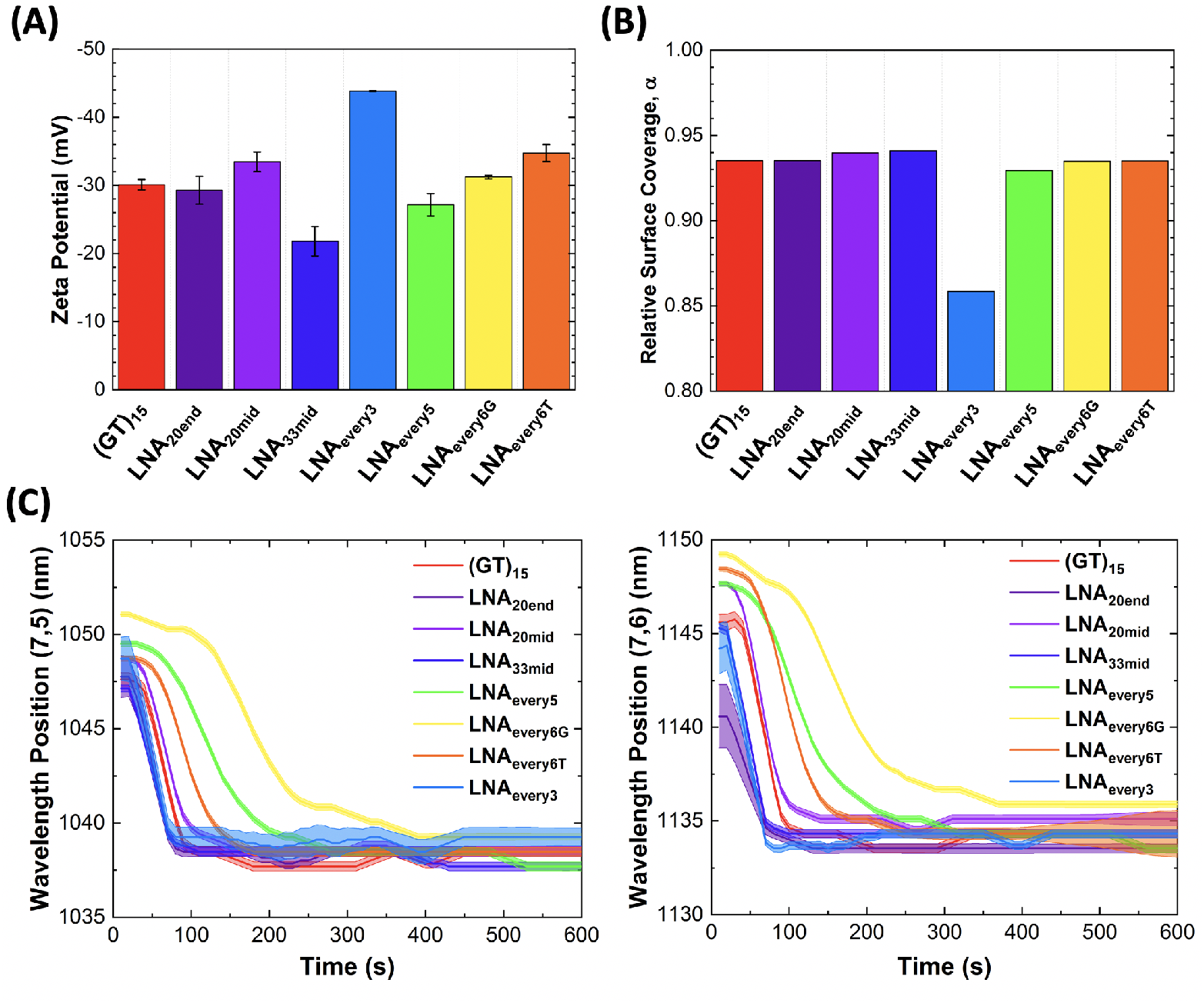
Wrapping behavior of DNA and LNA sequences. **(A)** *ζ*-potential of SWCNTs functionalized with (GT)_15_ and LNA analogues in DI water. Measurements were performed in triplicates with error bars representing 1σ standard deviation, as calculated from individual data shown in **Table SI.2**. **(B)** The relative surface coverage of the different DNA and LNA polymer wrappings calculated relative to N-Methyl-2-pyrrolidone (NMP)^28^. **(C)** Modulation of the fluorescence emission wavelength position of the (7,6) and (7,5) chirality peaks of all sequences parameters as a function of time following SDOC addition. Shaded regions represent the standard deviation (3σ) of the peak fit used at each time point.

In order to further probe differences in surface coverage between the wrappings, the relative surface coverage, α, was estimated based on the effective dielectric constant, *ϵ_eff_*^28,30^ (**Figure 4 (B)**)

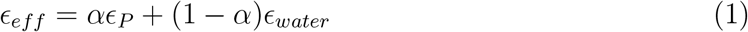

where *ϵ*_water_ and *ϵ*_P_ are the contributions to the dielectric constant from the surrounding water and wrapping polymer, respectively. For both LNA and DNA sequences, *ϵ*_P_=4 was used^31^. *ϵ*_eff_ was obtained from the slope of the linear fit of the solvatochromic shift, (*E*_11_)^2^Δ*E*_11_, versus the diameter of the nanotube to the power of negative 4, *d*^-4^, for the various chiralities present in each suspension (**Figure SI.12**). The experimental solvatochromic shifts were determined by extracting the peak positions for the different chiralities in the PLE plots obtained (**Figure 2**). All data used to calculate the values of *ϵ*_eff_ and α are included in **Table S2** and energy shift values are included in **Table S3**. All LNA sequences had similar calculated surface coverage values to nanotubes suspended using (GT)_15_ with the exception of LNA_every3_ (**Figure 4 (B)**). As this approach considers all nanotube chiralities to approximate an average surface coverage independent of chirality, it is not capable of capturing chirality specific effects. This is particularly problematic for the LNA_every3_, which exhibits a strong preference towards the (6,5) chirality. We note that while the energy shift values were the same for LNA_every3_ and all other sequences when examining the (6,5) chirality, large deviations were observed for the larger diameter nanotubes, such as the (10,2), suspended using the LNA_every3_ (**Table S3**). Furthermore, as a result of the selectivity of LNA_every3_ towards the (6,5) chirality, the fluorescence intensity of the other chirality peaks was significantly lower. This in turn leads to a poorer fit of the data (indicated by the lower Adj. R^2^ value **Table S2**), which can result in a significant underestimation of α. More generally, we believe that this method of approximating surface coverage can lead to discrepancies if strong chirality preference is displayed by the different wrappings. However, it does provide information on the relative accessibility of the nanotube surface to water by comparing the experimentally determined solvatochromic shifts for a given chirality (**Table S3**).

The effective binding affinities of the different wrappings were characterized by examining surfactant displacements kinetics, as done previously^24^. In this method, surfactants (such as SDOC) added to the solution competitively bind to the nanotube surface and displace the DNA wrapping. This displacement results in blue-shifting of the fluorescence peak positions at a rate that depends on the DNA binding affinity^24^. The relative binding affinities of the various oligonucleotide wrappings can therefore be inferred by continuously monitoring the positions of the (7,6) and (7,5) fluorescence emission peaks following SDOC addition (final concentration 0.1%). As shown in **Figure 4 (C)** and **SI.6**, the LNA_every6T_, LNA_every6G_, and LNA_every5_ showed slower displacement kinetics compared to (GT)_15_ for both the (7,5) and (7,6) peaks, implying stronger binding affinity for these engineered wrappings to both of these chiralities.

We further examined the fluorescence stability of the different wrappings in the presence of CaCl_2_ (**Figure 5, SI.13, SI,14**). Previous studies have shown that the fluorescence of (GT)_15_-SWCNT sensors undergoes changes in the presence of varying cation concentrations^19^. In agreement with these observations, we observe both quenching (−22.6 ± 2.7% and −12.4 ± 2.7%) and wavelength red-shifting (1.6 ± 0.0 nm and 2.3 ± 0.2 nm) for the (7,5) and (7,6) (GT)_15_-SWCNT chirality peaks following the addition of CaCl_2_ (**Figure 5 (A)**). Similarly, certain LNA-SWCNTs, including LNA_20end_, LNA_33mid_, and LNA_every3_, exhibited large fluorescence changes upon addition of CaCl_2_ (**Figure SI.13, SI.14**). We attribute this to an instability of these wrappings on the (7,5) and (7,6) chiralities, in line with our observations from the SDOC replacement experiments (**Figure 4 (C)**).

**Figure 5:**
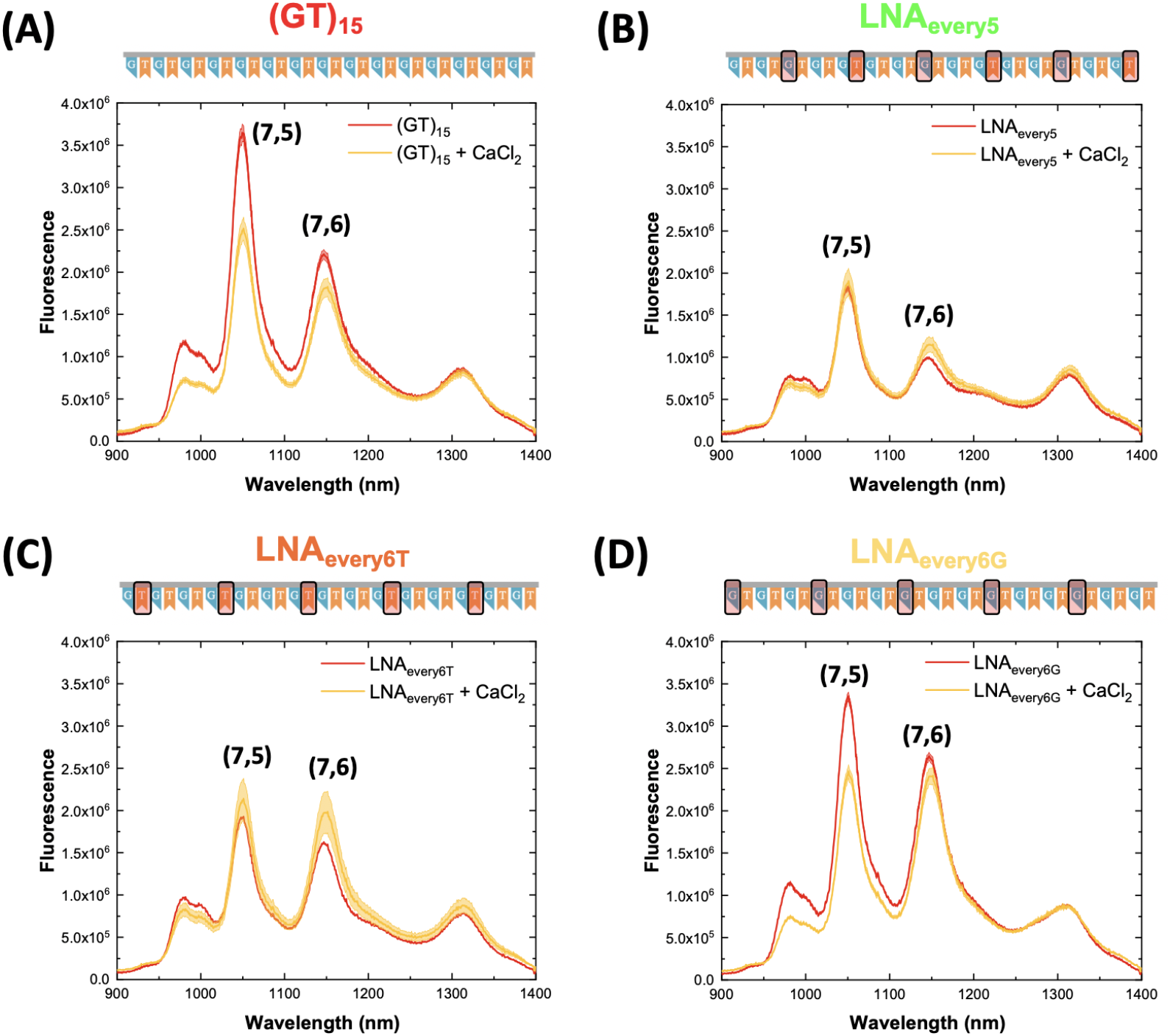
Response of DNA- and LNA-sensors to CaCl_2_. Representative spectra of the response of **(i)** (GT)_15_, **(B)** LNA_every5_, **(C)** LNA_every6T_, and **(D)** LNA_every6G_ sensors following the addition of 0.5 M CaCl_2_ (final concentration: 5 mM, excitation: 660 nm). Graphs include the spectra before (**red**) and after addition of CaCl_2_ (**yellow**). The solid line represents the average wavelength shift with the shaded regions representing 1σ standard deviation (n = 3 technical replicates). All samples were incubated for 30 min. Spectra for all other LNA sequences are included in **Figure SI.13**.

Conversely, LNA-SWCNT complexes with more spaced locking demonstrated superior stability in the presence of the divalent cations. For example, the LNA_every5_-SWCNT (**Figure 5 (B)**) showed only minor changes in the fluorescence intensity of both the (7,5) or (7,6) peaks (−6.5 ± 1.0% and +4.6 ± 1.0%, respectively) and the shifting of the (7,5) was reduced by approximately 44% (**Figure SI.14**). Interestingly, despite the same periodicity of locking for both the LNA_every6T_ (**Figure 5 (C)**) and LNA_every6G_ (**Figure 5 (D)**) sequences, we observed pronounced differences in the behavior of these sensors. Whereas the LNA_every6T_-SWCNT exhibited only a minor increase in the fluorescence intensity of the (7,5) chirality (+0.3 ± 3.2%), the LNA_every6G_-SWCNT had pronounced quenching (−24.5 ± 3.4%). This suggests that the base itself, and not only the periodicity of locking, can greatly impact the stability of the LNA-SWCNT sensors. Moreover, the differences in the response of the (7,5) and (7,6) peaks implies that the fluorescence changes are dependent on the interaction of a particular LNA sequence with a specific chirality and not simply a direct function of the percentage locking. This observed chirality specificity further supports our previous observations of different fluorescence behaviors for certain LNA sequences in the PLE plots (**Figure 2 (A)**). Chirality specific responses are also observed for the LNA_20mid_-SWCNT (**Figure SI.13**), where the changes in intensity and wavelength position of the (7,5) chirality (+19.3 ± 16.3% and 0.9 ± 0.2 nm) were much lower than for the (7,6) peak (+55.6 ± 20.4% and 3.2 ± 0.2 nm).

Current dopamine sensors based on (GT)_x_-SWCNTs are ‘turn-on’ sensors, exhibiting strong relative fluorescence intensity changes on recognition of the neurotransmitter^8,9^. As a result, any competing changes in the fluorescence intensity could result in inaccurate concentration measurements. Given the variability of Ca^2+^ concentrations in different biofluids and the fluctuations of Ca^2+^ involved in neurotransmission, large intensity responses of SWCNT-based sensors to this cation could be detrimental to their applicability *in vitro* and *in vivo*. At high Ca^2+^ concentrations, we show that the (GT)_15_-SWCNTs exhibit the largest intensity decrease for both the (7,5) and (7,6) chiralities. Furthermore, as we have shown in previous work^19^, absolute cation concentration is more important than the nature of the individual cations. Therefore, in ionically complex solutions the (GT)_15_-SWCNT would lead to a significant underestimation of dopamine concentration. Conversely, the LNA_33mid_-, LNA_20mid_-, and LNA_every3_-SWCNTs showed large intensity increases after the addition of CaCl_2_, which would lead to overestimation of the dopamine concentration. To overcome the problems associated with significant underestimation or overestimation, an ideal sensor would exhibit minimal intensity changes, if any, in the presence of the Ca^2+^.

Additional measurements were carried out across a wider range of CaCl_2_ concentrations (0.05 mM – 50 mM) to further probe the stability of these sequences. Spectra were collected following an incubation period of 30 min and peak wavelength shifts and intensity changes were calculated for the predominant (7,5) and (7,6) chiralities (**Figure SI.14**). Concentration-dependent wavelength red-shifting was observed for the (7,5) and (7,6) chiralities for all sequences, with maximum recorded shifts exceeding 6 nm for the LNA_33mid_. At the highest Ca^2+^ concentration tested, five of the LNA sequences (LNA_20end_, LNA_20mid_, LNA_every6G_, LNA_every5_, and LNA_every6T_) showed lower shifting than (GT)_15_-SWCNTs for the (7,5) chirality peak. Moreover, compared to (GT)_15_, over one-order-of-magnitude higher concentration of Ca^2+^ was required to induce any wavelength response for the (7,5) peak for four of these sequences (LNA_20end_, LNA_20mid_, LNA_every6G_, and LNA_every6T_). Although the magnitude of shifting for the (7,6) peak compared to (7,5) peak was greater for all sensors at higher Ca^2+^ concentrations, several LNA-SWCNTs again exhibited superior stability towards cation-induced wavelength changes, indicative of greater LNA conformational stability^19^. For example, the magnitude of shifting for the LNA_every6T_-SWCNTs (1.8 ± 0.5 nm) was 38% lower than for the (GT)_15_-SWCNTs (3.0 ± 0.3 nm). It is also worth noting that at the highest Ca^2+^ concentration, the red-shift of the LNA_20mid_ and LNA_20end_ sensors was 4.4 ± 0.7 nm and 2.3 ± 0.2 nm, respectively, compared to the 2.9 ± 0.3 nm shift of the (GT)_15_, despite their shifting response being significantly lower than the (GT)_15_-SWCNT for the (7,5) peak. This large disparity in the (7,5) and (7,6) shifting response again highlights the chirality specific behavior of the different LNA sequences.

These observations further emphasize the importance of considering locking position, in addition to total locking percentage, when designing LNA-SWCNT sensors. Furthermore, these results prove that for LNA, increasing the percentage of locked bases does not always equate to a more stable LNA sensor as is clearly shown by comparing the LNA_every6T_ (16.6% locked) sensor to LNA_33mid_ and LNA_every3_ (33.3% locked). These findings enabled us to identify several LNA-SWCNTs that exhibit superior signal stability in varying Ca^2+^ concentrations compared to (GT)_15_-SWCNTs. Amongst the sequences studied, LNA_every6T_- and LNA_every5_-SWCNTs were found to provide the greatest stability against undesirable intensity changes across the range of calcium concentrations for both the (7,5) and (7,6) peaks. At high Ca^2+^ concentrations, the LNA_every6G_- and LNA_20end_-SWCNTs also demonstrated improved stability compared to the (GT)_15_-SWCNT complex (**Figure SI.14**).

The increased resistance of certain LNA sequences towards ion-induced fluorescence changes has been attributed to the increased conformational rigidity of these sequences ^19^. Owing to this, it was uncertain whether these sensors would retain the ability to interact with other bioanalytes, especially those such as dopamine which are thought to modulate DNA-SWCNT fluorescence by also inducing DNA conformational changes^9,19^. In order to determine the validity of this concern, we examined the response of our DNA- and LNA-SWCNT sensors to a range of dopamine concentrations in the absence of CaCl_2_ (**Figure 6, SI.15, SI.16**). Concentration-dependent intensity changes were seen across multiple SWCNT chiralities (**Figure 6**), with pronounced increases observed for both the (7,5) and (7,6) chiralities. Although some LNA-SWCNTs have reduced turn-on responses compared to (GT)_15_-SWCNTs, all LNA sensors retained their ability to interact with dopamine (**Figure SI.15**) despite the changes in their sensitivity to Ca^2+^. As shown in **Figure 6** (and **Figure SI.17**), the limit of detection for the LNA sensors matched the (GT)_15_-SWCNTs at 10 nM dopamine. We also note that the turn-on response of LNA_every3_-SWCNTs was significantly greater than for (GT)_15_-SWCNTs for the (7,5) peak (15.5× versus 7.7×). Similarly, the LNA_every5_-SWCNTs had comparable responses to the (GT)_15_-SWCNTs for the (7,5) peak and marginally higher turn-on responses for the (7,6) peak, especially at intermediate dopamine concentrations (7.7× versus 5.7× in 1 *μ*M dopamine and 16.0× versus 14.7× in 10 μM dopamine).

**Figure 6:**
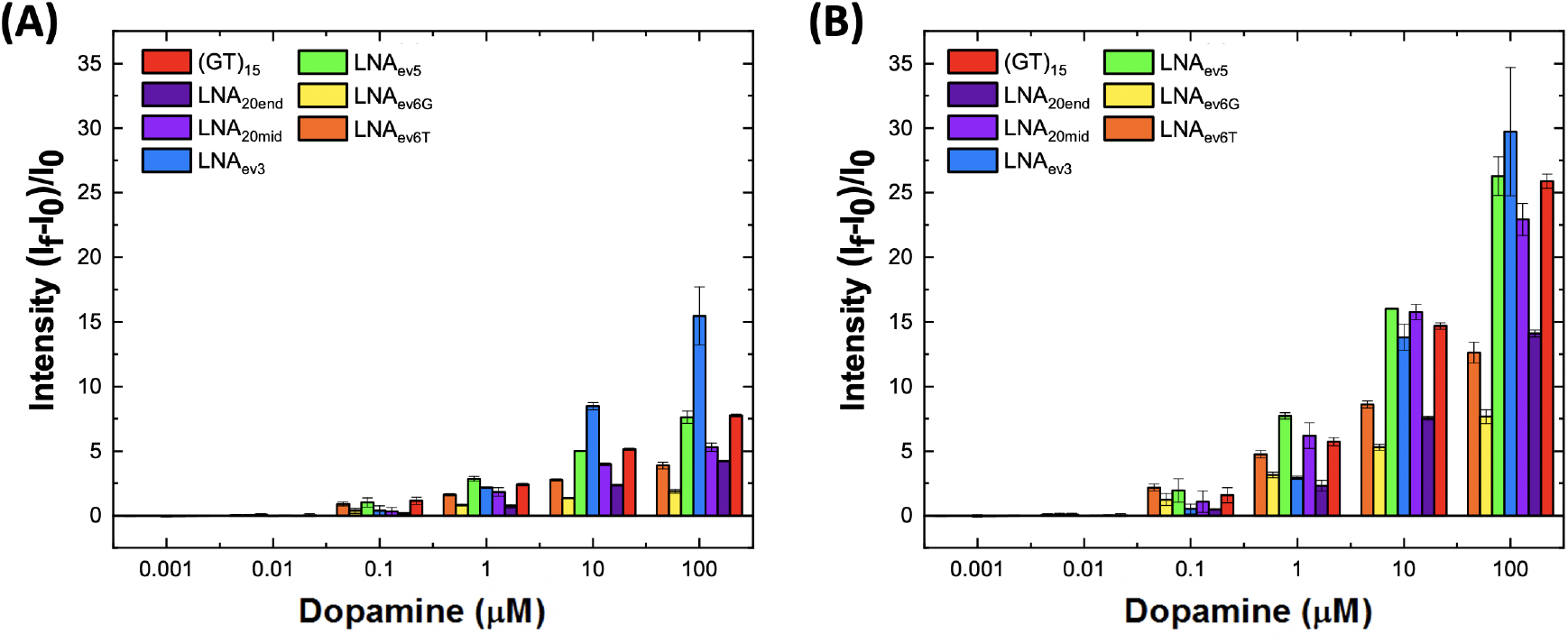
Concentration-dependent intensity response of DNA- and LNA-SWCNTs to dopamine in the absence of CaCl_2_ (excitation: 660 nm). Relative intensity change of the **(A)** (7,5) peak and **(B)** (7,6) peak following the addition of Dopamine. All samples were incubated for 30 min. Error bars represent 1σ standard deviation (n = 3 technical replicates). Both graphs are plotted on the same axis scales for comparison.

The resilience of LNA-sensors to salt-induced fluorescence perturbations^19^, coupled with their retained ability to interact with dopamine, can thus provide a general basis for designing sensors with improved diagnostic capabilities. To this end, we tested the ability of the DNA- and LNA-SWCNTs to detect dopamine (100 μM) in the presence of CaCl_2_ (5 mM) (**Figure 7, Figure SI.18**). Sensors were first incubated with CaCl_2_ and spectra were recorded after 30 min. Freshly prepared dopamine solution was subsequently added to the sensor solution and spectra were again acquired following an additional incubation period.

**Figure 7:**
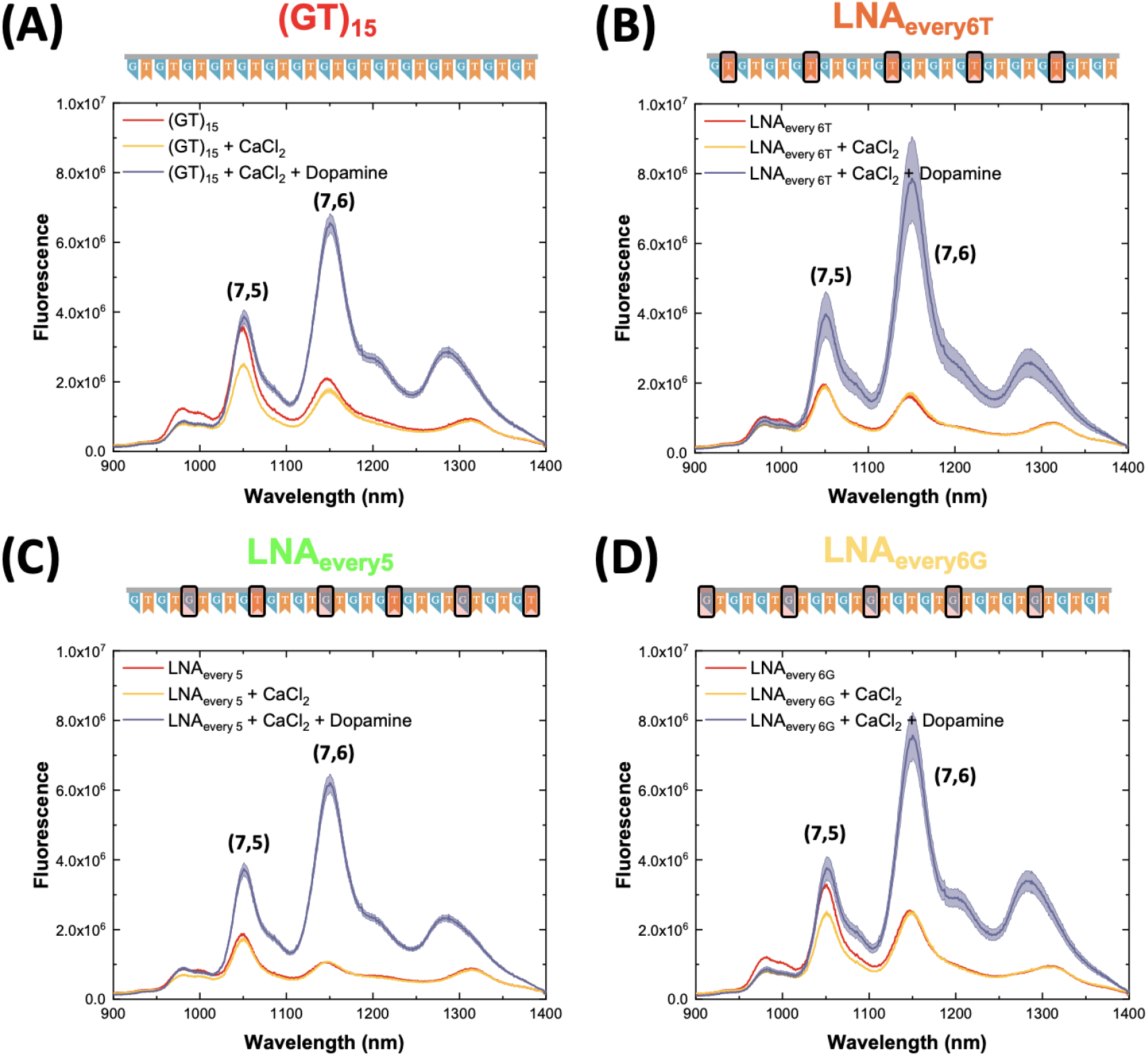
Deconvoluting the response of LNA sensors to dopamine and CaCl_2_. **(A-D)** Spectral response of (GT)_15_ and representative LNA sensors following the addition of 0.5 M CaCl_2_ (final concentration: 5 mM) and 10 mM dopamine (final concentration: 100 *μ*M, excitation: 660 nm). Graphs include the spectra before addition (**red**), following the addition of CaCl_2_ (**yellow**) and following subsequent addition of dopamine (**purple**). All samples were incubated for 30 min post-addition. The solid line represents the average wavelength shift with the shaded regions representing 1σ standard deviation (n = 3 technical replicates).

In line with our previous observations, following the addition of CaCl_2_ the fluorescence intensity of the (GT)_15_-SWCNTs decreased for both the (7,5) and (7,6) chiralities. As expected, the addition of dopamine increased the fluorescence of the sensor, however, due to the lower starting fluorescence in the presence of Ca^2+^, the (7,5) peak only marginally increased versus the initial spectrum (**Figure 7 (A)**).

The undesirable fluorescence modulation was further exacerbated at higher Ca^2+^ concentrations for the (GT)_15_-SWCNTs, to the point where in the presence of 50 mM CaCl_2_ the (7,5) peak in the final spectrum (post dopamine addition) even appeared to quench compared to the original intensity (**Figure SI.19**). Contrary to this, the LNA_every6T_- (**Figure 7 (B)**), LNA_every5_- (**Figure 7 (C)**), and LNA_20mid_-SWCNTs (**Figure SI.18**) showed only minor changes in the fluorescence intensity of both the (7,5) and (7,6) peaks in the presence of 5 mM Ca^2+^. Furthermore, the presence of the cations did not interfere with the ability of these sensors to interact with dopamine, with significant turn-on responses observed for each of these LNA-SWCNTs.

Although the fluorescence of the LNA_every6G_-SWCNT (7,5) peak quenched following the addition of Ca^2+^, the (7,6) peak intensity was unchanged within error. In line with the observations for (GT)_15_-SWCNTs, the addition of dopamine recovered the fluorescence of the (7,5) peak, although final intensity increases were marginal versus the original spectrum. Despite not reacting to the Ca^2+^ cations, the (7,6) peak fluorescence intensity was significantly enhanced by the addition of dopamine. In fact, the final intensity of the LNA_every6G_-SWCNTs was higher (normalized to sensor concentration) than the (GT)_15_-SWCNTs, implying that these sensors would additionally benefit from increased penetration depth in biological samples^12,32^. Furthermore, the distinct responses of the (7,5) and (7,6) peaks to CaCl_2_ and dopamine enabled simultaneous multi-modal sensing of these two analytes. By exciting the sensors at 660 nm and monitoring the fluorescence intensity of the two peaks over time, information on the concentration of both calcium (via the (7,5) peak) and dopamine (via the (7,6) peak) could be inferred. This is, to the best of our knowledge, the first optical sensor that can simultaneously detect calcium and dopamine, opening the door to a plethora of new and improved biosensing opportunities.

Recent work by Pinals *et al.* demonstrated that protein adsorption onto the nanotube surface when sensors are placed in biologically complex solutions can attenuate the optical response of (GT)_6_-SWCNT sensors to dopamine^33^. This decrease was attributed to protein adsorption on the nanotube surface, which resulted in both an increase of the baseline fluorescence and decrease of the final fluorescence intensity post-dopamine addition. Given these findings, we sought to investigate whether prolonged incubation in complex biological media would affect the ability of the LNA-SWCNTs to detect dopamine (**Figure 8**).

**Figure 8:**
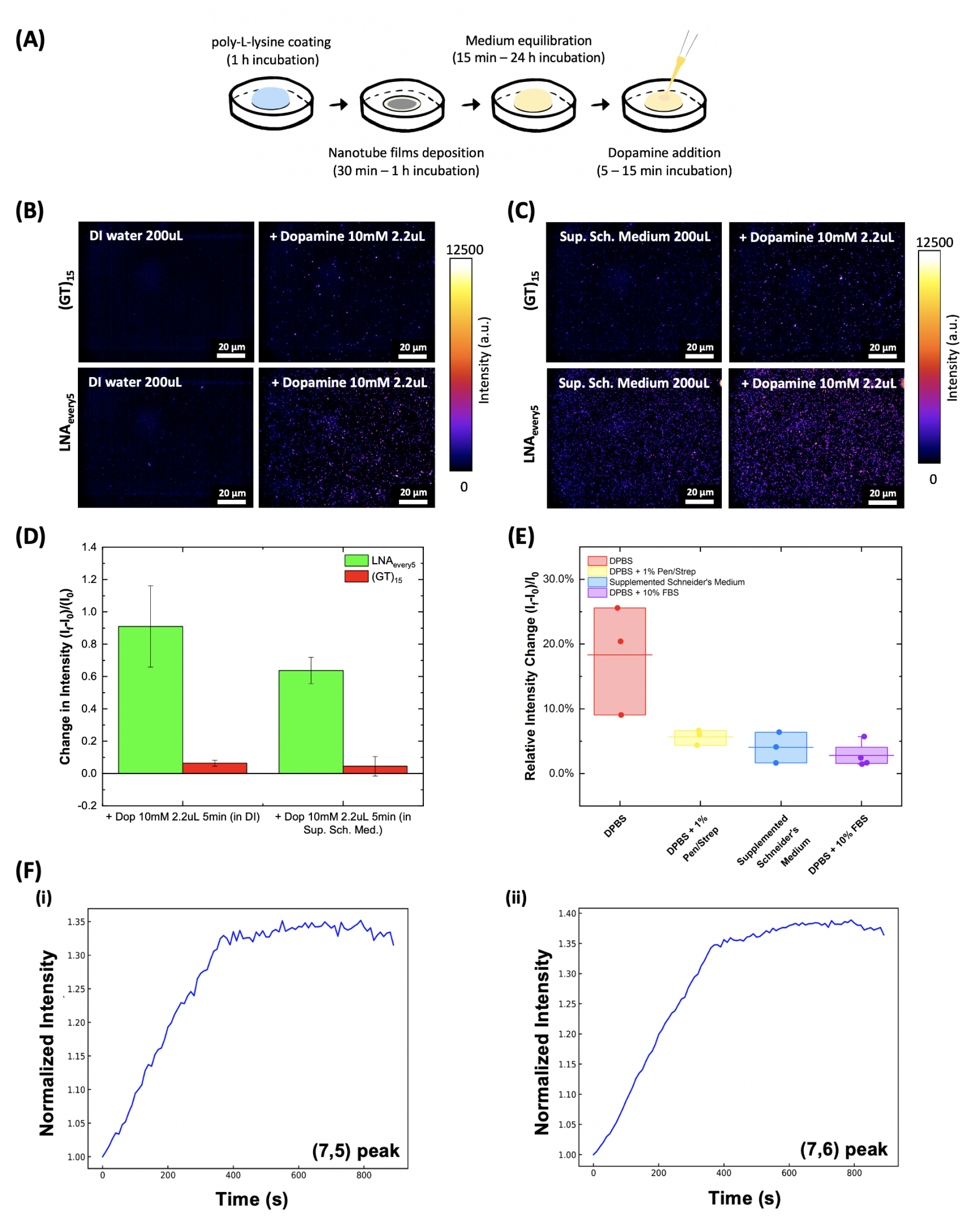
Comparison of DNA- and LNA-SWCNT sensors for dopamine sensing in complex media conditions. **(A)** Schematic detailing the experimental procedure used to prepare nanotube films to study the effect of medium composition on nanotube fluorescence. All nanotube films were incubated in medium for either 15 min or 24 h prior to measurement. **(B)** nIR fluorescence images of sensors that were immobilized onto a glass surface and incubated in DI water before (**left**) and after (**right**) addition of dopamine (final concentration 100 *μ*M). **(C)** nIR fluorescence images of sensors that were immobilized onto a glass surface and incubated in supplemented Schneider’s medium (10% FBS, 1% Pen/Strep) before (**left**) and after (**right**) addition of dopamine (final concentration 100 μM). **(D)** Comparison of the intensity increase of the sensors following addition of dopamine (100 μM) in DI water and supplemented Schneider’s medium. Intensity changes were calculated from the mean intensity of the wide-field images obtained (**Figure SI.28 – SI.31**, excitation 780 nm, emission filter 980 nm LP) versus the mean intensity of the sensors in either DI water or supplemented Schneider’s medium. Error bars represent the standard deviation from three technical replicates. **(E)** Fluorescence intensity increase of the LNA_every5_ films incubated media with different composition following the addition of dopamine (final concentration: 100 μM). Intensity changes were calculated from the mean intensity of the wide-field images obtained (**Figure SI.32 – SI.35**). Central line represents the average turn-on response. Dots represent the turn-on response of individual replicates (n = 3 – 4 technical replicates). **(F)** Fluorescence intensity response of the **(i)** (7,5) and **(ii)** (7,6) chirality of LNA_every5_ sensors following the addition of exogenous of dopamine (final concentration: 100 μM) in a *Drosophila* neuronal cell culture. *Drosophila* neurons were extracted from the brains of L3 larvae and re-suspended post-digestion in supplemented Schneider’s medium. Neuronal cell cultures contained transgenic dopamine neurons induced to express CsChrimson (representing approximately 1% of the sample). Cells were grown for 24 h at room temperature in 96-well plates in the presence of SWCNT prior to measurement (excitation: 660 nm, exposure time: 10 s).

The dopamine sensing capability of surface immobilized LNA- and DNA-SWCNTs were compared in DI water and in a complex biological medium, supplemented Schneider’s medium (10% FBS, 1% Pen/Strep) using a custom-built NIR microscope^34^. Supplemented Schneider’s medium was chosen due to its widespread use in culturing *Drosophila* neurons^35^. (GT)_15_-SWCNTs and LNA_every5_-SWCNTs were used to prepare nanotube films (**Figure 8 (A)**, as described in Materials and Methods) by incubating equal concentrations of nanotubes in DI water on poly-L-lysine coated petri dishes. Nanotube films were covered with 200 μL of either DI water or supplemented Schneider’s medium, and left to incubate for 5 mins prior to acquiring any images. Both (GT)_15_- and LNA_every5_-SWCNTs had lower starting fluorescence in DI water than in the complex medium in agreement with observations by Pinals *et al.*. Moreover, in all preparations LNA_every5_-SWCNTs produced denser films leading to higher fluorescence compared to (GT)_15_ (**Figure 8 (B – C), SI.28 – SI.31**) under the 780 nm excitation. This was in part attributed to differences in the surface binding affinity of the LNA- and DNA-SWCNTs^25^, but also to differences in the relative fluorescence of these sensors at 780 nm excitation (**Figure 2 (A)**).

The addition of dopamine resulted in an intensity increase of the sensor films in both DI water and supplemented Schneider’s medium. The relative intensity increase (**Figure 8 (D)**) normalised to the initial intensity was determined using the mean intensity of the wide-field images collected (**Figure SI.28 – SI.31**). Although both sensors possessed the ability to interact with dopamine in DI water, the turn-on response of the LNA_every5_-SWCNTs (+90.9 ± 25.2%) was significantly higher than for (GT)_15_-SWCNTs (+6.4 ± 1.9%). In supplemented Schneider’s medium, the ability of (GT)_15_-SWCNTs to detect dopamine via intensity changes decreased, and no significant increase in intensity of the film was detected following the dopamine addition (+4.5 ± 6.0%). While the extent of the turn-on response also reduced for LNA_every5_-SWCNTs, a substantial increase was still recorded (+63.7 ± 8.1%) following the addition of dopamine. Furthermore, the overall intensity of the LNA_every5_-SWCNT films both pre- and post-dopamine was greater in supplemented Schneider’s medium than in DI water.

We further examined the impact of longer incubation times (> 24 h) using three different media compositions and DPBS as a control (**Figure SI.32 – SI.37**). DPBS, a pH balanced salt solution with no Ca^2+^ or Mg^2+^ ions, was used as a negative control to account for changes in the fluorescence properties of nanotube films stored at room temperature for extended periods of time. By comparing the fluorescence turn-on response following dopamine addition for films incubated in DPBS to those incubated in a more complex medium, we could examine the impact of additives such as proteins (FBS) or antibiotics (Pen/Strep).

LNA_every5_-SWCNT films were prepared and incubated in medium for 24 h at room temperature. All sensor films retained an ability to interact with dopamine, however we note that the fluorescence turn-on response was significantly reduced for films incubated in a more complex medium compared to those incubated in DPBS (**Figure 8 (E)**). In addition, we observed a significant decrease in the mean fluorescence intensity of films (prior to dopamine addition) incubated in any medium containing FBS (**Figure SI.32 – SI.37**), although this did not significantly impact the ability of these sensors to interact with dopamine compared to sensors incubated with only antibiotics. Despite this relative decrease, by increasing the density of the nanotube sensor films, we could improve the average turn-on response of the sensors incubated in supplemented Schneider’s medium following dopamine addition from +4.0 ± 2.4% to +7.9 ± 0.3% (**Figure SI.36**). Further increases in film density enabled even greater enhancement, with the ultrahigh density films incubated in 10% FBS exhibiting a dopamine turn-on response of +61.7 ± 17.8% (**Figure SI.37**). These findings suggest that although extended incubation periods can impact the ability of the nanotube sensor films to interact with dopamine, the extent by which reactivity is reduced can be tuned by adjusting the sensor density^9^.

Finally, we tested the sensor response of the improved LNA_every5_-SWCNT sensors in the presence of Drosophila larvae neurons. Drosophila neurons were extracted from the brains of L3 larvae and re-suspended post-digestion in supplemented Schneider’s medium. Neurons were incubated in LNA_every5_-SWCNT covered, poly-L-lysine coated 96-well plates overnight, and additional LNA_every5_ sensors were mixed into the well immediately prior to measurement. Kinetic measurements were collected following the addition of exogenous dopamine (final concentration: 100 μM) for both the (7,5) and (7,6) chirality (**Figure 8 (F)**). Significant turn-on responses were recorded for both peaks with final intensity increases of 31.5% and 36.4% for the (7,5) and (7,6) chirality peaks, respectively. This indicates that our sensors remain capable of detecting dopamine even in the presence of cells and suggests that these sensors could be used for *in vitro* sensing applications.

## Conclusion

SWCNTs provide a platform that can overcome many of the restrictions currently facing biosensing technologies as they enable real-time continuous monitoring^11,36^ and provide a means of increasing spatiotemporal resolution^9^. Research into DNA-SWCNTs has thus far largely focused on expanding biochemical detection capabilities, and many sensors are now capable of long-term imaging for a wide variety of analytes^8,30,37–39^. Despite this every growing library of DNA-SWCNT sensors, little work has been done to optimise existing sensors for *in vitro* and *in vivo* conditions.

In this work, we took the (GT)_15_-SWCNT sensors, known for their ability to detect dopamine^8,10,40^ and systematically introduced locked bases along the sequence in order to examine the effect of *‘locking’* on their sensing behaviour. Initial characterization of the sensors showed that both locking periodicity and percentage affects the relative affinity of the sequences towards certain chiralities. Further experiments sought to examine the differences between the optical response of the sensors to increasing concentrations of Ca^2+^ cations^19^. Optical fluctuations caused by the presence of calcium may lead to inaccuracies in concentration readouts and, owing the central role of Ca^2+^ cations in the neurotransmission^23^, could limit the use of these sensors for *in vitro* and *in vivo* applications. By incorporating certain LNA bases we improved the stability of (GT)_15_-SWCNTs towards undesirable fluorescence modulation in the presence of Ca^2+^. This stability appeared to depend strongly on the nature of base that is locked, with no systematic dependence observed to the percentage of locking. Furthermore, we observed a strong chirality specificity where depending on the periodicity of locking and nature of bases (G or T) locked, only certain nanotube chiralities exhibited improved resilience towards salt induced fluorescence changes.

In addition to studying the optical behaviour of these sensors in the presence of Ca^2+^, we also probed their response to dopamine. The presence of locked bases did not destroy the ability of the sensors to interact with dopamine nor did it negatively impact the limit of detection. In fact, for LNA_every3_-SWCNTs and LNA_every5_-SWCNTs, an improved turn-on response rate versus (GT)_15_-SWCNTs was obtained for the (7,5) and (7,6) chiralities, respectively. Moreover, not only did the LNA sensors show superior sensing capabilities to dopamine in the presence of Ca^2+^ cations, but the nanobioengineered hybrid LNA_every6G_-SWCNT also provided an unprecedented sensing capability to monitor both Ca^2+^ and dopamine simultaneously. This is, to the best of our knowledge, the first optical sensor that is capable of detecting these two analytes concurrently, a differentiation that could be used to push forward our understanding of the intricate mechanisms of neuronal signalling.

Furthermore, we demonstrated that our LNA sensors retain their superior sensing capabilities in more complex media, such as supplemented Schneider’s medium, where high levels of proteins and other small molecule species are present. This ability enabled us demonstrate the reversibility of the LNA_every5_-SWCNTs to exogenous dopamine in the presence of *Drosophila* neurons.

Based on these findings, we believe that this methodology can be used to generate a new class of SWCNT sensors. The use of LNA provides both a solution for the limitations currently faced by DNA-SWCNTs in ionically complex solutions and enables more rational design of SWCNT based sensors for a wider range of biological applications and conditions. Furthermore, it does not reduce the biocompatibility or sequence space offered by DNA-SWCNTs. Specifically, the ability of our sensors to identify and measure dopamine with higher selectivity provides powerful tool for use in the clinical diagnostics of diseases such as Parkinson’s Disease.

## Acknowledgement

The authors are thankful for funding support from the Swiss National Science Foundation (SNSF) Assistant Professor (AP) Energy Grant. The authors also thank Prof. Kevin Sivula from the Laboratory for Molecular Engineering of Optoelectronic Nanomaterials (LIMNO) at EPFL for granting access to the AFM facility. DM is funded by a Marie Skłodowska Curie postdoctoral fellowship under the Eurotech Postdoc Programme. The Eurotech Postdoc Programme is co-funded by the European Commission under its framework programme Horizon 2020. Grant Agreement number 754462.

This document was prepared as an account of work sponsored by an agency of the United States government. Neither the United States government nor Lawrence Livermore National Security, LLC, nor any of their employees makes any warranty, expressed or implied, or assumes any legal liability or responsibility for the accuracy, completeness, or usefulness of any information, apparatus, product, or process disclosed, or represents that its use would not infringe privately owned rights. Reference herein to any specific commercial product, process, or service by trade name, trademark, manufacturer, or otherwise does not necessarily constitute or imply its endorsement, recommendation, or favoring by the United States government or Lawrence Livermore National Security, LLC. The views and opinions of authors expressed herein do not necessarily state or reflect those of the United States government or Lawrence Livermore National Security, LLC, and shall not be used for advertising or product endorsement purposes.

## Supplementary Information

Data supporting the findings of this study are available within the paper and its Supplementary Information file. All other relevant data and other findings of this study are available from the corresponding author upon reasonable request.

## Materials and Methods

### Materials

All chemicals and materials were purchased from Sigma-Aldrich, unless specified otherwise. Purified HiPco-SWCNTs were purchased from NanoIntegris (Lot. No. HP29-064). All ssDNA and ssLNA sequences were purchased from Microsynth.

### Preparation of SWCNT Solutions

Purified HiPco-SWCNTs (~0.5 mg) were suspended in 500 μL of either ssDNA or ssLNA solution (100 μM dissolved in DI water). The mixture was sonicated in an ice bath for 90 mins (power = 40W) using a Cup Horn sonicator (140 mm, Qsonica). Following sonication, the samples were centrifuged (Eppendorf Centrifuge 5424R) for 4 hours at 21,130 x g and 4 °C to remove SWCNT aggregates. The supernatant was collected and dialysed for 23 hours in 2 L of DI water (14,000 Da MWCO cellulose membrane). Post-dialysis, all samples were centrifuged again for 4 hours (21,130 x g and 4 °C) to remove any additional aggregates that may have formed.

Absorbance spectra of all samples were acquired using a UV/vis/nIR scanning spectrometer (Shimadzu 3600 Plus) with a quartz cuvette (Suprasil quartz, path length 3 mm, Hellma). The concentrations for all samples were calculated using an extinction coefficient Abs_632nm_ = 0.036 L mg^-1^ cm^-1^ and diluted to 10 mg/L for all measurements, unless otherwise stated. All DNA- and LNA-SWCNT solutions were stored at 4 ^°^C in order mitigate aggregation of the SWCNTs^38^.

### Fluorescence spectroscopy measurements

Fluorescence emission spectra were collected using a custom built near-infrared optical setup on an inverted Nikon Eclipse Ti-E microscope (Nikon AG Instruments), as described previously ^12,36^. Measurements were recorded with LightField (Princeton Instruments) in combination with a custom-built LabView (National Instruments) software for automation purposes. An exposure time of 10 seconds and laser excitation with band width of 10 nm and relative power of 100% was used for all measurements, unless stated otherwise.

Sample preparation was carried out in 384-well plates (MaxiSorp, Nunc). For all sensor response experiments, 49 *μL* of DNA- or LNA-SWCNT solution (10 mg/L) was added to a well and excited at 660 ± 5 nm. Following the acquisition of the initial spectrum, 0.5 μL of either CaCl_2_ solution (varying concentrations, dissolved in DI)) or DI water (for control experiments) was added, and the wells were sealed (Empore, 3M). Samples were incubated for 10-30 minutes (at room temperature) prior to recording the final spectrum. Following the acquisition of the spectra post the first addition, 0.5 μL of freshly prepared analyte solution (varying concentrations, dissolved in DI) was added. The mixture was incubated for an additional 10-30 minutes prior to acquiring the final spectra. A grating of 75 mm^-1^ was used to collect the nanotube emission between 900 – 1400 nm. Spectral fitting was performed using a custom Python program.

Photoluminescence excitation (PLE) maps were acquired between 525 nm and 800 nm with a 5 nm step. 50 μL aliquots of DNA- and LNA-SWCNT solutions (10 mg/L) were added to a 384-well plate, which was then sealed (Empore, 3M) to avoid evaporation during the measurement. The results were analysed using a custom Matlab code (Matlab R2017b, Mathworks).

### Surfactant replacement

Surfactant replacement of the DNA- and LNA-SWCNT solutions was performed using 1% (w/v) sodium deoxycholate (SDOC). All fluorescence experiments involving the real-time addition of SDOC were performed in a 384-well plate (MaxiSorp, Nunc). Following an initial time period of ~ 60 seconds to collect sufficient DNA/LNA-SWCNT control data, 2.5 μL of 1% SDOC was added to 22.5 μL of SWCNT solution (10 mg/L) to give a final SDOC concentration of 0.1%. Spectra were then collected continuously for 1 hour with a constant exposure time of 10 seconds.

Samples for absorbance experiments were prepared by mixing 63 μL of SWCNT solution (10 mg/L) with 7 μL of SDOC 1%. Spectra for the samples were collected prior to addition and at various time points post-addition. All measurements were carried out using a UV/vis/nIR scanning spectrometer (Shimadzu 3600 Plus) with a quartz cuvette (Suprasil quartz, path length 3 mm, Hellma).

### AFM imaging and sample preparation

DNA- and LNA-SWCNT solutions (20 μL) were drop-casted onto freshly cleaved mica substrates and dried at room temperature. Morphological characterization of the samples was performed using a commercial AFM setup (Cypher, Asylum Research) equipped with a commercial Si cantilever (AC240TS-R3, Asylum Research). Topography, phase, and amplitude images were acquired in standard tapping mode. Images were analysed with standard tools (e.g. plane subtraction and profile extraction) featured in the AFM data analysis software (Gwyddion 2.52).

### Zeta-potential measurements

Zeta potential measurements were carried out with a Zetasizer Nano ZS analyzer (Malvern), using folded capillary cells. All DNA- and LNA-SWCNT solutions were diluted in DI water to yield a final concentration of 2 mg/L.

### Circular dichroism measurements

Samples were analysed in a circular dichroism spectrometer across a wavelength range 200 – 300 nm in 0.2 nm intervals, using a 1-mm path length quartz cuvette (Suprasil quartz, Hellma). DNA and LNA samples were diluted to a final concentration of 7.5 μM in either DI water or 0.25 M CaCl_2_ and a volume of 300 μL was used for all experiments.

### *Drosophila* growth and neuron extraction

Tyrosine hydroxylase-GAL4 and UAS-CsChrimson animals were obtained from the Bloomington Drosophila Stock Center (Stock #8848, full genotype: w[*]; P{w[+mC]=ple-GAL4.F}3 and stock #55135, full genotype: w[1118]; P{y[+t7.7] w[+mC]=20XUAS-IVS-CsChrimson.mVenus}attP40, respectively). Parental flies were raised at 25 degrees and 50% relative humidity in 12hr:12hr dark:light cycles. Larval progeny of the above lines was raised in the dark, on food supplemented with all-trans retinal (~0.2 mM).

Drosophila neuronal cell culture was performed as described by Egger *et al.*^35^. Briefly, L3 larvae were picked from food vials, washed in 70% ethanol and placed in petri dishes containing supplemented Schneider’s medium (Schneider’s medium (Gibco, cat. no. 21720-024) supplemented with 1% (vol/vol) penicillin-streptomycin (1% Pen/Strep, Life Technologies, cat. no. 15070063) and 10% (vol/vol) heat-inactivated fetal bovine serum (10% FBS, Sigma)). Brains were dissected out and immediately rinsed with DPBS (Life Technologies) with 1% Pen/Strep prior to being processed for dissociation. Tissue was digested for 1 hr at RT with collagenase I (0.5 mg ml^-1^) (Sigma, cat. no. C9697). Digested tissue was rinsed with supplemented Schneider’s medium and finally resuspended in supplemented Schneider’s medium. Tissue was subsequently physically triturated by pipetting up and down at least 200 times.

### NIR widefield imaging

Samples were imaged using a custom-built optical setup consisting of an inverted microscope (Eclipse Ti-U, Nikon AG Instruments) with either an oil-immersion TIRF 100 x or 40 x objective (N.A. 1.49, Nikon) coupled to either an InGaAs camera (NIRvana 640 ST, Princeton Instruments) or Shamrock 303i spectrometer (Andor) coupled to EMCCD visible camera (iXon Ultra 888, Andor). For near-infrared images, samples were illuminated using a TriLine LaserBank system (Cairn Research) at 780 nm and fluorescence was collected using a 980 nm long-pass (LP) filter (Semrock). For visible images, samples were illuminated using LED excitation at either 395 nm (emission filter 417-477 nm) or 470 nm (emission filter 502-538 nm). Images were acquired using the Nikon NIS-Elements software (Nikon Instruments).

Nanotubes were coated onto poly-L-lysine coated glass petri dishes by incubating 10 μL of SWCNT solution (10 mg/L) and 40 μL DI water for 30 min followed by washing with DI water.

For single molecule images of nanotube responses in the absence of cells, coated petri dishes were initially imaged in the absence of any medium or buffer. Supplemented Schneider’s medium (10% FBS, 1% Pen/Strep) (200 μL) was added to the petri dish and the samples were imaged immediately, as well as following an incubation period of 5 minutes. Following this, 2.2 μL of freshly prepared dopamine solution (10 mM) was added. The samples were again imaged immediately and following 5 minutes of incubation. All imaging was performed using the TIRF 100 x objective, the 780 nm laser and the 980 nm LP emission filter with an exposure time of 1 second (relative power of 100%).

For cell culturing experiments, 150 μL of supplemented Schneider’s medium was added to the nanotube coated petri dishes. *Drosophila* neurons were prepared for culturing as detailed above and 50 μL of re-suspended brain solution (~50,000 neurons) was added to each petri dish and pipetted 2-3 times to mix. Cells were incubated at room temperature under ambient conditions overnight. Imaging of the samples was performed in both the visible and near-infrared.

## References

(1) Hyman, S. E. Neurotransmitters. Current Biology 2015, 15, 749–754.

(2) Kim, D. S.; Kang, E. S.; Baek, S.; Choo, S. S.; Chung, Y. H.; Lee, D.; Min, J.; Kim, T. H. Electrochemical detection of dopamine using periodic cylindrical gold nanoelectrode arrays. Scientific Reports 2018, 8, 1–10.

(3) Marceglia, S.; Foffani, G.; Bianchi, A. M.; Baselli, G.; Tamma, F.; Egidi, M.; Priori, A. Dopamine-dependent non-linear correlation between subthalamic rhythms in Parkinson’s disease. Journal of Physiology 2006, 571, 579–591.

(4) Kandimalla, R.; Reddy, P. H. Therapeutics of Neurotransmitters in Alzheimer’s Disease. Journal of Alzheimer’s Disease 2017, 57, 1049–1069.

(5) Jamwal, S.; Kumar, P. Insight Into the Emerging Role of Striatal Neurotransmitters in the Pathophysiology of Parkinson’s Disease and Huntington’s Disease: A Review. Current Neuropharmacology 2018, 17, 165–175.

(6) World Health Organisation, Neurological Disorders: Public Health Challenges; Nonserial Publication; World Health Organization, 2006.

(7) Gillen, A. J.; Boghossian, A. A. Non-covalent Methods of Engineering Optical Sensors Based on Single-Walled Carbon Nanotubes. Frontiers in Chemistry 2019, 7, 1–13.

(8) Beyene, A. G.; Alizadehmojarad, A. A.; Dorlhiac, G.; Goh, N.; Streets, A. M.; Král, P.; Vuković, L.; Landry, M. P. Ultralarge Modulation of Fluorescence by Neuromodulators in Carbon Nanotubes Functionalized with Self-Assembled Oligonucleotide Rings. Nano Letters 2018, 18, 6995–7003.

(9) Kruss, S.; Salem, D. P.; Vuković, L.; Lima, B.; Vander Ende, E.; Boyden, E. S.; Strano, M. S. High-resolution imaging of cellular dopamine efflux using a fluorescent nanosensor array. Proceedings of the National Academy of Sciences 2017, 114, 1789–1794.

(10) Kruss, S.; Landry, M. P.; Vander Ende, E.; Lima, B. M.; Reuel, N. F.; Zhang, J.; Nelson, J.; Mu, B.; Hilmer, A.; Strano, M. Neurotransmitter detection using corona phase molecular recognition on fluorescent single-walled carbon nanotube sensors. Journal of the American Chemical Society 2014, 136, 713–724.

(11) Beyene, A. G.; Delevich, K.; Del Bonis-O’Donnell, J. T.; Piekarski, D. J.; Lin, W. C.; Wren Thomas, A.; Yang, S. J.; Kosillo, P.; Yang, D.; Prounis, G. S. et al. Imaging striatal dopamine release using a nongenetically encoded near infrared fluorescent catecholamine nanosensor. Science Advances 2019, 5, 1–12.

(12) Lambert, B.; Gillen, A. J.; Schuergers, N.; Wu, S. J.; Boghossian, A. A. Directed evolution of the optoelectronic properties of synthetic nanomaterials. Chemical Communications 2019, 55, 3239–3242.

(13) Gillen, A. J.; Siefman, D. J.; Wu, S. J.; Bourmaud, C.; Lambert, B.; Boghossian, A. A. Templating colloidal sieves for tuning nanotube surface interactions and optical sensor responses. Journal of Colloid and Interface Science 2020, 565, 55–62.

(14) Jeong, S.; Yang, D.; Beyene, A.; Gest, A. M.; Landry, M. High Throughput Evolution of Near Infrared Serotonin Nanosensors. 2019, 673152.

(15) Godin, A. G.; Varela, J. A.; Gao, Z.; Danné, N.; Dupuis, J. P.; Lounis, B.; Groc, L.; Cognet, L. Single-nanotube tracking reveals the nanoscale organization of the extracellular space in the live brain. Nature Nanotechnology 2017, 12, 238–243.

(16) Paviolo, C.; Soria, F. N.; Ferreira, J. S.; Lee, A.; Groc, L.; Bezard, E.; Cognet, L. Nanoscale exploration of the extracellular space in the live brain by combining single carbon nanotube tracking and super-resolution imaging analysis. Methods 2020, 174, 91–99.

(17) Del Bonis-O’Donnell, J. T.; Page, R. H.; Beyene, A. G.; Tindall, E. G.; McFarlane, I. R.; Landry, M. P. Dual Near-Infrared Two-Photon Microscopy for Deep-Tissue Dopamine Nanosensor Imaging. Advanced Functional Materials 2017, 27, 1–10.

(18) Salem, D. P.; Gong, X.; Liu, A. T.; Koman, V. B.; Dong, J.; Strano, M. S. Ionic strength mediated phase transitions of surface adsorbed DNA on single-walled carbon nanotubes. Journal of the American Chemical Society 2017, 139, 16791–16802.

(19) Gillen, A. J.; Kupis-Rozmyslowicz, J.; Gigli, C.; Schürgers, N.; Boghossian, A. A. Xeno Nucleic Acid Nanosensors for Enhanced Stability Against Ion-induced Perturbations. The Journal of Physical Chemistry Letters 2018, acs.jpclett.8b01879.

(20) Eaton, M. E.; Macias, W.; Youngs, R. M.; Rajadhyaksha, A.; Dudman, J. T.; Konradi, C. L-type Ca2+ channel blockers promote Ca2+ accumulation when dopamine receptors are activated in striatal neurons. Brain Res Mol Brain Res 2004, 131, 65–72.

(21) Patel, J. C.; Witkovsky, P.; Avshalumov, M. V.; Rice, M. E. Mobilization of calcium from intracellular stores facilitates somatodendritic dopamine release. Journal of Neuroscience 2009, 29, 6568–6579.

(22) Brini, M.; Calì, T.; Ottolini, D.; Carafoli, E. Neuronal calcium signaling: Function and dysfunction. Cellular and Molecular Life Sciences 2014, 71, 2787–2814.

(23) Catoni, C.; Calì, T.; Brini, M. Calcium, dopamine and neuronal calcium sensor 1: Their contribution to Parkinson’s disease. Frontiers in Molecular Neuroscience 2019, 12, 1–8.

(24) Jena, P. V.; Safaee, M. M.; Heller, D. A.; Roxbury, D. DNA-Carbon Nanotube Complexation Affinity and Photoluminescence Modulation Are Independent. ACS Applied Materials and Interfaces 2017, 9, 21397–21405.

(25) Safaee, M. M.; Gravely, M.; Rocchio, C.; Simmeth, M.; Roxbury, D. DNA Sequence Mediates Apparent Length Distribution in Single-Walled Carbon Nanotubes. ACS Applied Materials and Interfaces 2019, 11, 2225–2233.

(26) Nißler, R.; Mann, F. A.; Chaturvedi, P.; Horlebein, J.; Meyer, D.; Vuković, L.; Kruss, S. Quantification of the Number of Adsorbed DNA Molecules on Single-Walled Carbon Nanotubes. Journal of Physical Chemistry C 2019, 123, 4837–4847.

(27) Campbell, J. F.; Tessmer, I.; Thorp, H. H.; Erie, D. A. Atomic Force Microscopy Studies of DNA-Wrapped Carbon Nanotube Structure and Binding to Quantum Dots. Journal of the American Chemical Society 2008, 130, 10648–10655.

(28) Choi, J. H.; Strano, M. S. Solvatochromism in single-walled carbon nanotubes. Applied Physics Letters 2007, 90, 88–91.

(29) Vogel, R.; Pal, A. K.; Jambhrunkar, S.; Patel, P.; Thakur, S. S.; Reátegui, E.; Parekh, H. S.; Saá, P.; Stassinopoulos, A.; Broom, M. F. High-Resolution Single Particle Zeta Potential Characterisation of Biological Nanoparticles using Tunable Resistive Pulse Sensing. Scientific Reports 2017, 7, 1–13.

(30) Bisker, G.; Dong, J.; Park, H. D.; Iverson, N. M.; Ahn, J.; Nelson, J. T.; Landry, M. P.; Kruss, S.; Strano, M. S. Protein-targeted corona phase molecular recognition. Nature Communications 2016, 7, 1–14.

(31) Heller, D. A.; Jeng, E. S.; Yeung, T.-K.; Martinez, B. M.; Moll, A. E.; Gastala, J. B.; Strano, M. S. Optical Detection of DNA Conformational Polymorphism on Single-Walled Carbon Nanotubes. Science 2006, 311, 508–511.

(32) Lee, M. A.; Nguyen, F. T.; Scott, K.; Chan, N. Y.; Bakh, N. A.; Jones, K. K.; Pham, C.; Garcia-Salinas, P.; Garcia-Parraga, D.; Fahlman, A. et al. Implanted Nanosensors in Marine Organisms for Physiological Biologging: Design, Feasibility, and Species Variability. ACS Sensors 2019, 4, 32–43.

(33) Pinals, R. L.; Yang, D.; Lui, A.; Cao, W.; Landry, M. P. Corona Exchange Dynamics on Carbon Nanotubes by Multiplexed Fluorescence Monitoring. Journal of the American Chemical Society 2020, 142, 1254–1264.

(34) Zubkovs, V.; Antonucci, A.; Schuergers, N.; Lambert, B.; Latini, A.; Ceccarelli, R.; Santinelli, A.; Rogov, A.; Ciepielewski, D.; Boghossian, A. A. Spinning-disc confocal microscopy in the second near-infrared window (NIR-II). Scientific Reports 2018, 8, 1–10.

(35) Egger, B.; Van Giesen, L.; Moraru, M.; Sprecher, S. G. In vitro imaging of primary neural cell culture from Drosophila. Nature Protocols 2013, 8, 958–965.

(36) Zubkovs, V.; Schuergers, N.; Lambert, B.; Ahunbay, E.; Boghossian, A. A. Mediatorless, Reversible Optical Nanosensor Enabled through Enzymatic Pocket Doping. Small 2017, 13, 1701654.

(37) Zhang, J.; Landry, M. P.; Barone, P. W.; Kim, J.-h. H.; Lin, S.; Ulissi, Z. W.; Lin, D.; Mu, B.; Boghossian, A. A.; Hilmer, A. J. et al. Molecular recognition using corona phase complexes made of synthetic polymers adsorbed on carbon nanotubes. Nature Nanotechnology 2013, 8, 959–968.

(38) Salem, D. P.; Landry, M. P.; Bisker, G.; Ahn, J.; Kruss, S.; Strano, M. S. Chirality dependent corona phase molecular recognition of DNA-wrapped carbon nanotubes. Carbon 2016, 97, 147–153.

(39) Harvey, J. D.; Jena, P. V.; Baker, H. A.; Zerze, G. H.; Williams, R. M.; Galassi, T. V.; Roxbury, D.; Mittal, J.; Heller, D. A. A carbon nanotube reporter of microRNA hybridization events in vivo. Nature Biomedical Engineering 2017, 1, 1–11.

(40) Polo, E.; Kruss, S. Impact of Redox-Active Molecules on the Fluorescence of Polymer-Wrapped Carbon Nanotubes. Journal of Physical Chemistry C 2016, 120, 3061–3070.

