## Supplementary information for "Distinguishing dopamine and calcium responses using XNA-nanotube sensors for improved neurochemical sensing"

The sequences used in this study are detailed in the table below where an asterisk indicates a locked base.

**Table 1: List of DNA and LNA sequences.**

| Name | Sequence |
|---|---|
| $(GT)_{15}$ | GTG TGT GTG TGT GTG TGT GTG TGT GTG TGT |
| LNA$_{every6T}$ | GT$^\star$G TGT GT$^\star$G TGT GT$^\star$G TGT GT$^\star$G TGT GT$^\star$G TGT |
| LNA$_{every6G}$ | G$^\star$TG TGT G$^\star$TG TGT G$^\star$TG TGT G$^\star$TG TGT G$^\star$TG TGT |
| LNA$_{every5}$ | GTG TG$^\star$T GTG T$^\star$GT GTG$^\star$ TGT GT$^\star$G TGT G$^\star$TG TGT$^\star$ |
| LNA$_{20mid}$ | GTG TGT GTG TGT G$^\star$T$^\star$G$^\star$ T$^\star$G$^\star$T$^\star$ GTG TGT GTG TGT |
| LNA$_{20end}$ | G$^\star$T$^\star$G$^\star$ TGT GTG TGT GTG TGT GTG TGT GTG T$^\star$G$^\star$T$^\star$ |
| LNA$_{every3}$ | GTG$^\star$ TGT$^\star$ GTG$^\star$ TGT$^\star$ GTG$^\star$ TGT$^\star$ GTG$^\star$ TGT$^\star$ GTG$^\star$ TGT$^\star$ |
| LNA$_{33mid}$ | GTG TGT GTG TG$^\star$T$^\star$ G$^\star$T$^\star$G$^\star$ T$^\star$G$^\star$T$^\star$ G$^\star$T$^\star$G TGT GTG TGT |

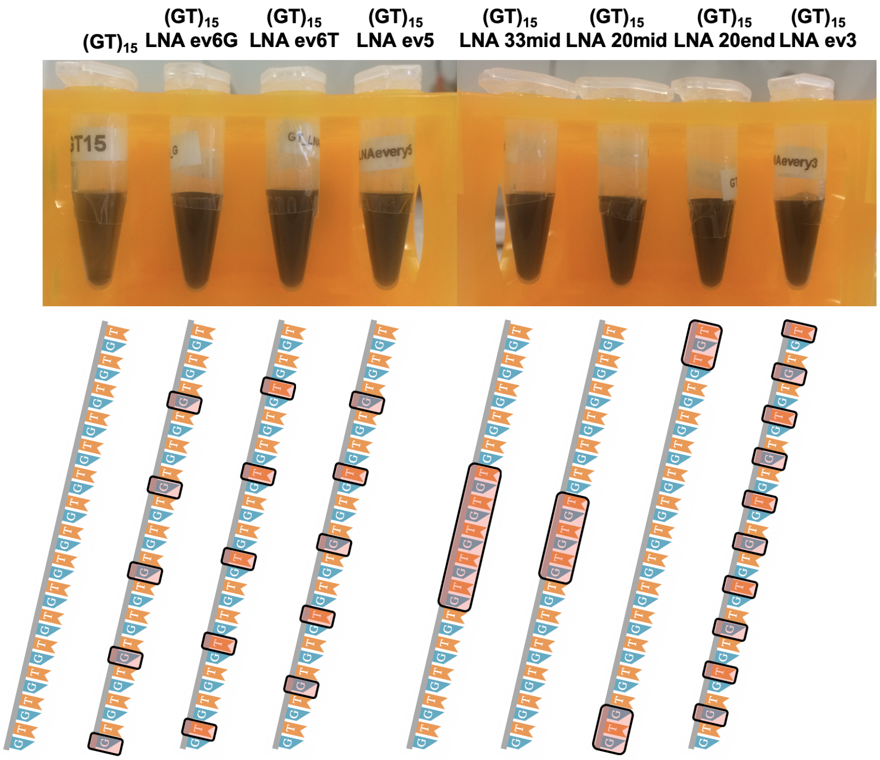


Figure 1: Max concentration solutions of DNA- and LNA-SWCNT suspension post sonication, dialysis and centrifugation.

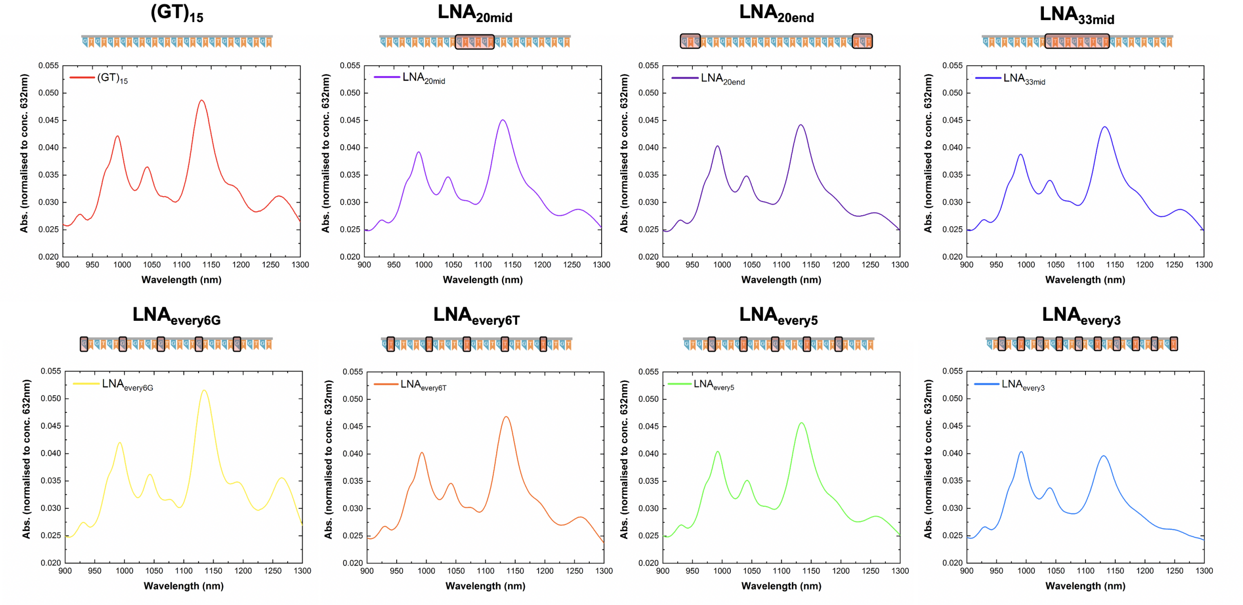


Figure 2: Absorbance spectra for all DNA- and LNA-SWCNT samples examined in this study. All spectra are normalized to concentration as determined using an extinction coefficient of 0.036 mg/L at $Abs_{632nm}$.

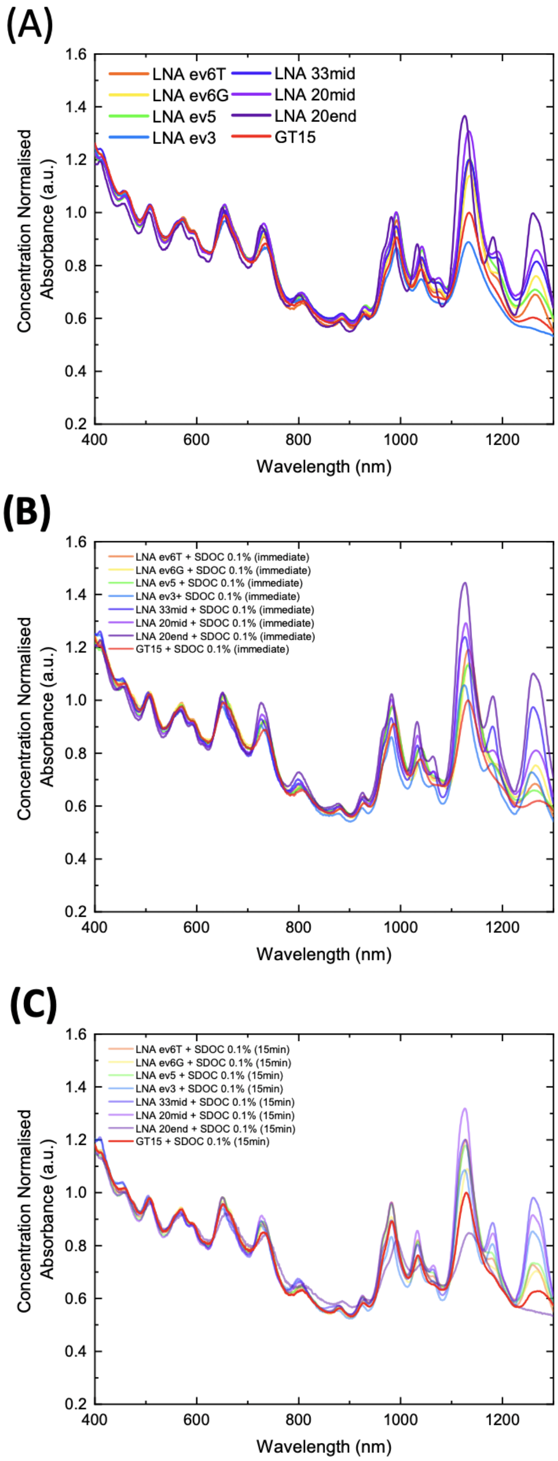


Figure 3: Absorbance spectra of all sequences used in this study before and after SDOC replacement. **(A)** Before, **(B)** immediately following the addition of SDOC (0.1% final concentration), and **(C)** following 15 min incubation post-addition. All spectra are normalized to concentration as determined using an extinction coefficient of 0.036 mg/L at $Abs_{632nm}$ and subsequently normalized to the $(GT)_{15}$ spectrum for comparison.

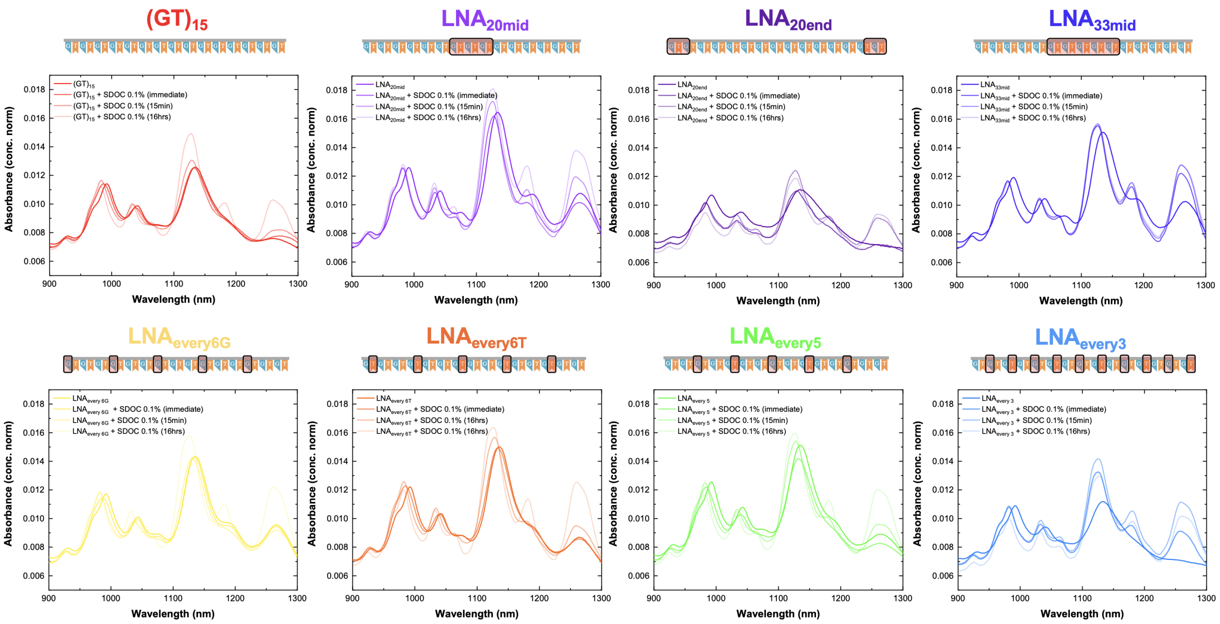


Figure 4: Absorbance spectra for all DNA- and LNA-SWCNT samples following addition of SDOC (0.1% final concentration) at various timepoints: immediately after addition, following 15 mins of incubation, and following 16 hours of incubation. All spectra are normalized to concentration as determined using an extinction coefficient of 0.036 mg/L at $Abs_{632nm}$.

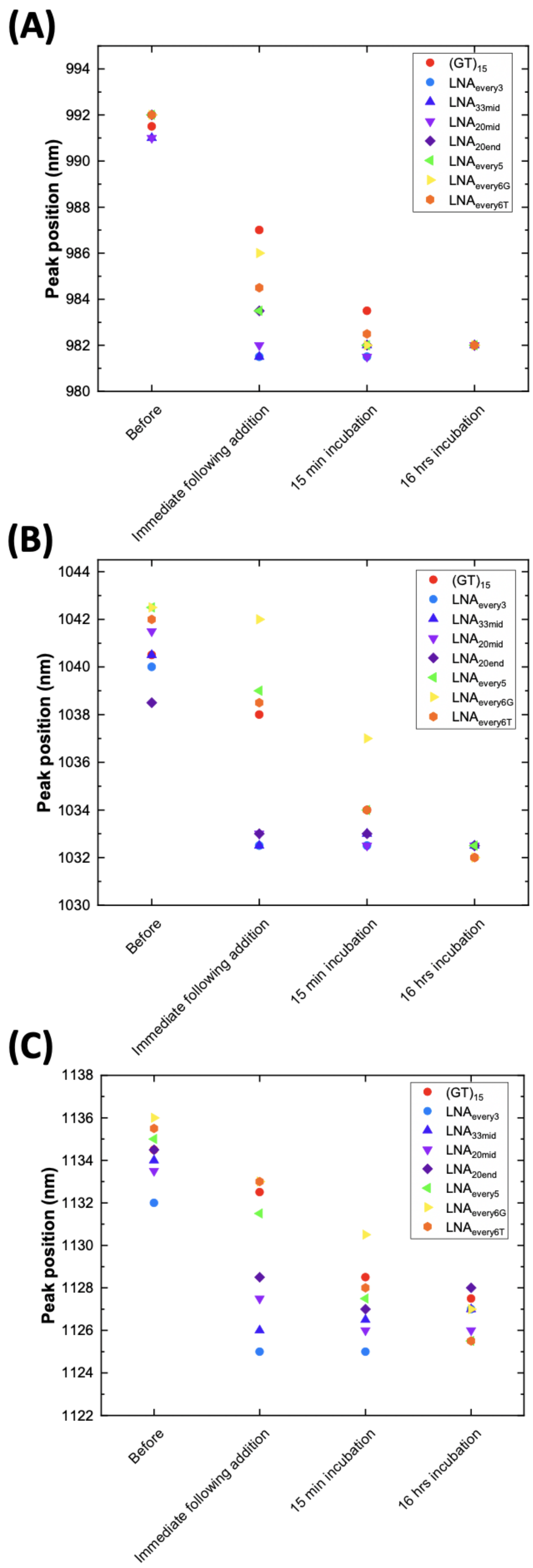


Figure 5: Wavelength position of the absorbance peaks at **(A)** ~ 990 nm, **(B)** ~ 1040 nm, and **(C)** ~ 1135 nm before SDOC and at three time points following SDOC addition (final SDOC concentration of 0.1%).

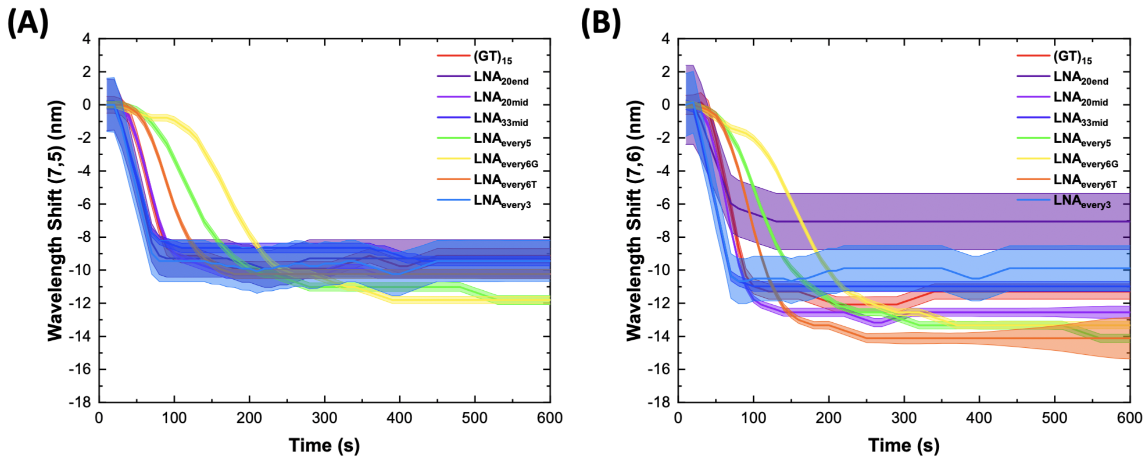


Figure 6: Wavelength shift of the fluorescence emission wavelength position of the **(A)** (7,5) and **(B)** (7,6) chirality peaks of all sequences parameters as a function of time following SDOC addition (0.1% final SDOC concentration).

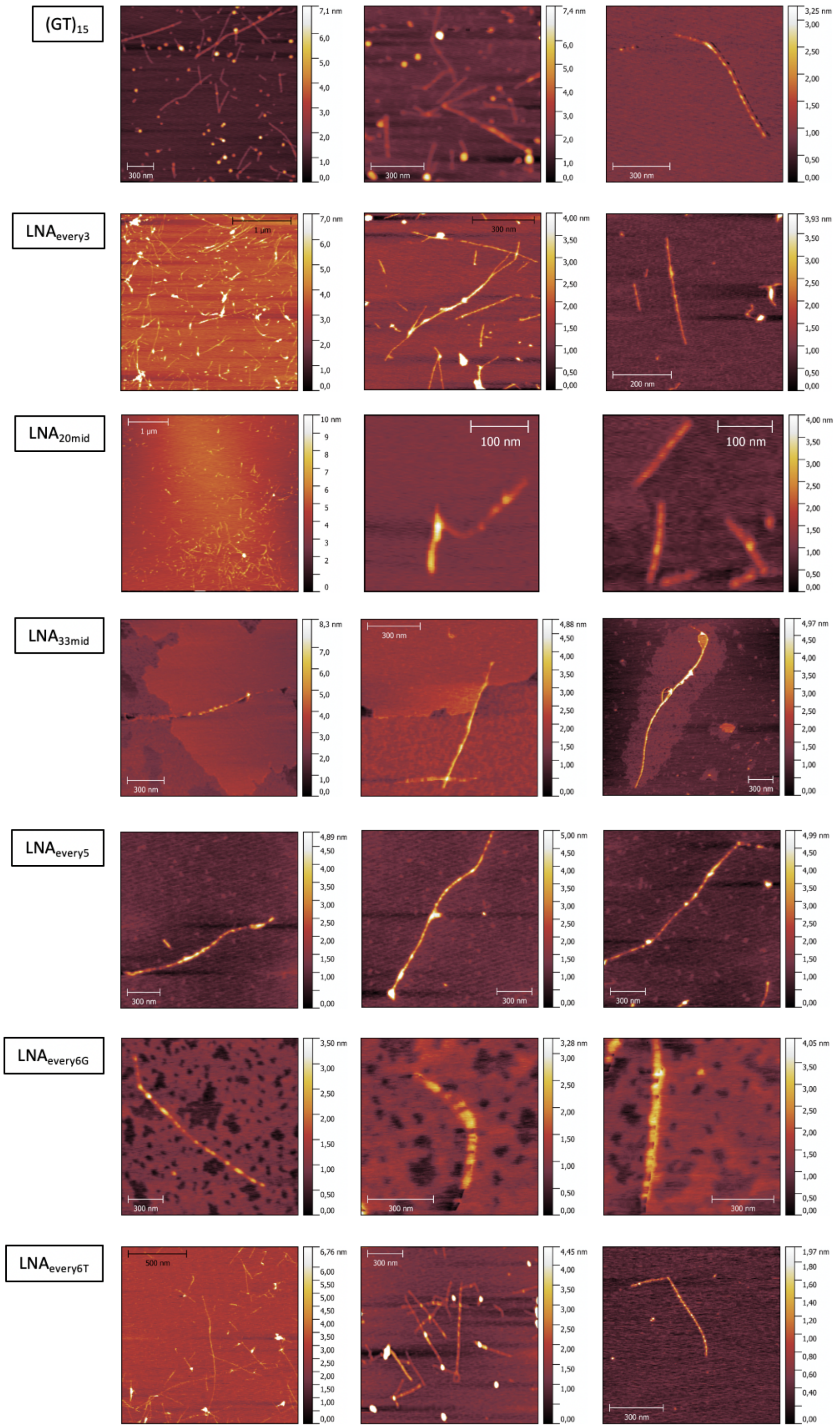


Figure 7: Additional AFM images collected for $(GT)_{15}$-, $LNA_{every3}$-, $LNA_{every5}$-, $LNA_{every6G}$-, $LNA_{every6T}$-, $LNA_{33mid}$-, and $LNA_{20mid}$-SWCNTs.

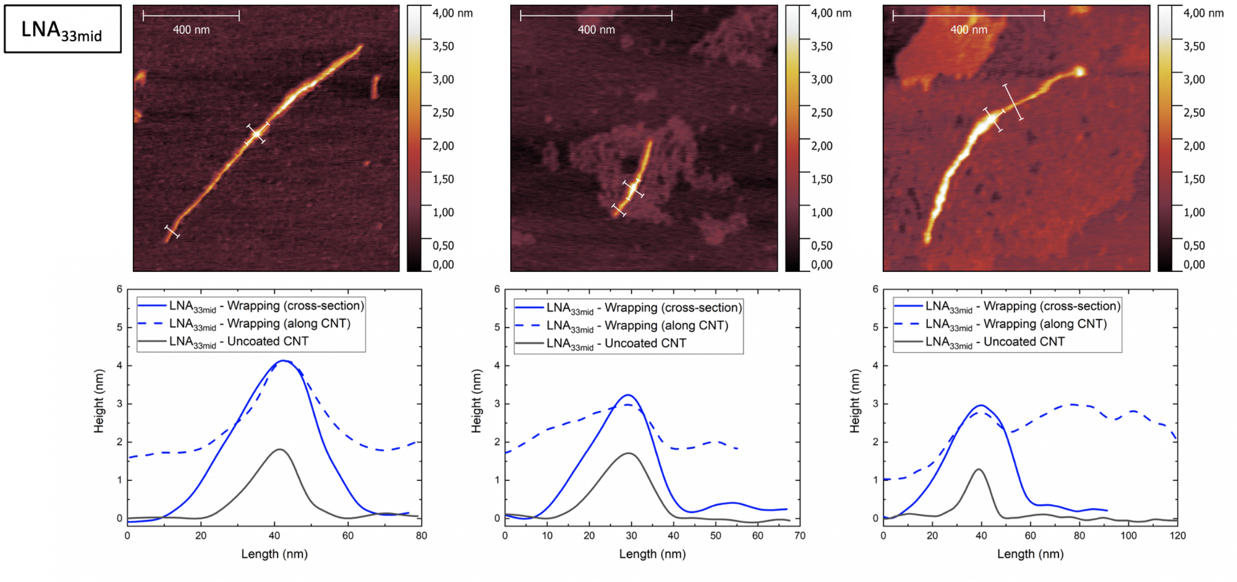


Figure 8: AFM height images (**top**) and extracted profiles (**bottom**) for $LNA_{33mid}$-SWCNTs. White lines indicate where height profiles were extracted from. The peak height extracted from cross-sectional and longitudinal profiles show good agreement. Areas of dense continuous coverage are indicated by the continuously higher peak values extracted in the profiles along the nanotube.

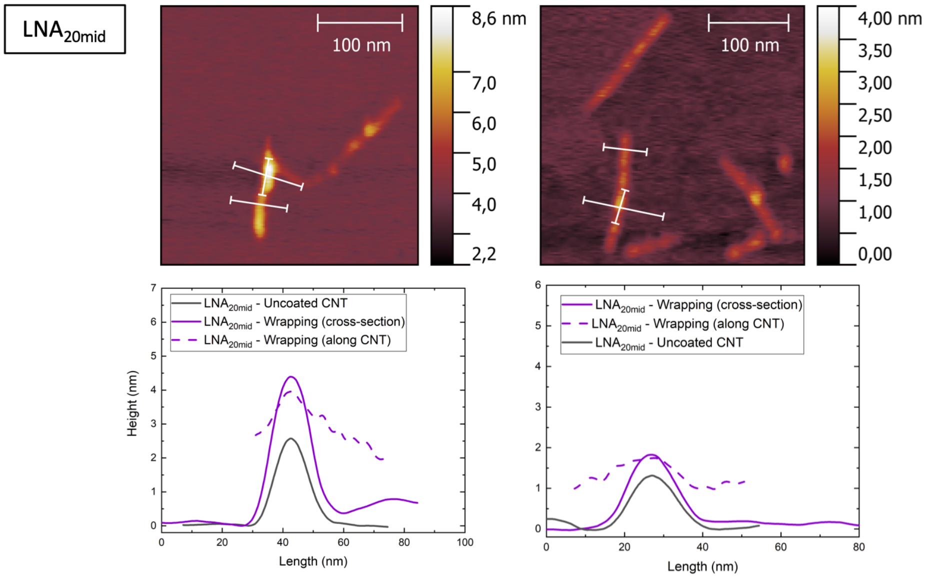


Figure 9: AFM height images (**top**) and extracted profiles (**bottom**) for $LNA_{20mid}$-SWCNTs. White lines indicate where height profiles were extracted from. The peak height extracted from cross-sectional and longitudinal profiles show good agreement. Areas of dense continuous coverage are indicated by the continuously higher peak values extracted in the profiles along the nanotube.

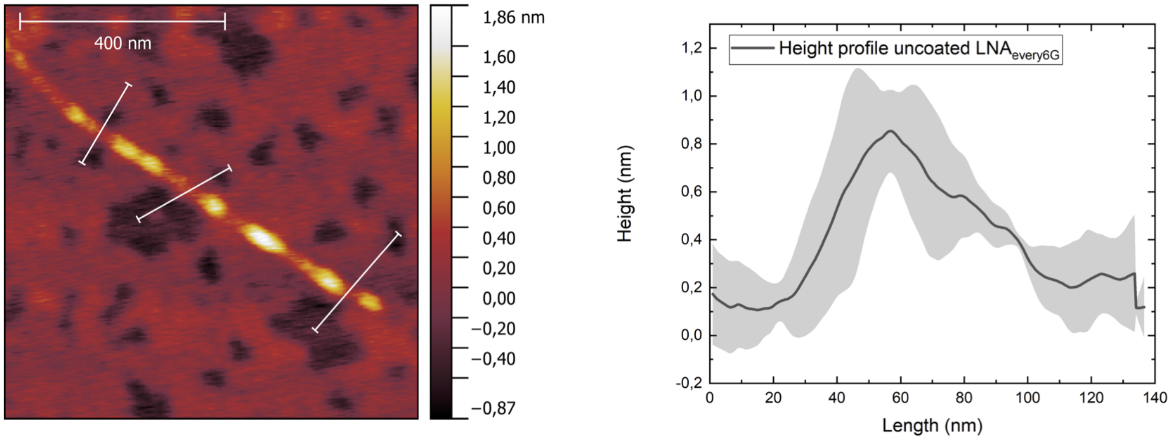


Figure 10: Due to the presence of an additional layer of material on the substrate of the $LNA_{every6G}$-SWCNTs, the heights estimated by the cross-section analysis shown in **Figure 3** were slightly underestimated. Additional measurements, taken from clear parts of the substrate show the true height of the nanotubes, which is used for comparison to the other wrappings. Average height profiles were extracted from three positions selected along the nanotube that overlapped with bare substrate (indicated by the white lines in AFM height images). Solid lines in the height profile represent the average height and shaded region represents $1\sigma$ standard deviation (n = 3 profiles).

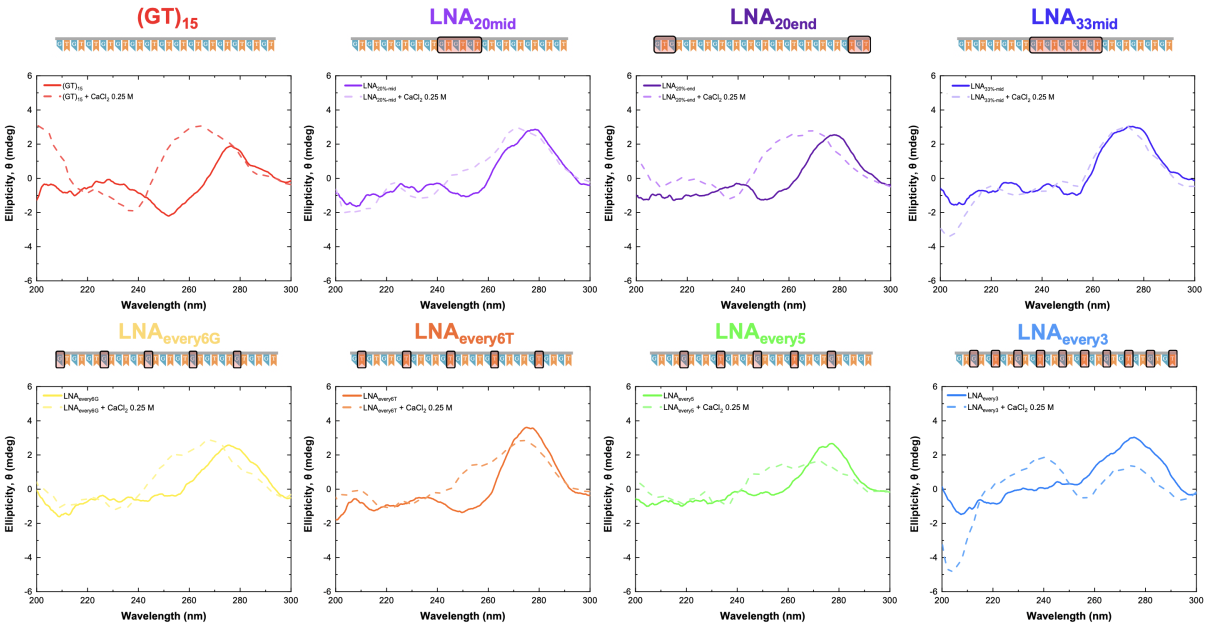


Figure 11: Circular dichroism measurements of DNA$(GT)_{15}$ and LNA$(GT)_{15}$ sequences on addition of $CaCl_2$.

**Table 2: Fit parameters for the solvatochromism shift model. The slope, Adjusted R-squared, effective dielectric constant ($\epsilon_{\text{eff}}$), and the relative surface coverage ($\alpha$).**

| Sequence | Slope | Adj. $R^2$ | $\epsilon_{\text{eff}}$ | $\alpha$ |
|---|---|---|---|---|
| $(GT)_{15}$ | 0.05886 ± 0.00622 | 0.93647 | 9.44792178 | 0.935221 |
| $LNA_{every6T}$ | 0.05888 ± 0.00625 | 0.93601 | 9.459323006 | 0.935085 |
| $LNA_{every6G}$ | 0.0589 ± 0.00636 | 0.93392 | 9.470750395 | 0.934949 |
| $LNA_{every5}$ | 0.05969 ± 0.00627 | 0.93728 | 9.944103765 | 0.929321 |
| $LNA_{every3}$ | 0.06574 ± 0.01069 | 0.88053 | 15.91040576 | 0.858378 |
| $LNA_{33mid}$ | 0.05797 ± 0.00427 | 0.97347 | 8.965713056 | 0.940955 |
| $LNA_{20mid}$ | 0.05815 ± 0.00547 | 0.94922 | 9.059429799 | 0.93984 |
| $LNA_{20end}$ | 0.05886 ± 0.00622 | 0.93647 | 9.44792178 | 0.935221 |

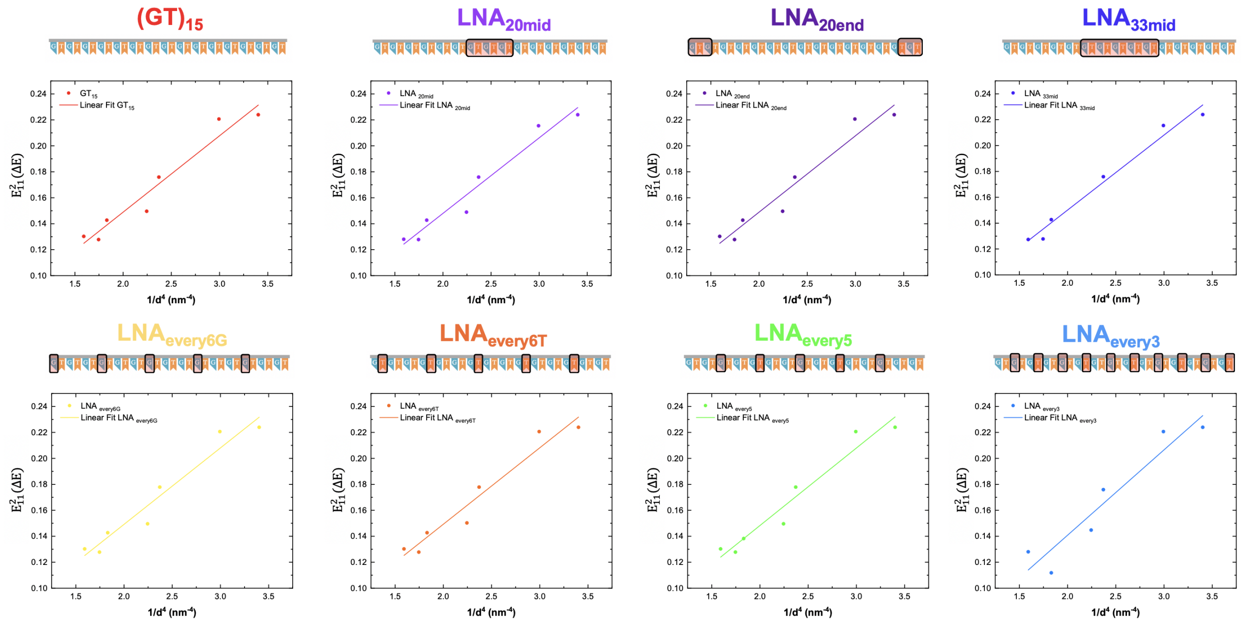


Figure 12: The solvatochromic shift as a function of the SWCNT diameter to the power of negative 4 ($d^{-4}$, **dots**) and its linear fit (**solid line**).

| **Chirality** | $\frac{1}{d^4}$ | $\mathbf{E}_{11}^{2}(\Delta\mathbf{E})$ | | | | | | | |
|---|---|---|---|---|---|---|---|---|---|
| **(n,m)** | $\mathbf{nm}^{-4}$ | $\mathbf{GT}_{15}$ | $\mathbf{LNA_{every6T}}$ | $\mathbf{LNA_{every6G}}$ | $\mathbf{LNA_{every5}}$ | $\mathbf{LNA_{every3}}$ | $\mathbf{LNA_{33mid}}$ | $\mathbf{LNA_{20mid}}$ | $\mathbf{LNA_{20end}}$ |
| (9,4) | 1.593 | 0.13024 | 0.13024 | 0.13024 | 0.13024 | 0.12805 | 0.12731 | 0.12805 | 0.13024 |
| (7,6) | 1.747 | 0.12777 | 0.12777 | 0.12777 | 0.12777 | | 0.12777 | 0.12777 | 0.12777 |
| (10,2) | 1.832 | 0.14272 | 0.14272 | 0.14272 | 0.1382 | 0.11179 | 0.14272 | 0.14272 | 0.14272 |
| (8,4) | 2.246 | 0.14961 | 0.15029 | 0.14961 | 0.14961 | 0.14475 | | 0.14892 | 0.14961 |
| (7,5) | 2.371 | 0.17591 | 0.17784 | 0.17784 | 0.17784 | 0.17591 | 0.17591 | 0.17591 | 0.17591 |
| (8,3) | 2.994 | 0.22067 | 0.22067 | 0.22067 | 0.22067 | 0.22067 | 0.21548 | 0.21548 | 0.22067 |
| (6,5) | 3.402 | 0.22392 | 0.22392 | 0.22392 | 0.22392 | 0.22392 | 0.22392 | 0.22392 | 0.22392 |

**Table 3:** Energy shift, $E_{11}^{2}(\Delta E)$, as a function of nanotube chirality for all DNA and LNA sequences. Energy shifts showed a decreasing trend with increasing diameter in agreement with previous reports[28].

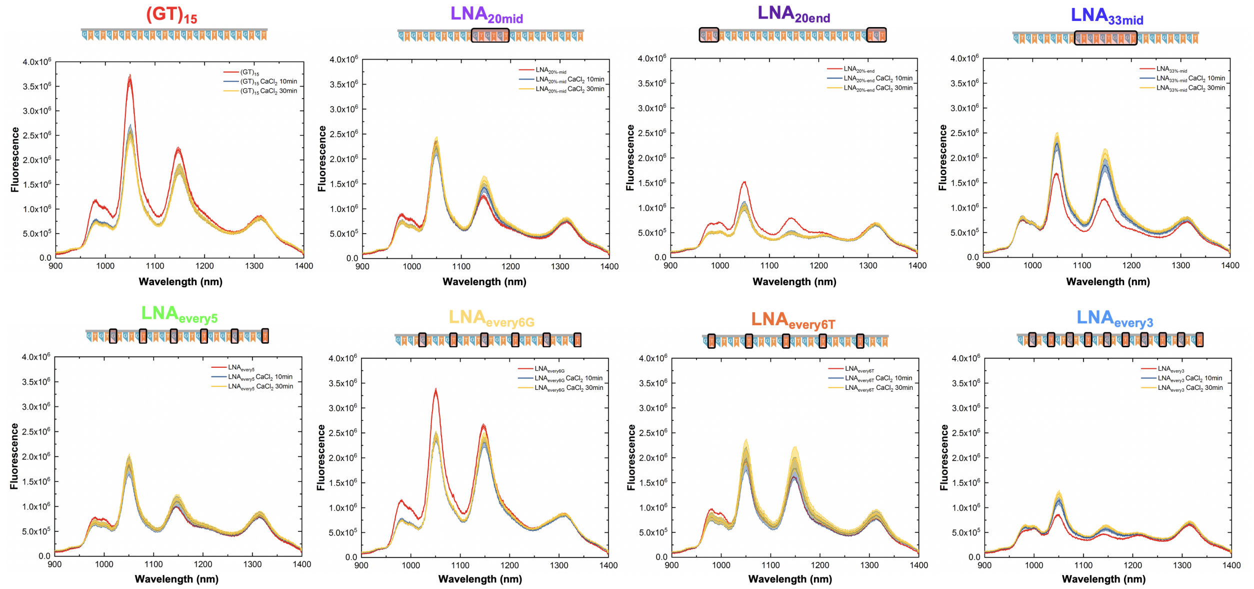


Figure 13: Spectral response of DNA and LNA sensors following the addition of 0.5 M $CaCl_2$ (final concentration: 5 mM, excitation: 660 nm). Graphs include the spectra before addition (**red**) and following either 10 min (**blue**) or 30 min (**yellow**) of incubation post addition. As no significant difference was observed between 10 min and 30 min incubation, 30 min incubations were used for all comparisons of wavelength shifting and intensity changes. The solid line represents the average wavelength shift with the shaded regions representing $1\sigma$ standard deviation (n = 3 technical replicates).

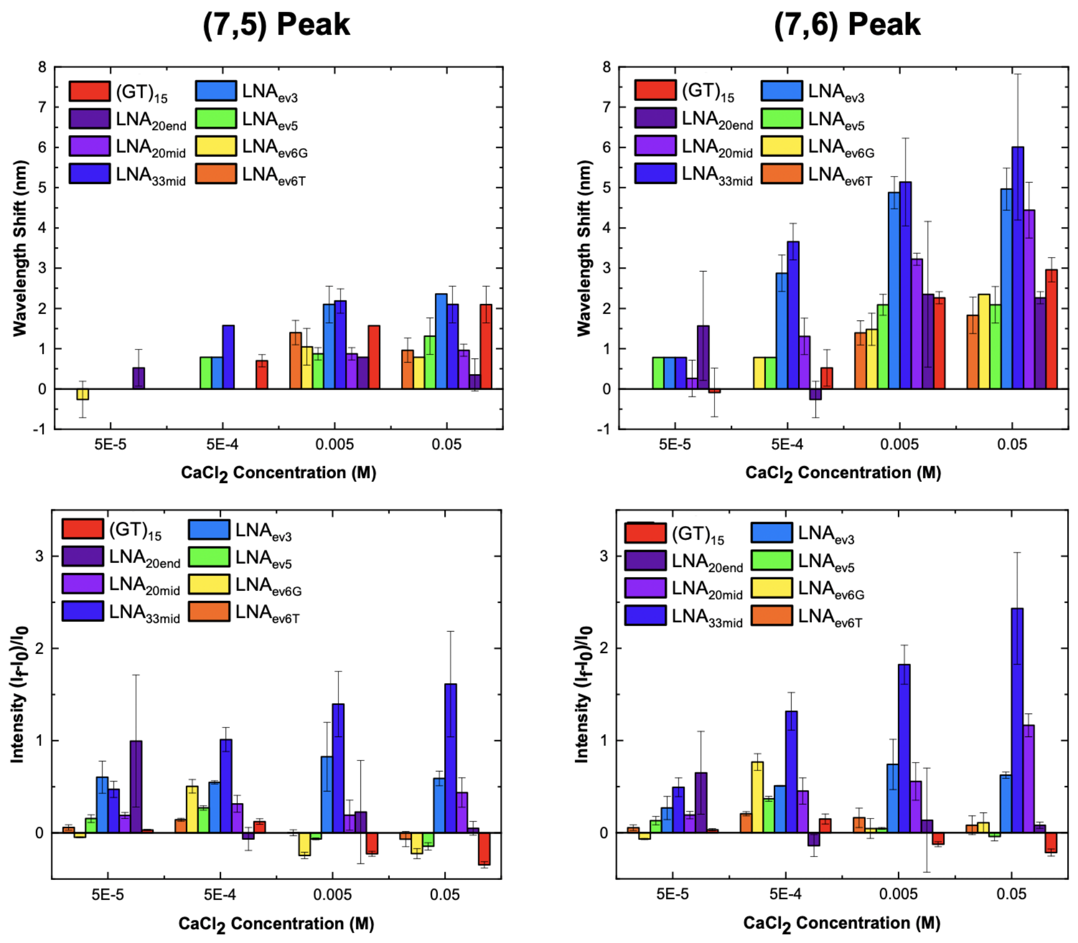


Figure 14: Response of DNA- and LNA- sensors to $CaCl_2$. Concentration-dependent (**top**) wavelength shift and (**bottom**) intensity change of the (7,5) peak (**left**) and (7,6) peak (**right**) following the addition of $CaCl_2$ (excitation: 660 nm). All samples were incubated for 30 min. Error bars represent $1\sigma$ standard deviation (n = 3 – 9 technical replicates).

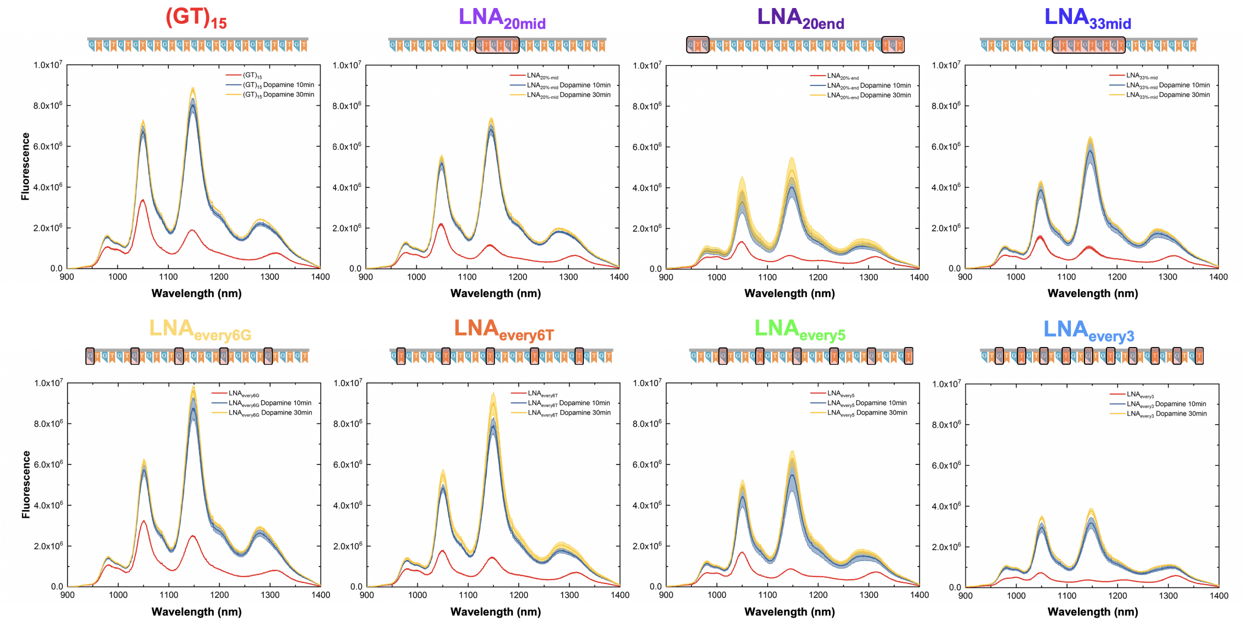


Figure 15: Spectral response of LNA sensors following the addition of 10 mM dopamine (final concentration: 100 $\mu$M, excitation: 660 nm). Graphs include the spectra before addition (**red**) and following either 10 min (**blue**) or 30 min (**yellow**) of incubation post addition. The solid line represents the average wavelength shift with the shaded regions representing $1\sigma$ standard deviation (n = 3 technical replicates).

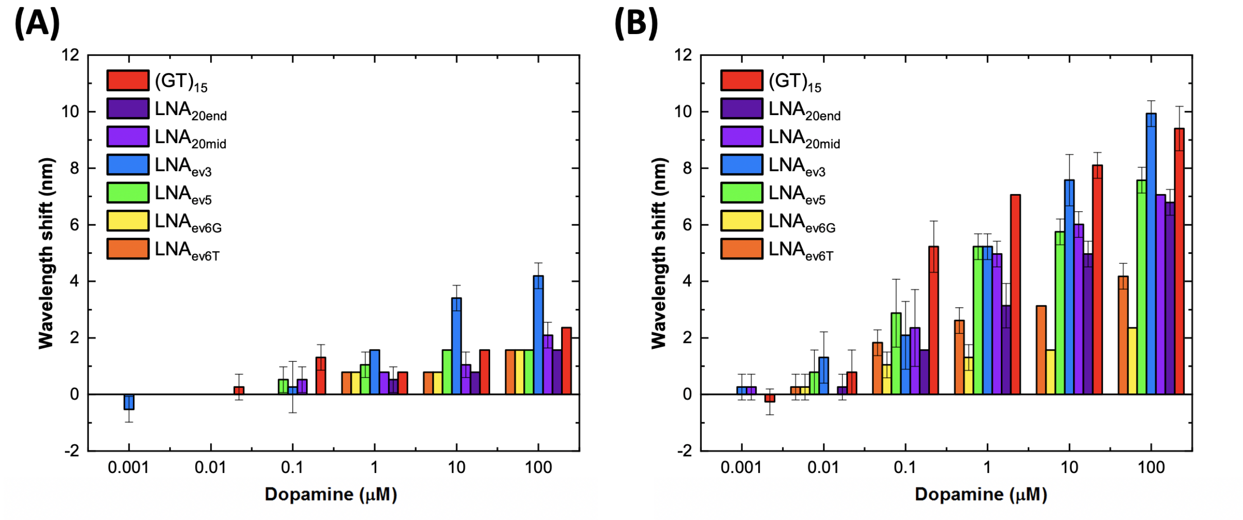


Figure 16: Concentration-dependent wavelength shifting response of DNA- and LNA-SWCNTs towards dopamine in the absence of $CaCl_2$ (excitation: 660 nm). Shift in the wavelength position of the **(A)** (7,5) and **(B)** (7,6) peak following the addition of dopamine. All samples were incubated for 30 min. Error bars represent $1\sigma$ standard deviation (n = 3 technical replicates).

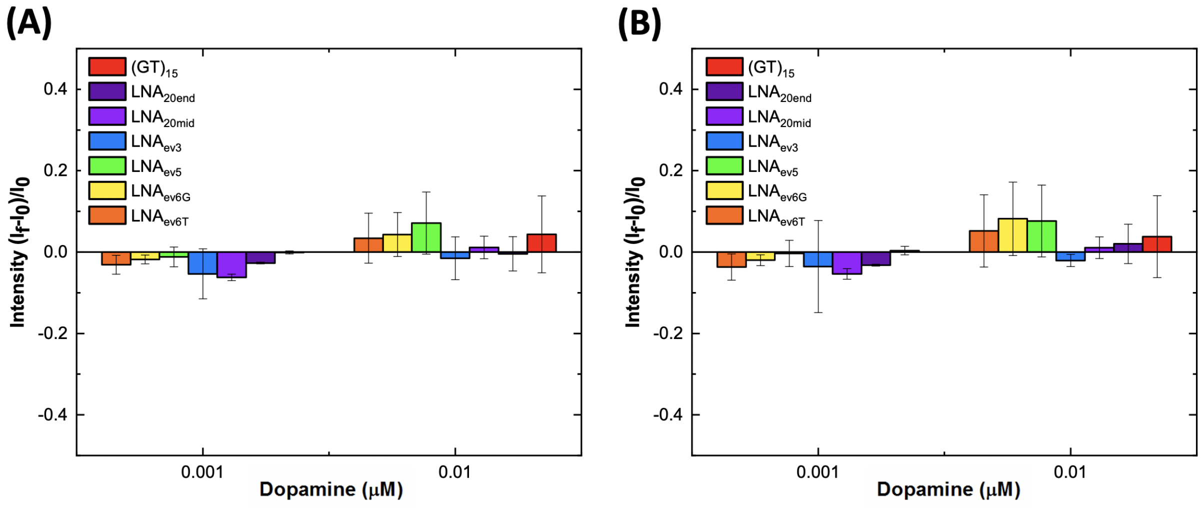


Figure 17: Concentration-dependent intensity response of DNA- and LNA-SWCNTs towards dopamine in the absence of $CaCl_2$ (excitation: 660 nm) at low concentration. Intensity change of the **(A)** (7,5) and **(B)** (7,6) peak following the addition of dopamine. All samples were incubated for 30 min. Error bars represent $1\sigma$ standard deviation (n = 3 technical replicates).

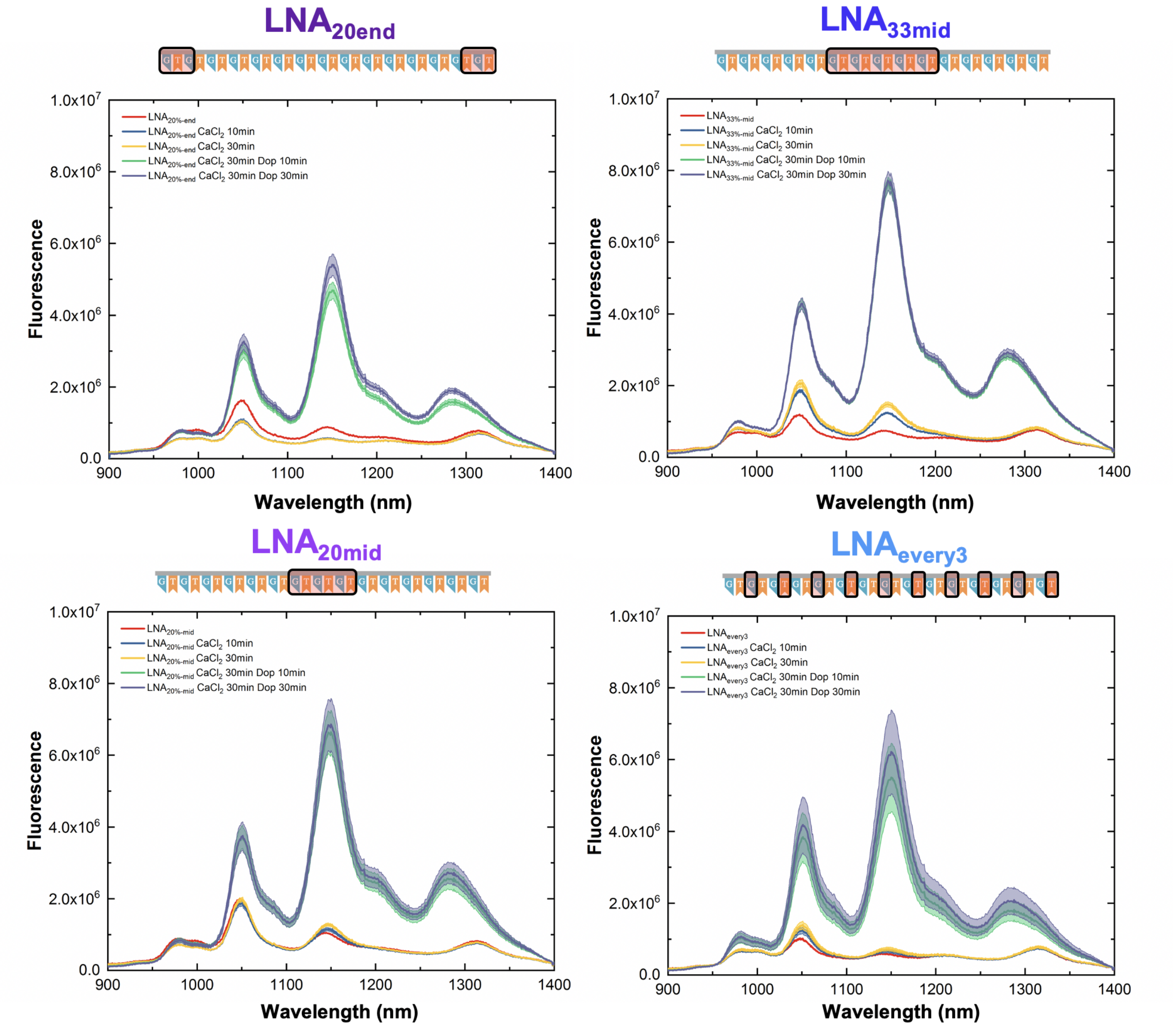


Figure 18: Spectral response of LNA sensors following the addition of 0.5 M $CaCl_2$ (final concentration: 5 $\mu$M) and 10 mM dopamine (final concentration: 100 $\mu$M, excitation: 660 nm). Graphs include the spectra before addition (**red**) and following the addition of $CaCl_2$ after 10 min (**blue**) and 30 min (**yellow**) of incubation and following subsequent addition of dopamine at 10 min (**green**) and 30 min (**purple**) post addition. The solid line represents the average wavelength shift with the shaded regions representing $1\sigma$ standard deviation (n = 3 technical replicates).

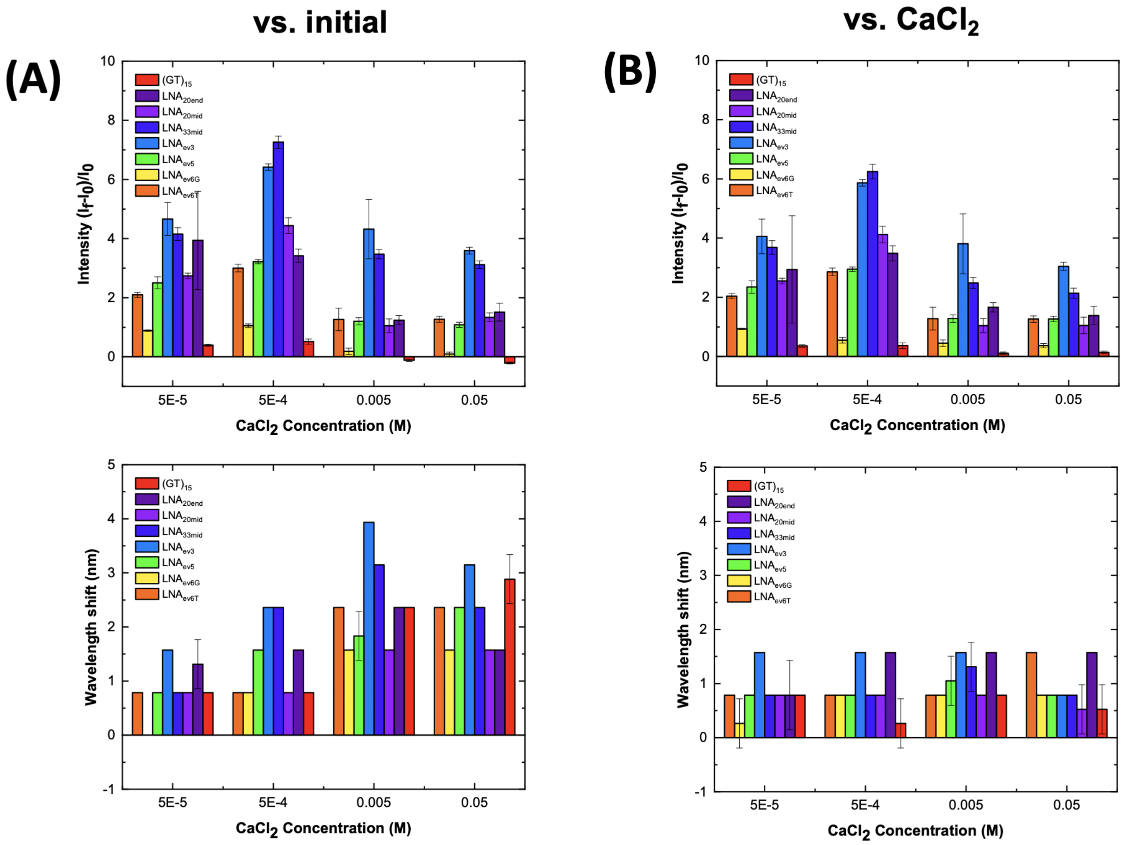


Figure 19: Fluorescence response of $(GT)_{15}$- and LNA-SWCNT sensors following the addition of 10 mM dopamine (final concentration: 100 $\mu$M) in the presence of various concentrations of $CaCl_2$ (excitation: 660 nm). Intensity (**top**) and wavelength (**bottom**) response of the (7,5) peak following dopamine addition were calculated versus **(A)** the initial spectrum (before $CaCl_2$ addition) and **(B)** the spectrum following incubation with the $CaCl_2$ solutions highlighting the bias that can be introduced to the sensor response in the presence of $CaCl_2$. All samples were incubated for 30 min following dopamine addition. Error bars represent 1$\sigma$ standard deviation (n = 3 technical replicates).

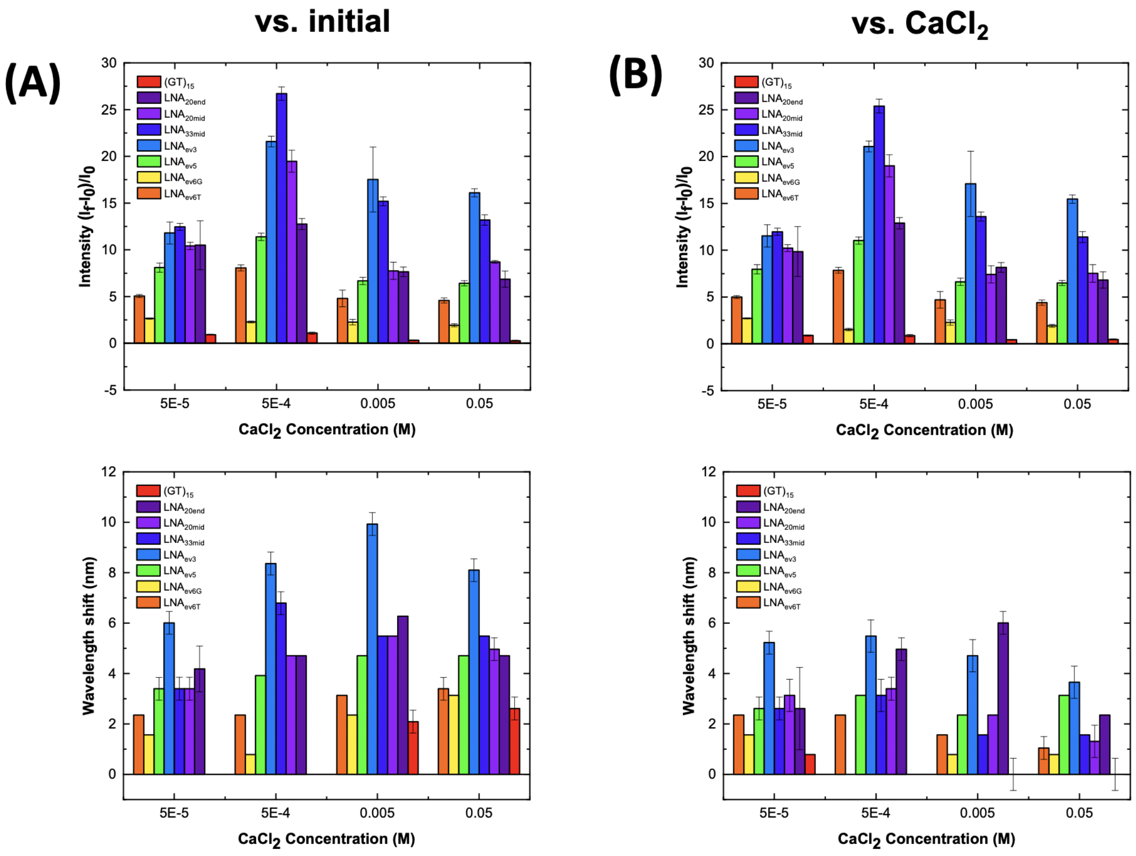


Figure 20: Fluorescence response of $(GT)_{15}$- and LNA-SWCNT sensors following the addition of 10 mM dopamine (final concentration: 100 $\mu$M) in the presence of various concentrations of $CaCl_2$ (excitation: 660 nm). Intensity (**top**) and wavelength (**bottom**) response of the (7,6) peak following dopamine addition were calculated versus **(A)** the initial spectrum (before $CaCl_2$ addition) and **(B)** the spectrum following incubation with the $CaCl_2$ solutions highlighting the bias that can be introduced to the sensor response in the presence of $CaCl_2$. All samples were incubated for 30 min following dopamine addition. Error bars represent $1\sigma$ standard deviation (n = 3 technical replicates).

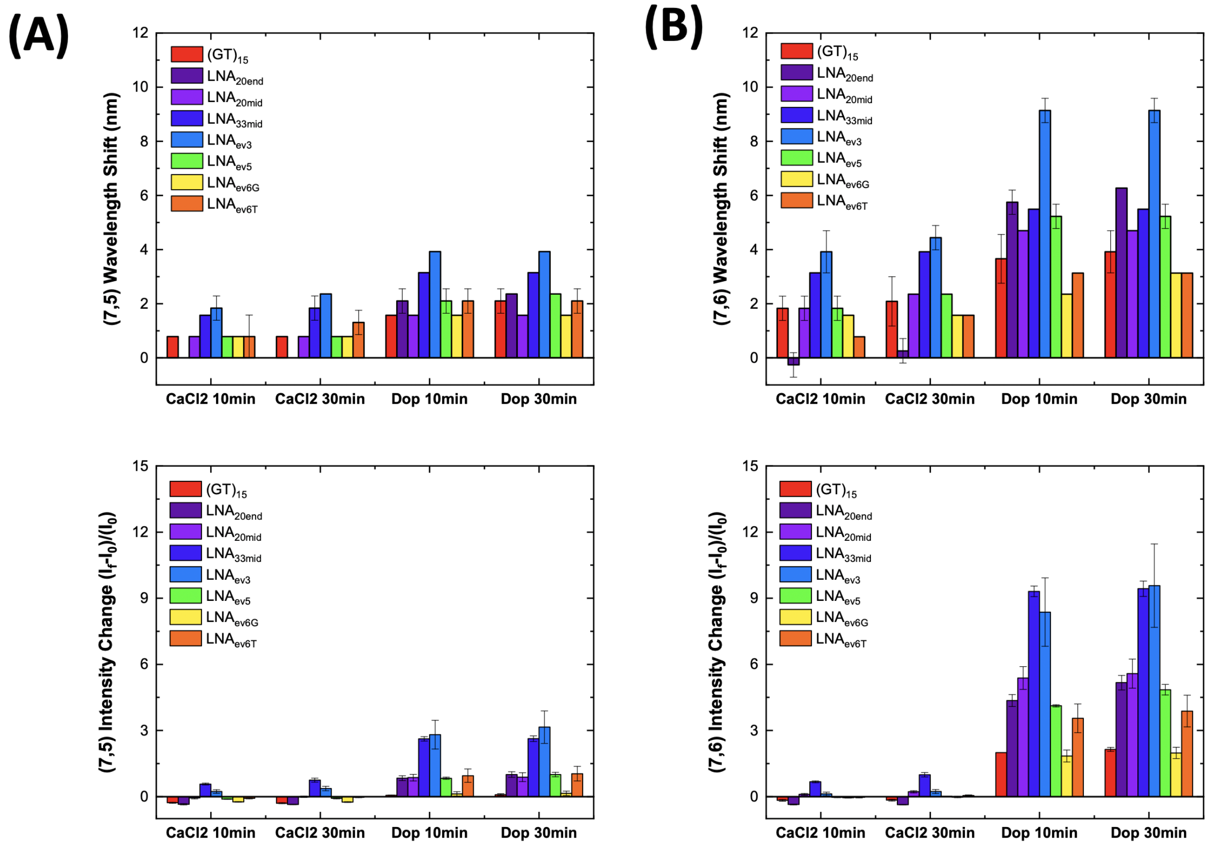


Figure 21: Comparison of the fluorescence response of $(GT)_{15}$- and LNA-SWCNTs for the **(A)** (7,5) and **(B)** (7,6) peaks following the addition of 0.5 M $CaCl_2$ (final concentration: 5 mM) and 10 mM dopamine (final concentration: 100 $\mu$M, excitation: 660 nm). Wavelength (**top**) and intensity (**bottom**) response of the (7,5) and (7,6) peaks following dopamine addition were calculated versus the initial spectrum (before initial $CaCl_2$ addition) for both time points. Error bars represent 1$\sigma$ standard deviation (n = 3 technical replicates).

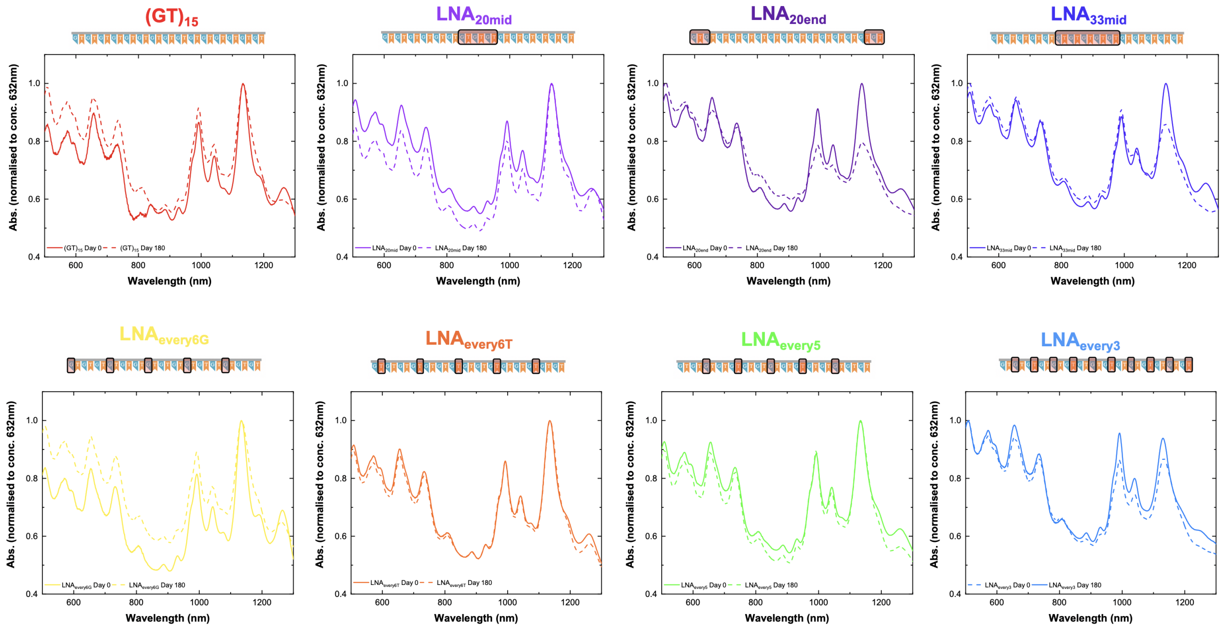


Figure 22: Effect of ageing on the absorbance spectra of all samples examined in this study. Absorbance spectra were collected immediately following suspension (**solid line**) and following 180 days incubation at 4°C (**dashed line**). All spectra are normalized to concentration using an extinction coefficient of 0.036 mg/L at $Abs_{632nm}$.

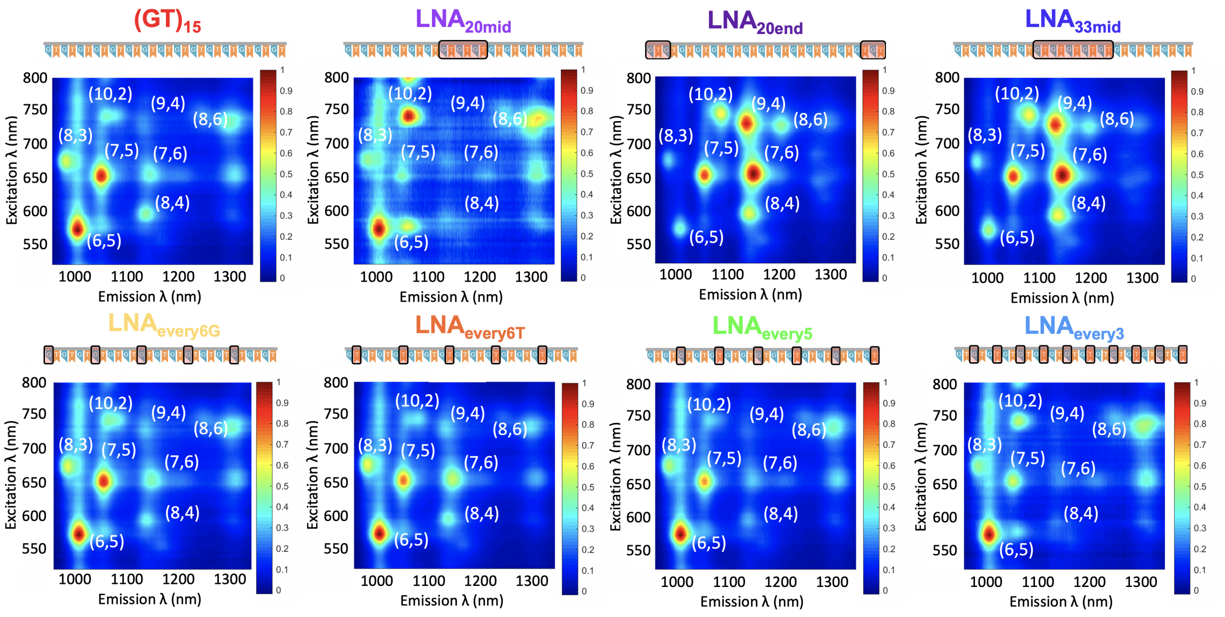


Figure 23: Effect of ageing on the fluorescence properties of all samples examined in this study. PLE maps of the original $(GT)_{15}$- and all LNA-SWCNT solutions following 180 days incubation at 4°C. All PLE maps were normalized to the maximum fluorescence intensity for comparison.

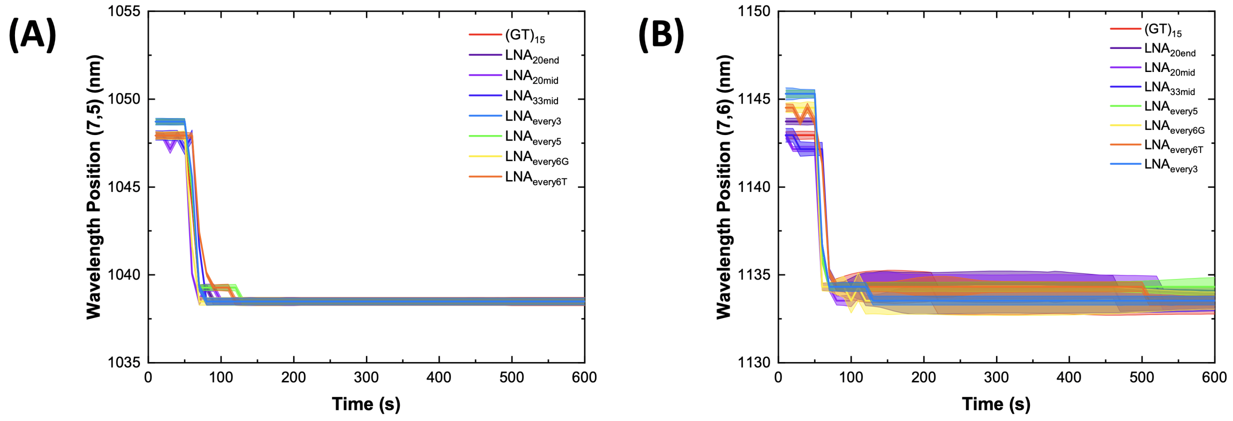


Figure 24: Modulation of fluorescence emission wavelength position of the **(A)** (7,5) and **(B)** (7,6) chirality peaks for all sequences as a function of time following SDOC addition. All SWCNT suspensions were prepared via the surfactant exchange method. The shaded region represents the error ($3\sigma$) of the peak fit at each time point.

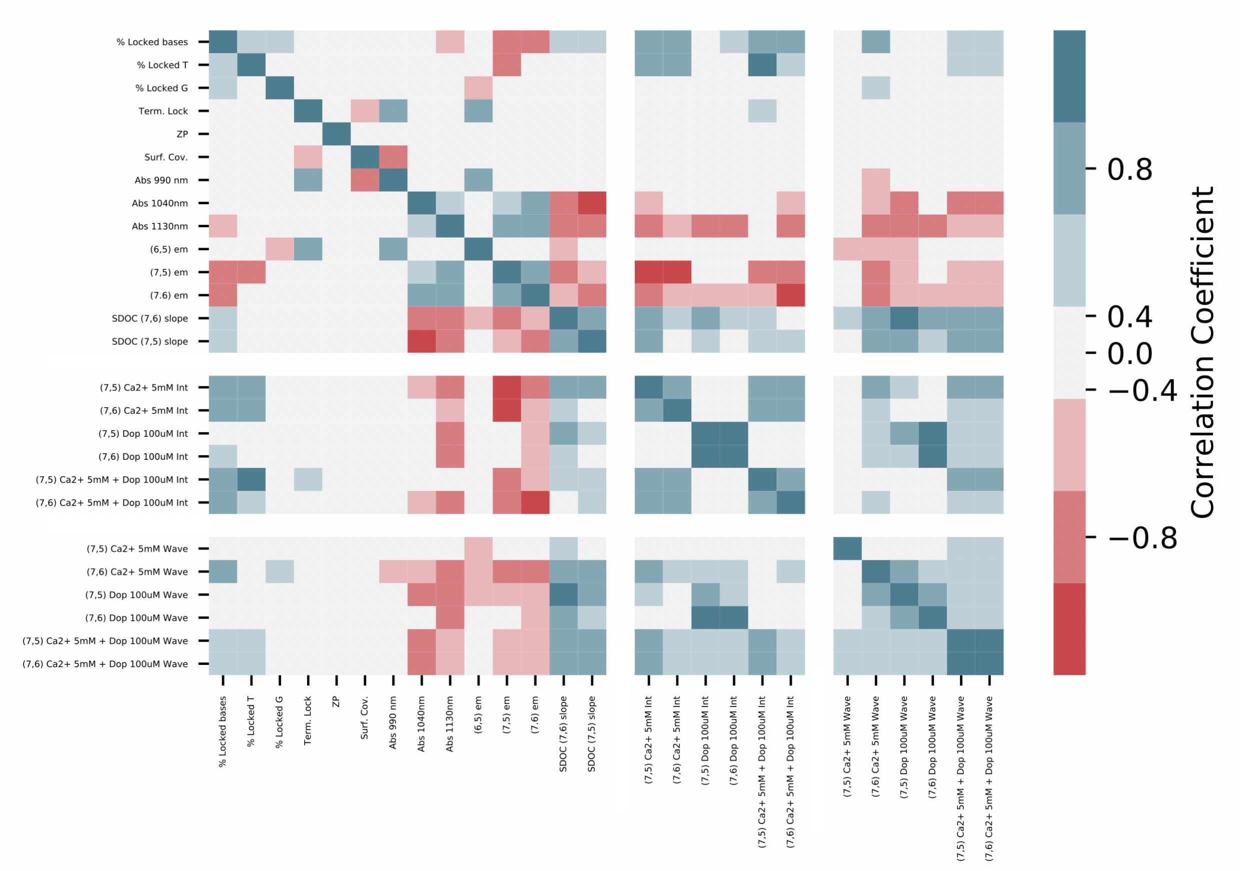


Figure 25: Data correlation matrix for the $(GT)_{15}$- and LNA-SWCNT for several different parameters measured. **Blue** hue indicates positive correlation and **red** hues indicate negative correlation.

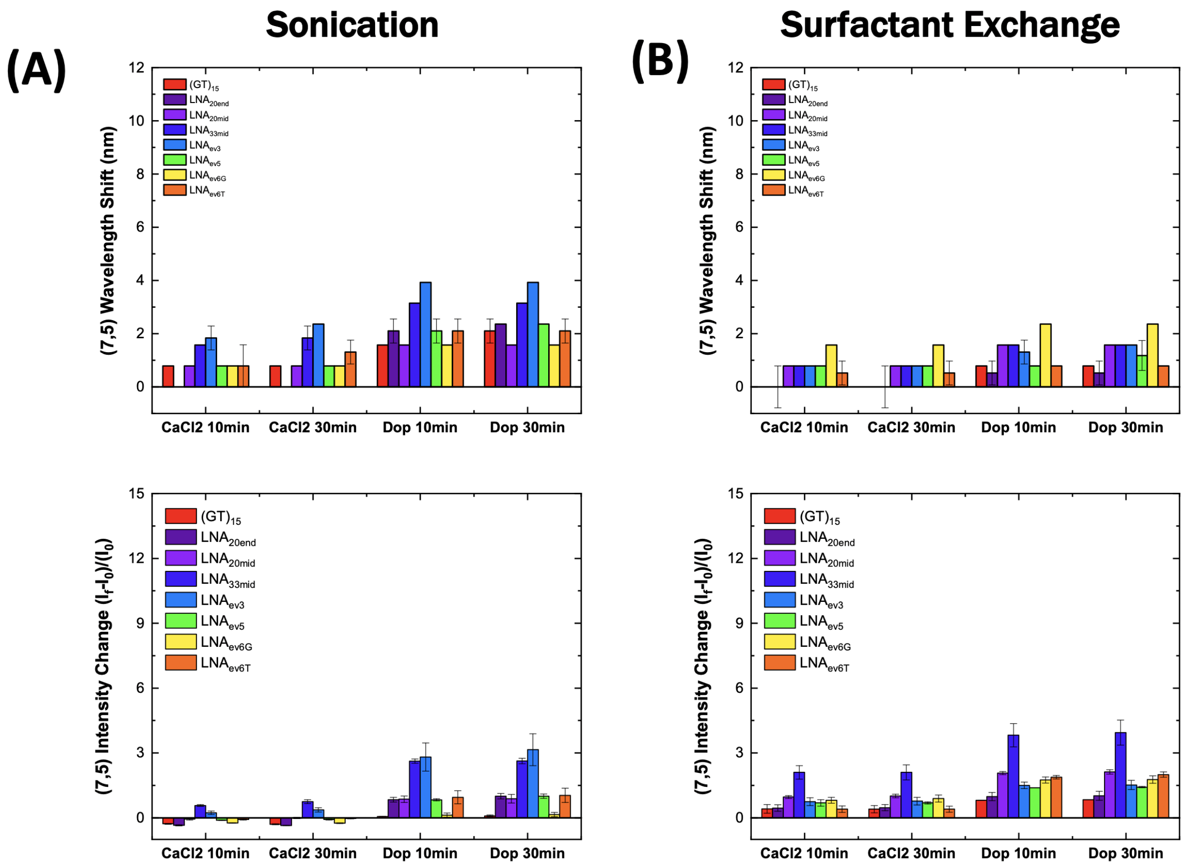


Figure 26: Comparison of the fluorescence response of $(GT)_{15}$- and LNA-SWCNTs prepared via either **(A)** sonication or **(B)** via a surfactant exchange protocol following the addition of 0.5 M $CaCl_2$ (final concentration: 5 mM) and 10 mM dopamine (final concentration: 100 $\mu$M, excitation: 660 nm). Intensity (**top**) and wavelength (**bottom**) response of the (7,5) peak following dopamine addition were calculated versus the initial spectrum (before initial $CaCl_2$ addition) for all time points. Error bars represent $1\sigma$ standard deviation (n = 3 technical replicates).

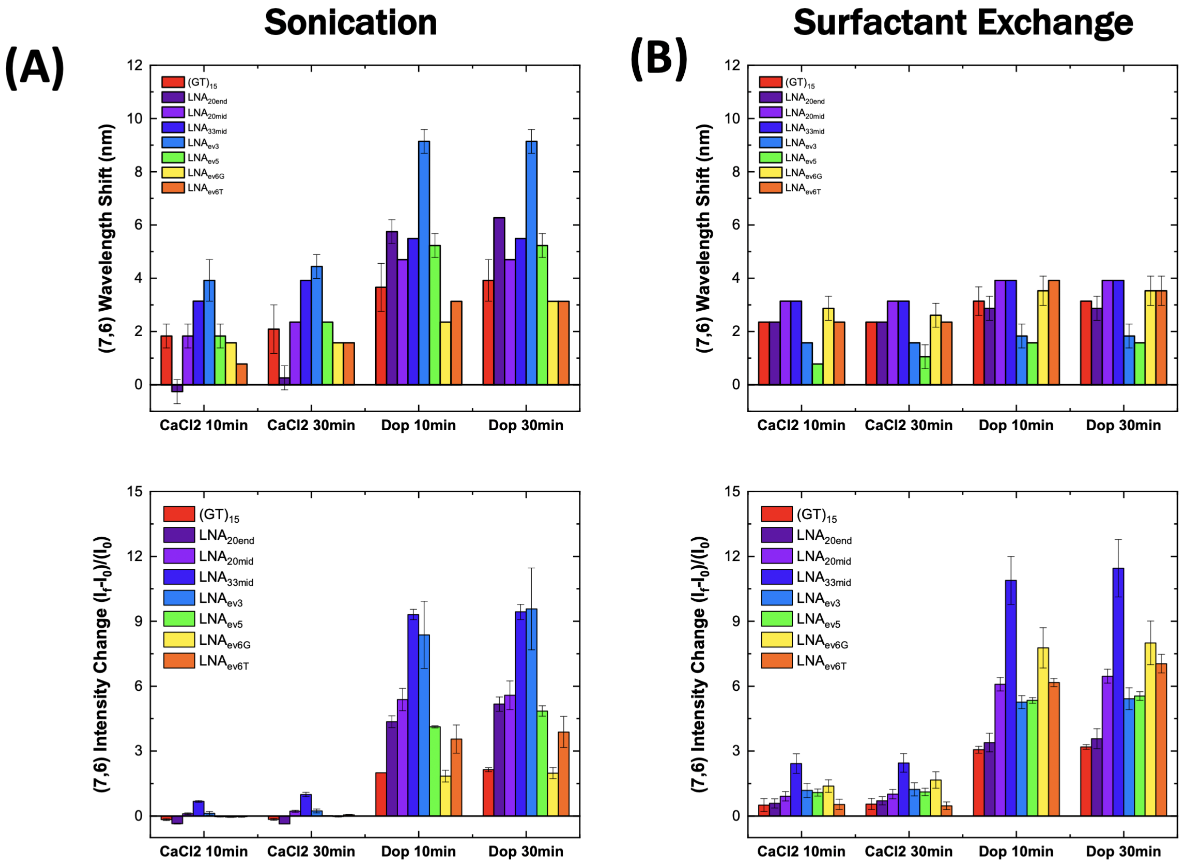


Figure 27: Comparison of the fluorescence response of $(GT)_{15}$- and LNA-SWCNTs prepared by either **(A)** sonication or **(B)** MeOH assisted surfactant exchange following the addition of 0.5 M $CaCl_2$ (final concentration: 5 mM) and 10 mM dopamine (final concentration: 100 $\mu$M, excitation: 660 nm). Intensity (**top**) and wavelength (**bottom**) response of the (7,6) peak following dopamine addition were calculated versus the initial spectrum (before initial $CaCl_2$ addition) for all time points. Error bars represent $1\sigma$ standard deviation (n = 3 technical replicates).

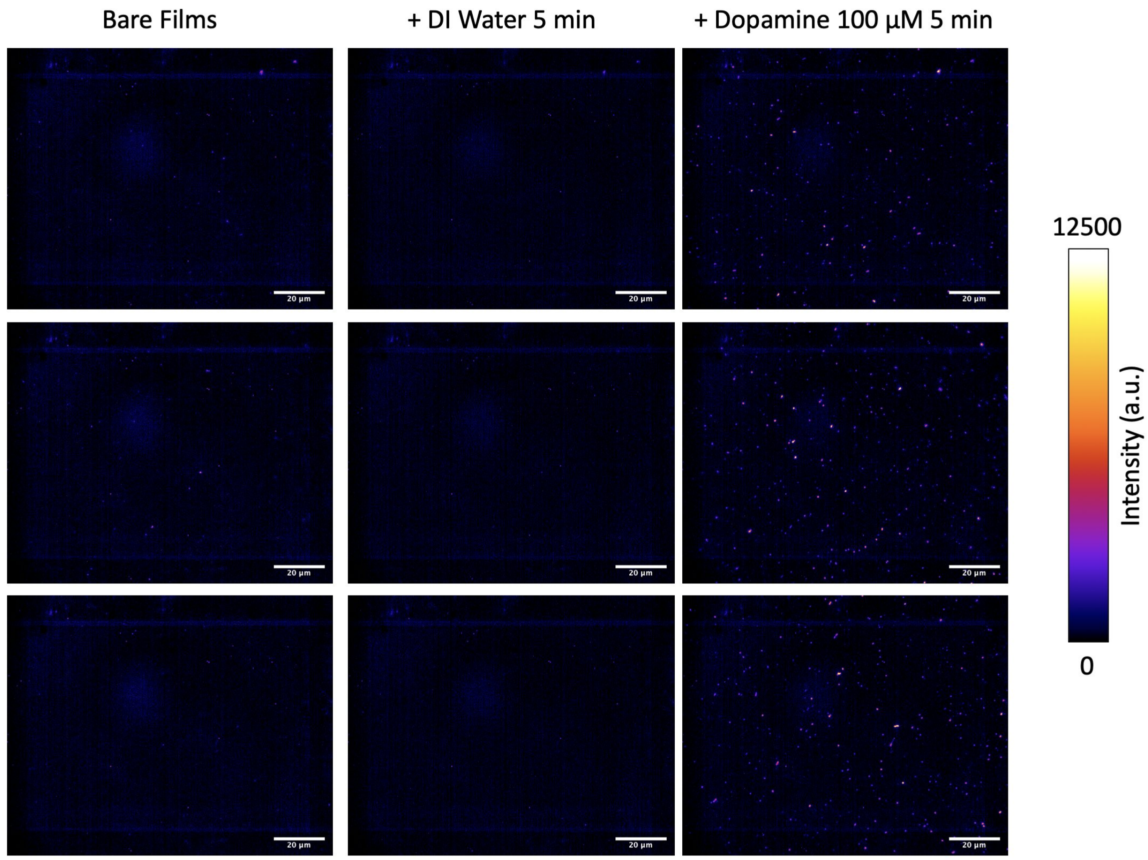


Figure 28: Single molecule response behavior of DNA-SWCNT sensor films towards dopamine (final concentration: 100 $\mu$M) in DI water. Three representative images of bare $(GT)_{15}$ sensor films (**left**), films coated with DI water (**middle**), and following the addition of dopamine (**right**). All samples were incubated for 5 min prior to acquiring the images (excitation: 780 nm, emission filter: 980 nm LP, exposure time: 1 s). Backgrounds were subtracted using a Gaussian blur filter (sigma = 50).

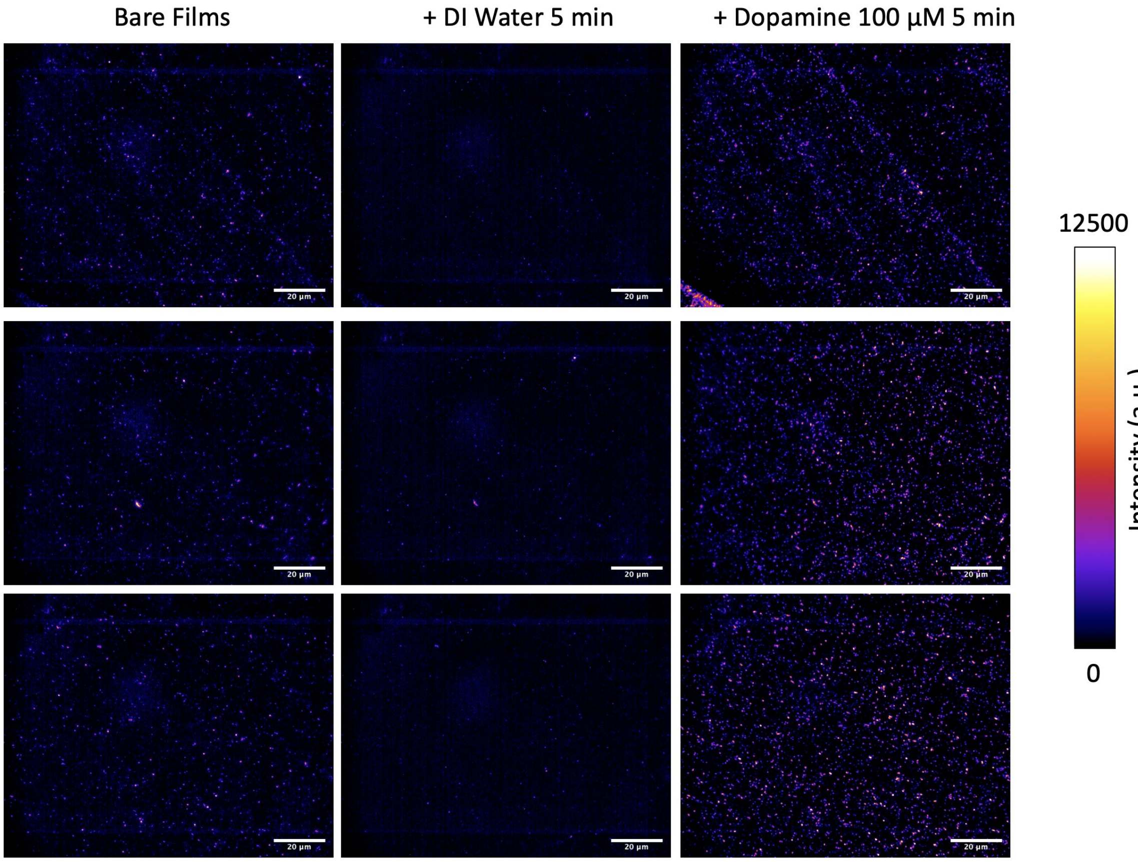


Figure 29: Single molecule response behavior of $LNA_{every5}$-SWCNT sensor films towards dopamine (final concentration: 100 uM) in DI water. Three representative images of bare $LNA_{every5}$ sensor films (**left**), films coated with DI water (**middle**), and following the addition of dopamine (**right**). All samples were incubated for 5 min prior to acquiring the images (excitation: 780 nm, emission filter: 980 nm LP, exposure time: 1 s). Backgrounds were subtracted using a Gaussian blur filter (sigma = 50).

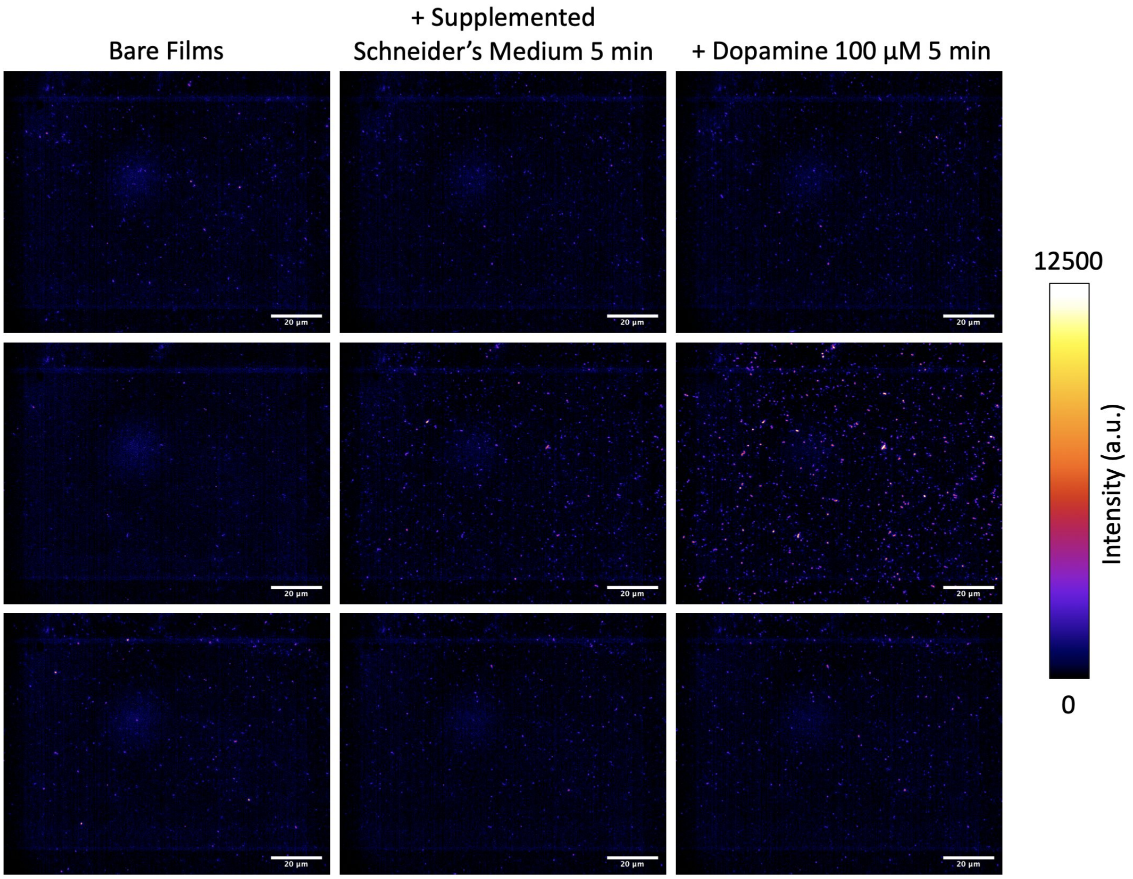


Figure 30: Single molecule response behavior of DNA-SWCNT sensor films towards dopamine (final concentration: 100 $\mu$M) in supplemented Schneider's medium (10% FBS, 1% Pen/Strep). Three representative images of bare $(GT)_{15}$ sensor films (**left**), films coated with supplemented Schneider's medium (**middle**), and following the addition of dopamine (**right**). All samples were incubated for 5 min prior to acquiring the images (excitation: 780 nm, emission filter: 980 nm LP, exposure time: 1 s). Backgrounds were subtracted using a Gaussian blur filter (sigma = 50).

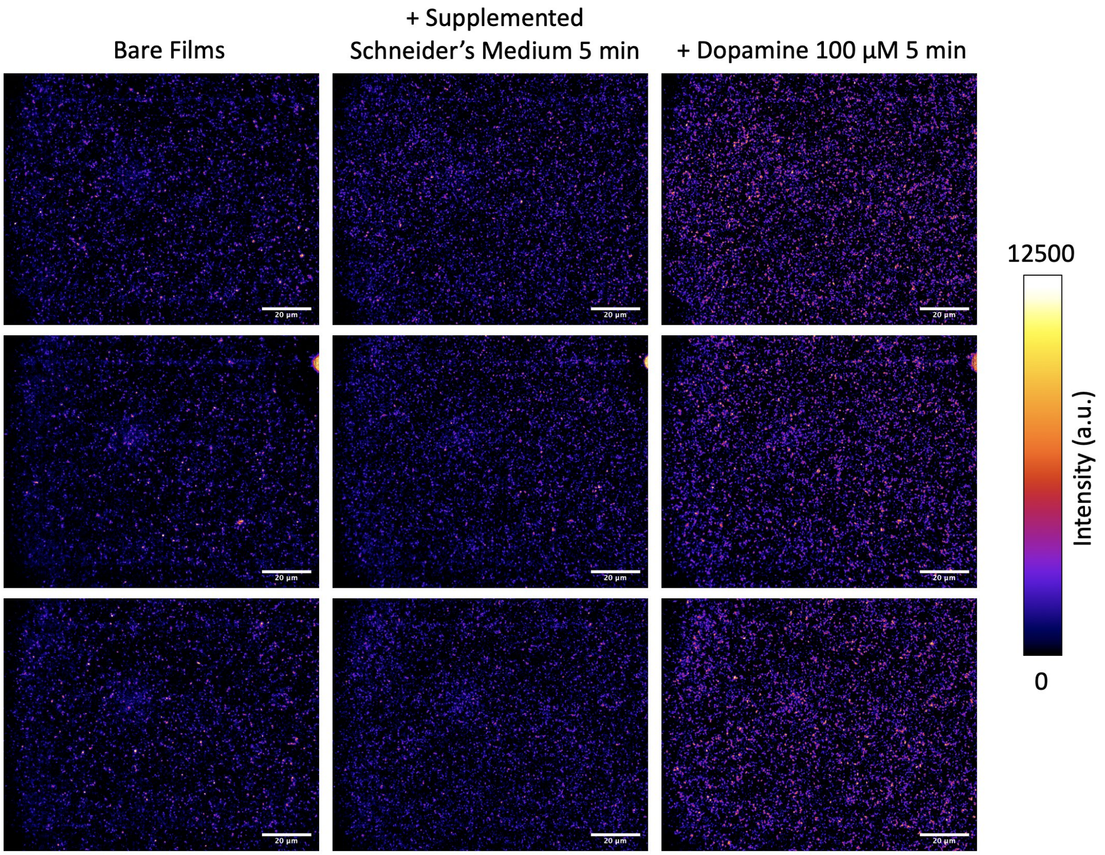


Figure 31: Single molecule response behavior of $LNA_{every5}$-SWCNT sensor films towards dopamine (final concentration: 100 $\mu$M) in supplemented Schneider's medium (10% FBS, 1% Pen/Strep). Three representative images of bare $LNA_{every5}$ sensor films (**left**), films coated with supplemented Schneider's medium (**middle**), and following the addition of dopamine (**right**). All samples were incubated for 5 min prior to acquiring the images (excitation: 780 nm, emission filter: 980 nm LP, exposure time: 1 s). Backgrounds were subtracted using a Gaussian blur filter (sigma = 50).

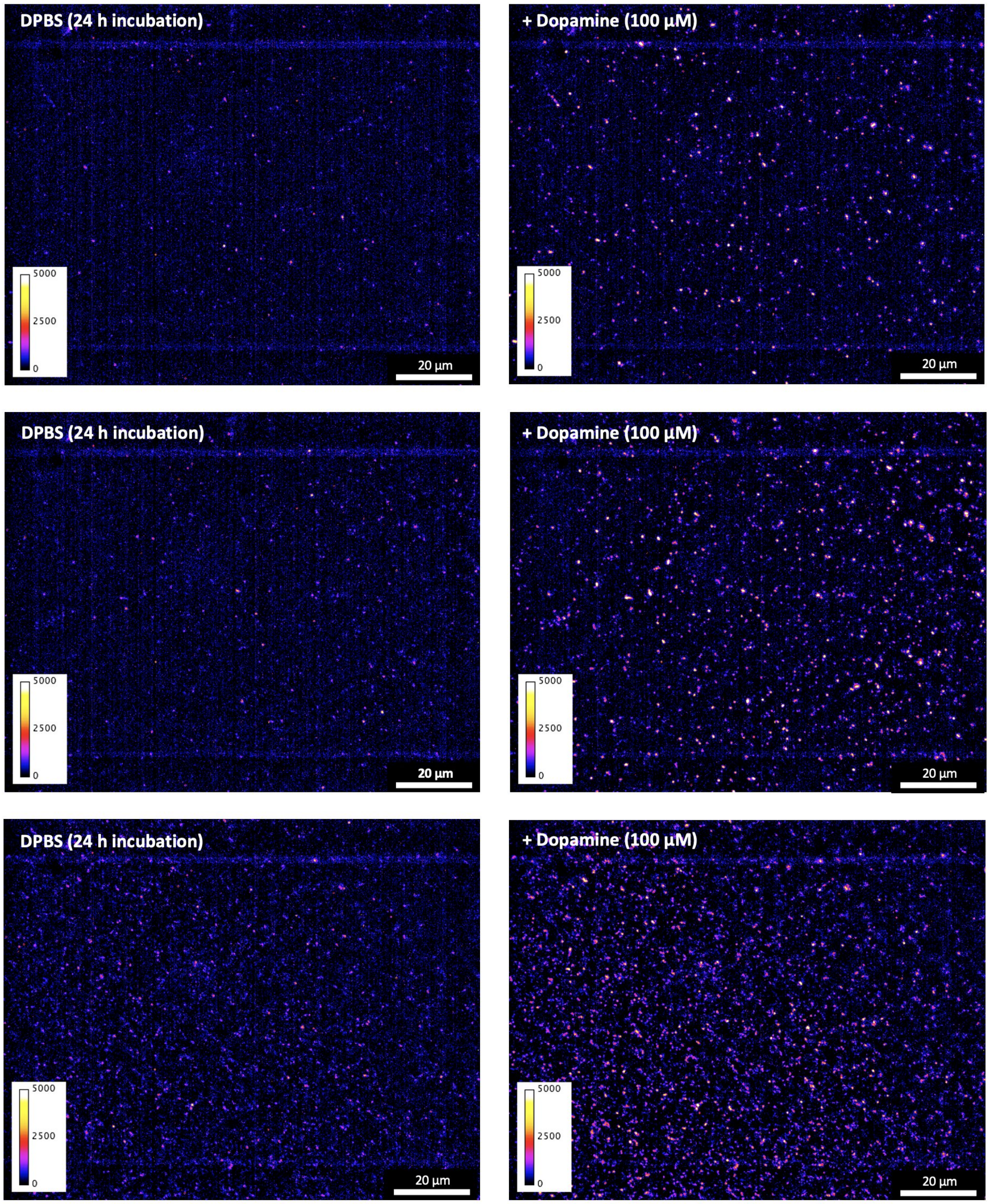


Figure 32: Single molecule response behavior of $LNA_{every5}$-SWCNT sensor films in DPBS towards dopamine (final concentration: 100 $\mu$M). All films were incubated for 24 h at room temperature prior to measurement. Three representative images of $LNA_{every5}$ sensor films before (**left**) and after (**right**) the addition of dopamine. All samples were incubated for 15 min post-dopamine addition prior to acquiring the images (excitation: 780 nm, emission filter: 980 nm LP, exposure time: 1 s). Backgrounds were subtracted using a Gaussian blur filter (sigma = 25).

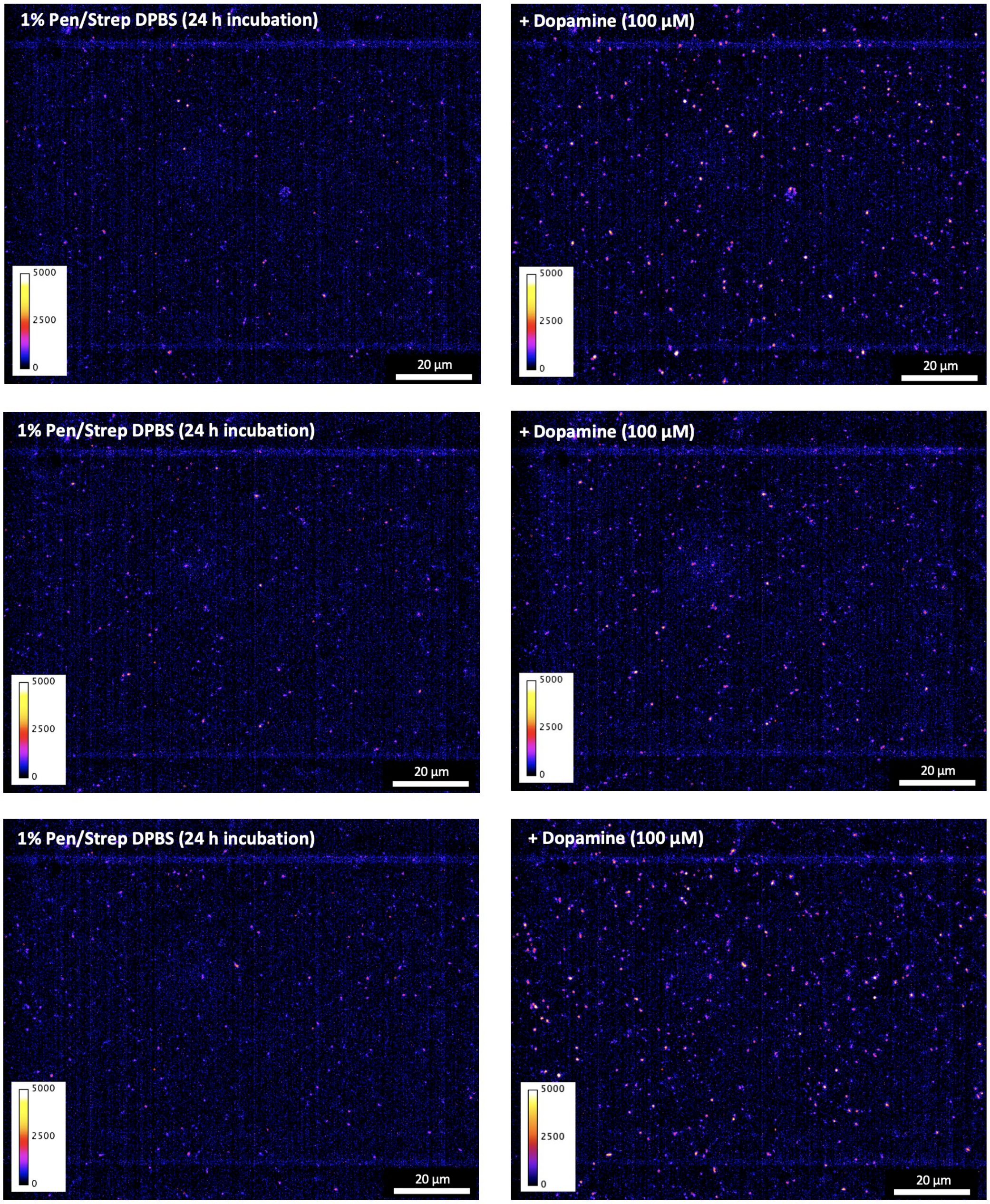


Figure 33: Single molecule response behavior of $LNA_{every5}$-SWCNT sensor films in 1% Pen/Strep in DPBS (Rinaldini solution) towards dopamine (final concentration: 100 $\mu$M). All films were incubated for 24 h at room temperature prior to measurement. Three representative images of $LNA_{every5}$ sensor films before (**left**) and after (**right**) the addition of dopamine. All samples were incubated for 15 min post-dopamine addition prior to acquiring the images (excitation: 780 nm, emission filter: 980 nm LP, exposure time: 1 s). Backgrounds were subtracted using a Gaussian blur filter (sigma = 25).

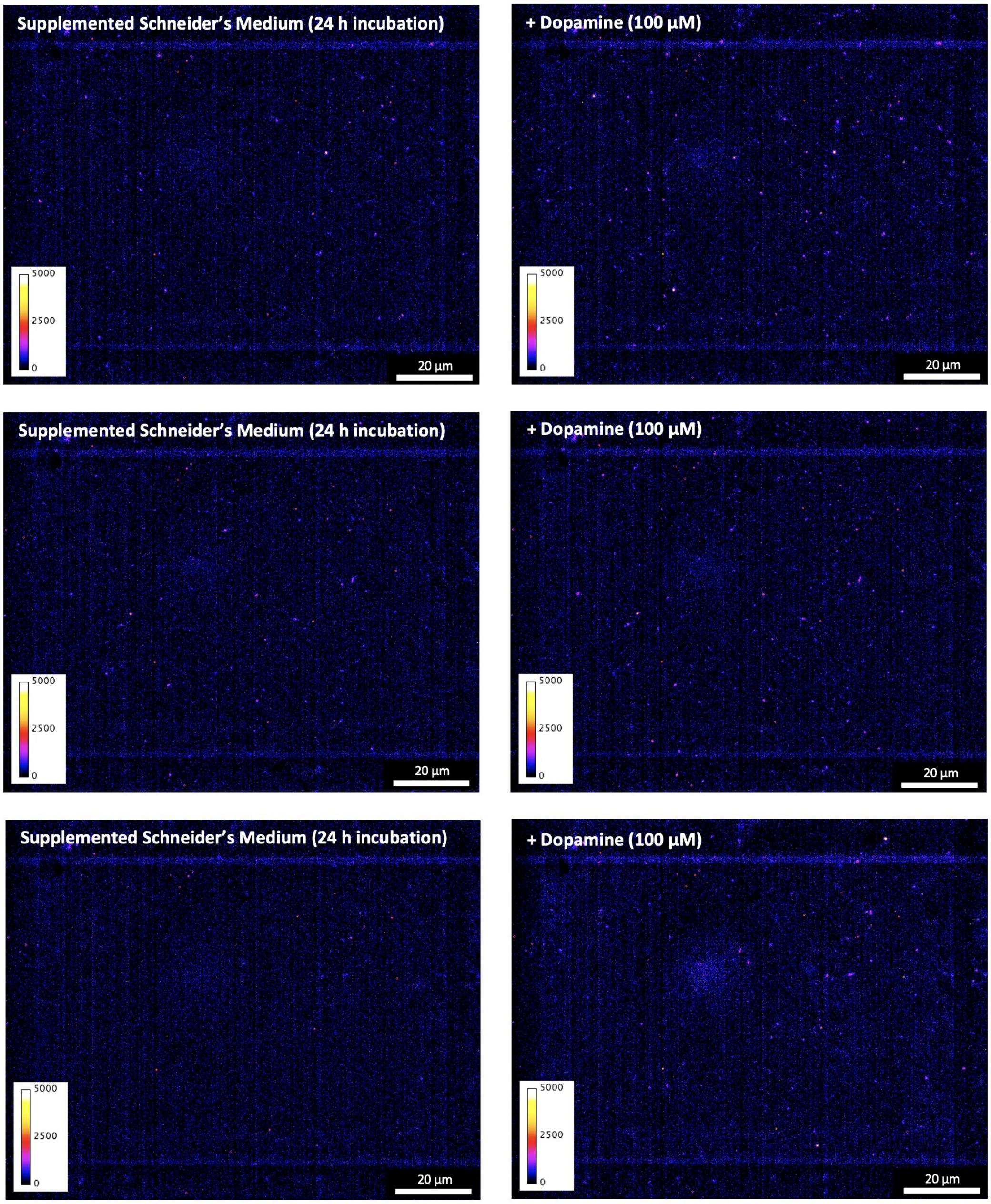


Figure 34: Single molecule response behavior of $LNA_{every5}$-SWCNT sensor films in supplemented Schneider's medium towards dopamine (final concentration: 100 $\mu$M). All films were incubated for 24 h at room temperature prior to measurement. Three representative images of $LNA_{every5}$ sensor films before (**left**) and after (**right**) the addition of dopamine. All samples were incubated for 15 min post-dopamine addition prior to acquiring the images (excitation: 780 nm, emission filter: 980 nm LP, exposure time: 1 s). Backgrounds were subtracted using a Gaussian blur filter (sigma = 25).

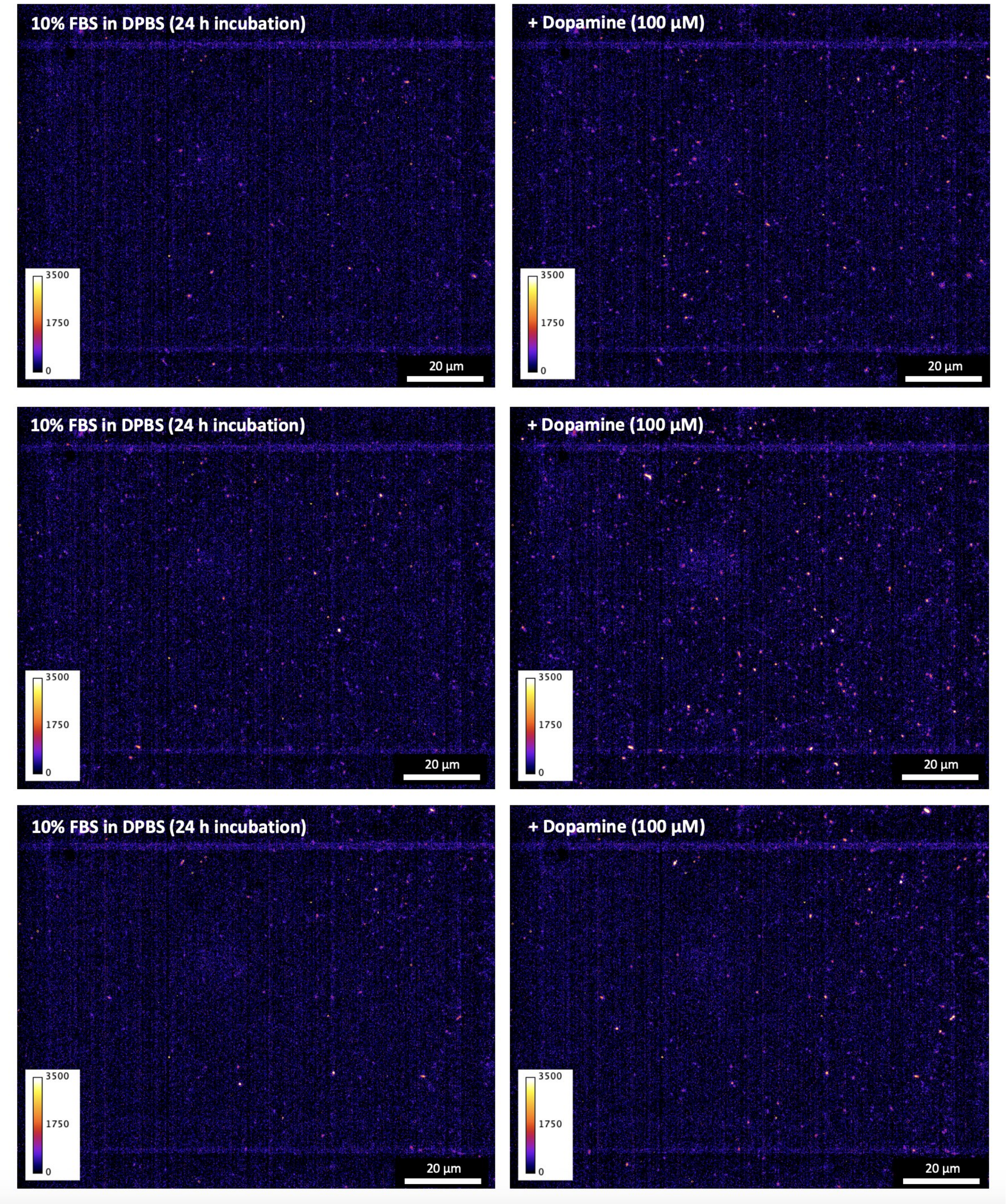


Figure 35: Single molecule response behavior of $LNA_{every5}$-SWCNT sensor films in 10% FBS (v/v) towards dopamine (final concentration: 100 $\mu$M). All films were incubated for 24 h at room temperature prior to measurement. Three representative images of $LNA_{every5}$ sensor films before (**left**) and after (**right**) the addition of dopamine. All samples were incubated for 15 min post-dopamine addition prior to acquiring the images (excitation: 780 nm, emission filter: 980 nm LP, exposure time: 1 s). Backgrounds were subtracted using a Gaussian blur filter (sigma = 25).

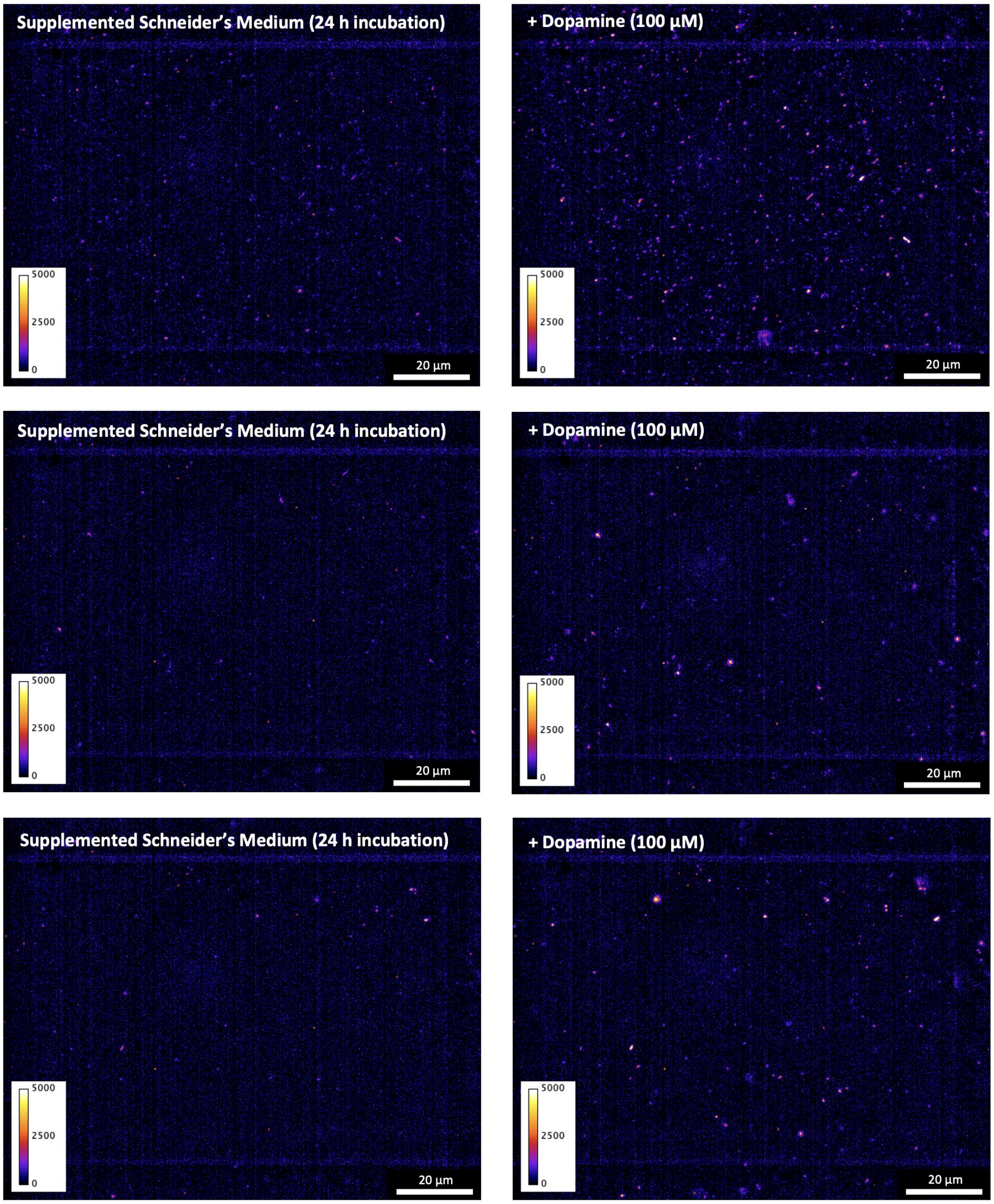


Figure 36: Single molecule response behavior of denser $LNA_{every5}$-SWCNT sensor films in supplemented Schneider's medium towards dopamine (final concentration: 100 $\mu$M). All films were incubated for 24 h at room temperature prior to measurement. Three representative images of $LNA_{every5}$ sensor films before (**left**) and after (**right**) the addition of dopamine. All samples were incubated for 15 min post-dopamine addition prior to acquiring the images (excitation: 780 nm, emission filter: 980 nm LP, exposure time: 1 s). Backgrounds were subtracted using a Gaussian blur filter (sigma = 25).

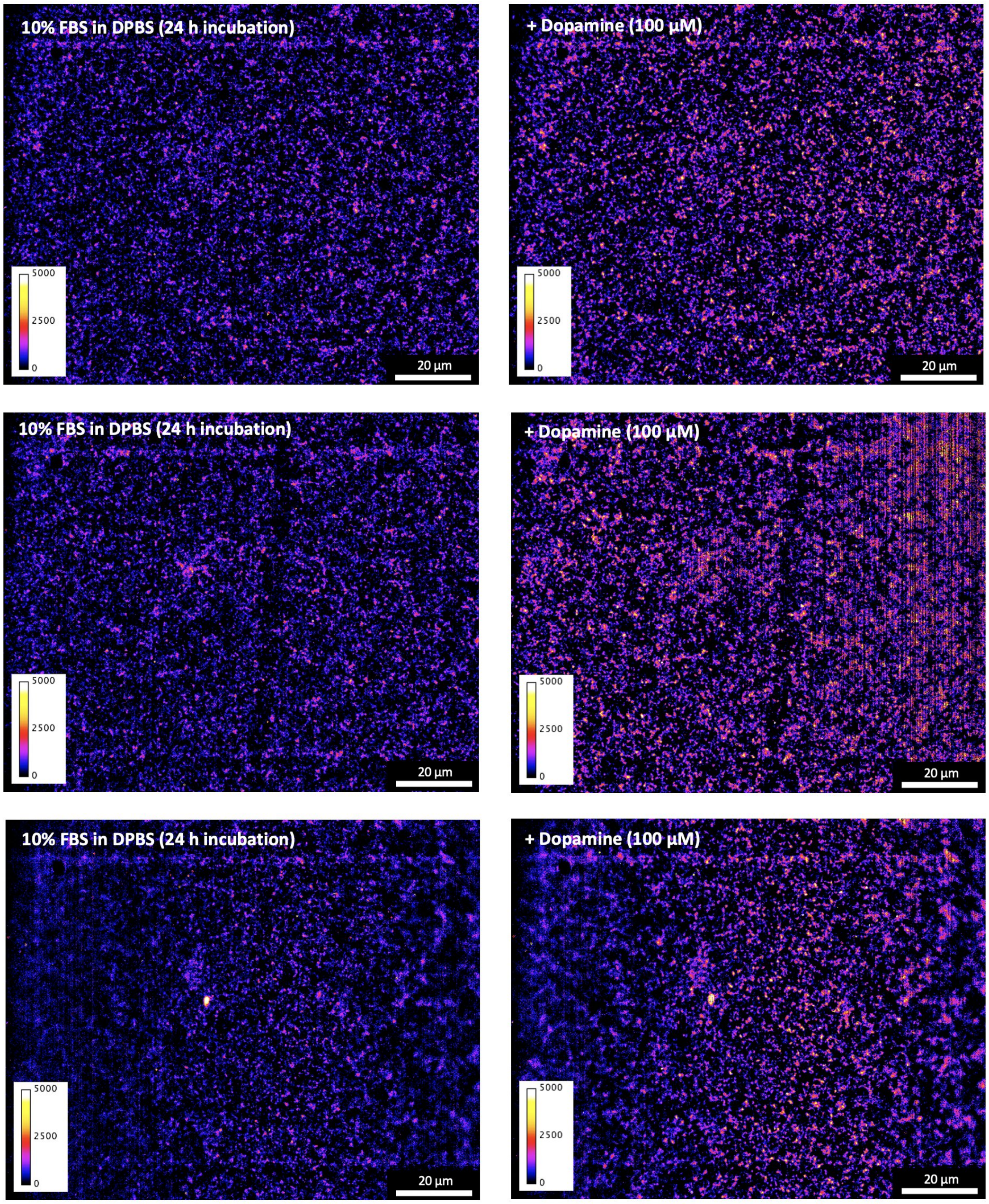


Figure 37: Single molecule response behavior of more ultrahigh density $LNA_{every5}$-SWCNT sensor films in 10% FBS (v/v) towards dopamine (final concentration: 100 $\mu$M). All films were incubated for 24 h at room temperature prior to measurement. Three representative images of $LNA_{every5}$ sensor films before (**left**) and after (**right**) the addition of dopamine. All samples were incubated for 15 min post-dopamine addition prior to acquiring the images (excitation: 780 nm, emission filter: 980 nm LP, exposure time: 1 s). Backgrounds were subtracted using a Gaussian blur filter (sigma = 25).

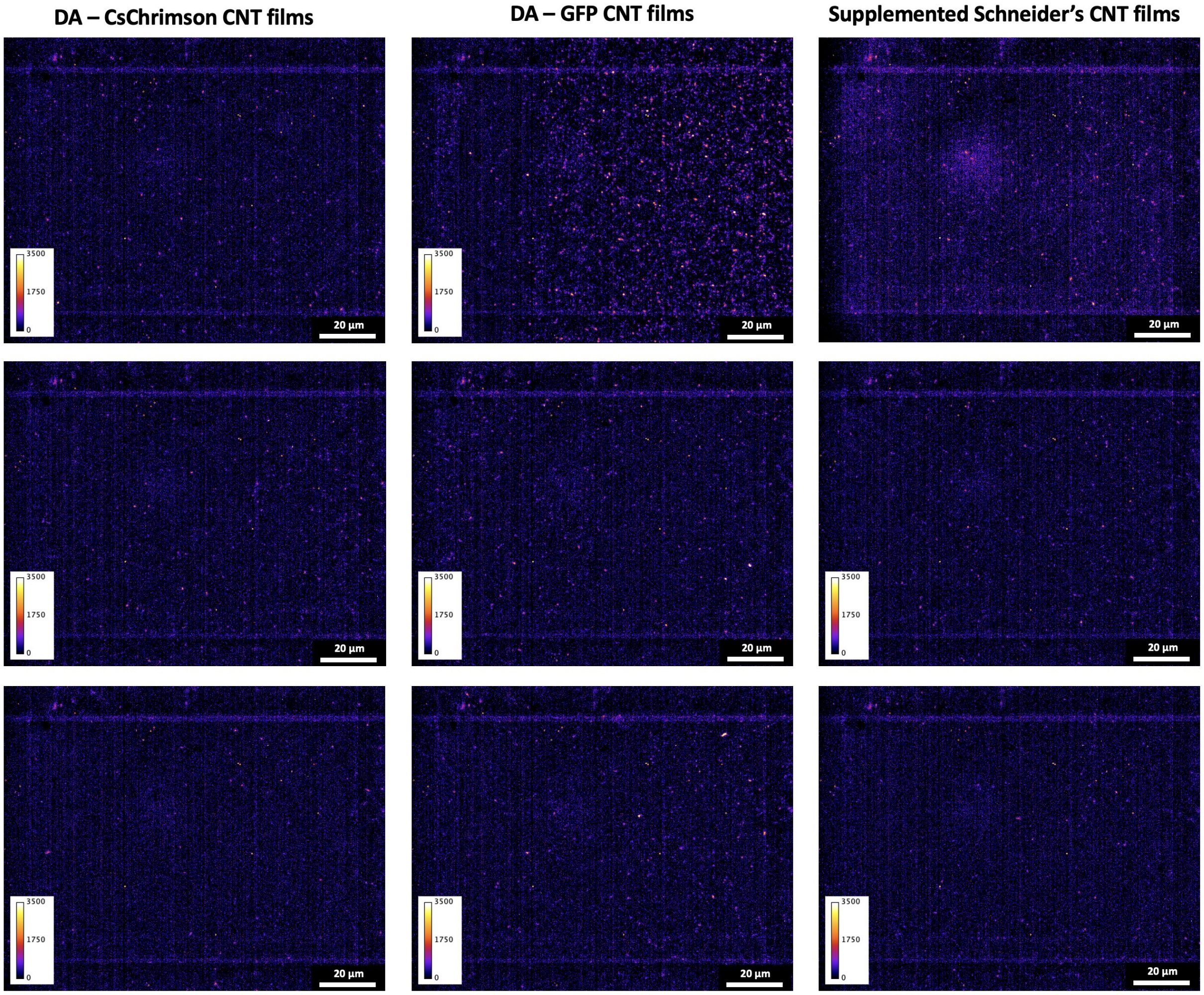


Figure 38: Comparison of $LNA_{every5}$-SWCNT sensor films following 48 h of incubation in supplemented Schneider's medium both with and without cells (excitation: 780 nm, emission filter: 980 nm LP, exposure time: 1 s). *Drosophila* neuronal cell cultures containing transgenic dopamine neurons (representing approximately 1% of the sample) induced to express either GFP (DA-GFP) or CsChrimson (DA-CsChrimson) were grown on the nanotube films for 48 h at room temperature in supplemented Schneider's medium. Films incubated in supplemented Schneider's medium at room temperature for 48 h without cells were used for comparison to examine whether there was any change in the fluorescence properties of the nanotubes due to the presence of cells. Backgrounds were subtracted using a Gaussian blur filter (sigma = 25).

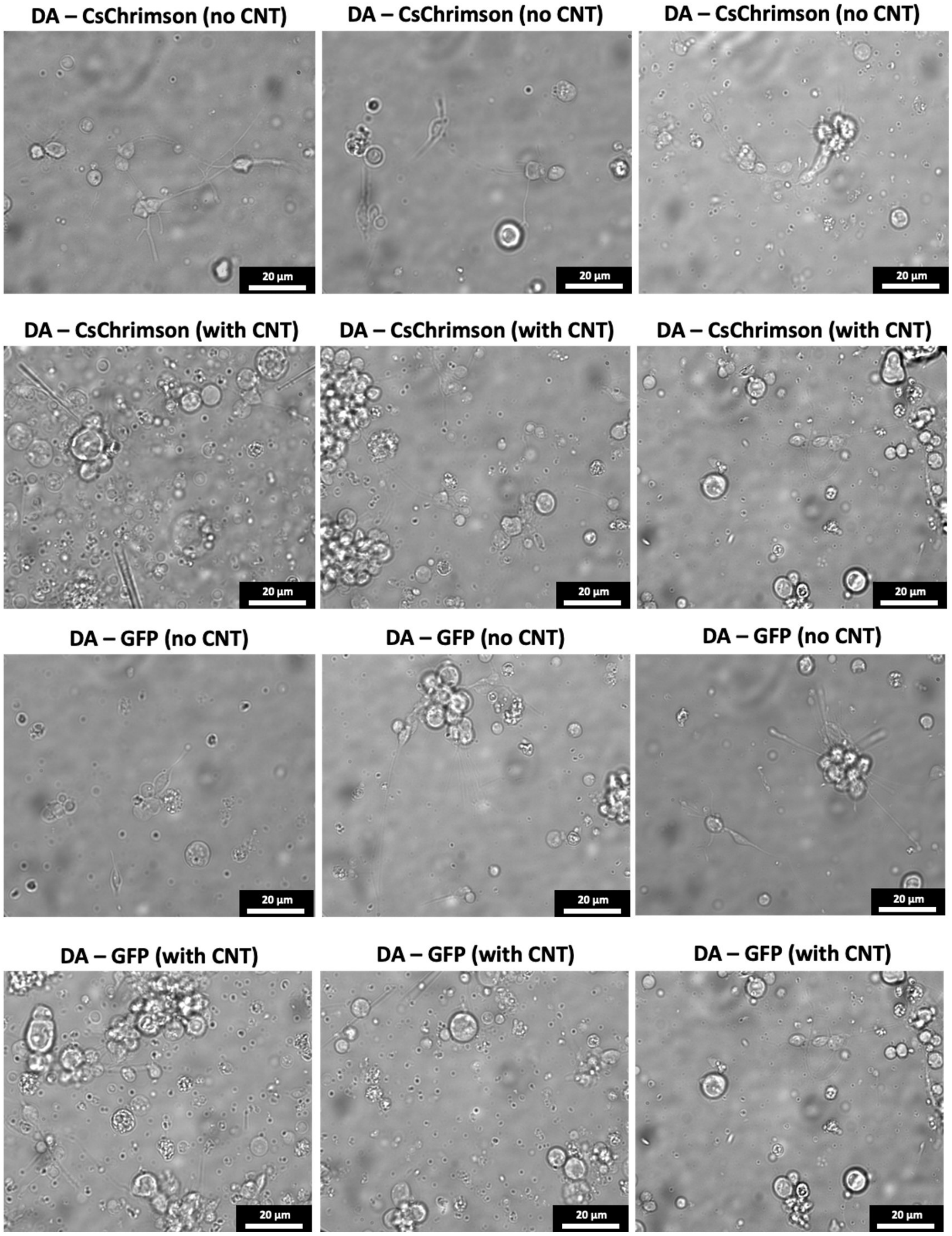


Figure 39: *Drosophila* neuronal cell cultures containing transgenic dopamine neurons (representing approximately 1% of the sample) were induced to express either GFP (DA-GFP) or CsChrimson (DA-CsChrimson). White light visible widefield images of the neurons were acquired following 48h of growth on poly-L-lysine coated glass petri dishes (**no CNT**) or $LNA_{every5}$-SWCNT covered poly-L-lysine coated glass petri dishes (**with CNT**) to examine the effect of nanotubes on cell growth.

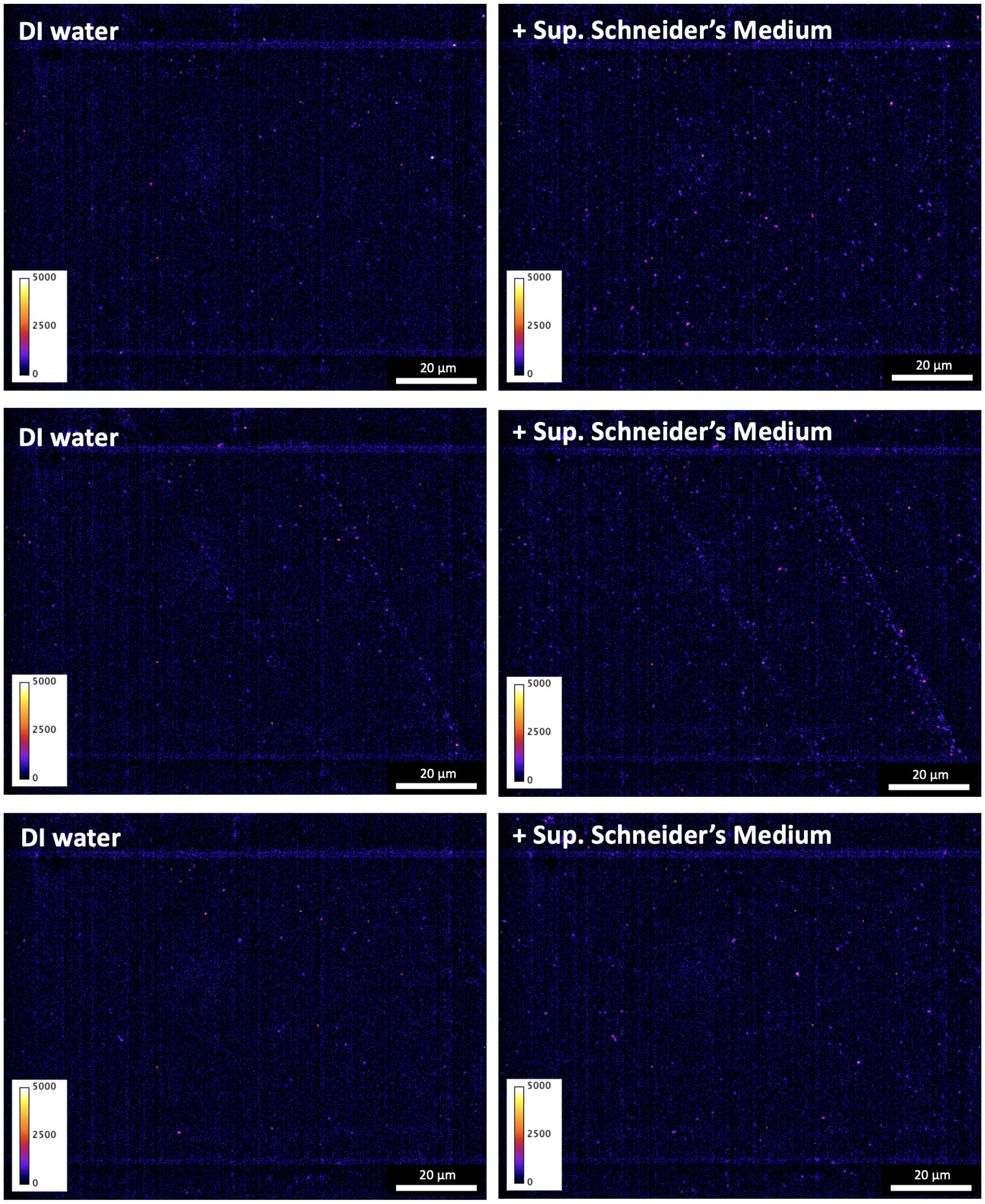


Figure 40: Single molecule response of $LNA_{every5}$-SWCNT sensor films in DI water to the addition of supplemented Schneider's medium. Films were initially wetted with 100$\mu$L of DI water and incubated for 15 min prior to adding 10 $\mu$L of supplemented Schneider's medium. Three representative images of $LNA_{every5}$ sensor films before (**left**) and after (**right**) the addition of medium were acquired. All samples were incubated for 15 min post medium addition prior to acquiring the images (excitation: 780 nm, emission filter: 980 nm LP, exposure time: 1 s). Backgrounds were subtracted using a Gaussian blur filter (sigma = 25).

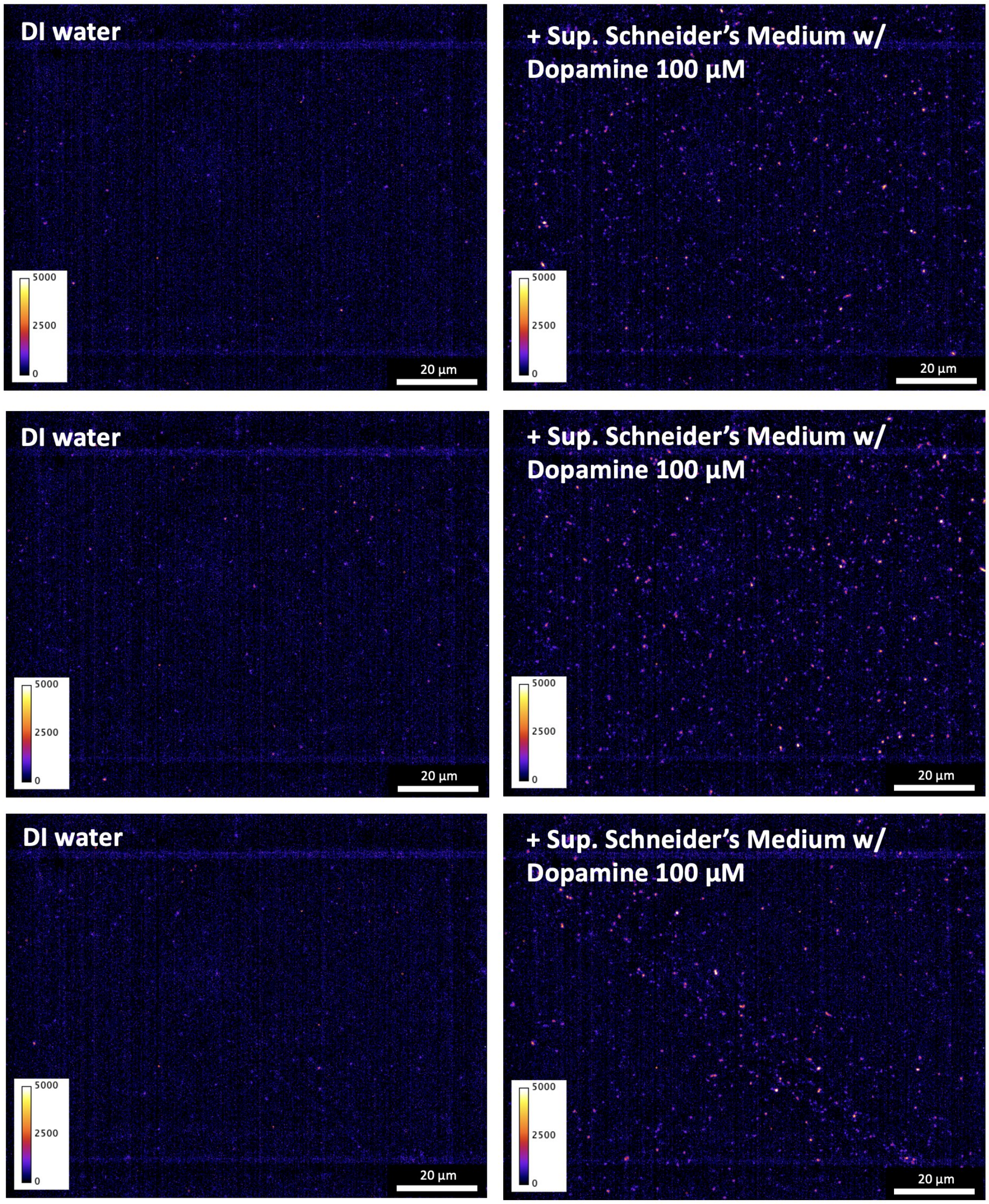


Figure 41: Single molecule response of $LNA_{every5}$-SWCNT sensor films in DI water to the addition of supplemented Schneider's medium spiked with 100 $\mu$M dopamine (final concentration: = 9.1 $\mu$M). Films were initially wetted with 100$\mu$L of DI water and incubated for 15 min prior to adding 10 $\mu$L of supplemented Schneider's medium freshly prepared with 100 $\mu$M dopamine. Three representative images of $LNA_{every5}$ sensor films before (**left**) and after (**right**) the addition of medium were acquired. All samples were incubated for 15 min post medium addition prior to acquiring the images (excitation: 780 nm, emission filter: 980 nm LP, exposure time: 1 s). Backgrounds were subtracted using a Gaussian blur filter (sigma = 25).

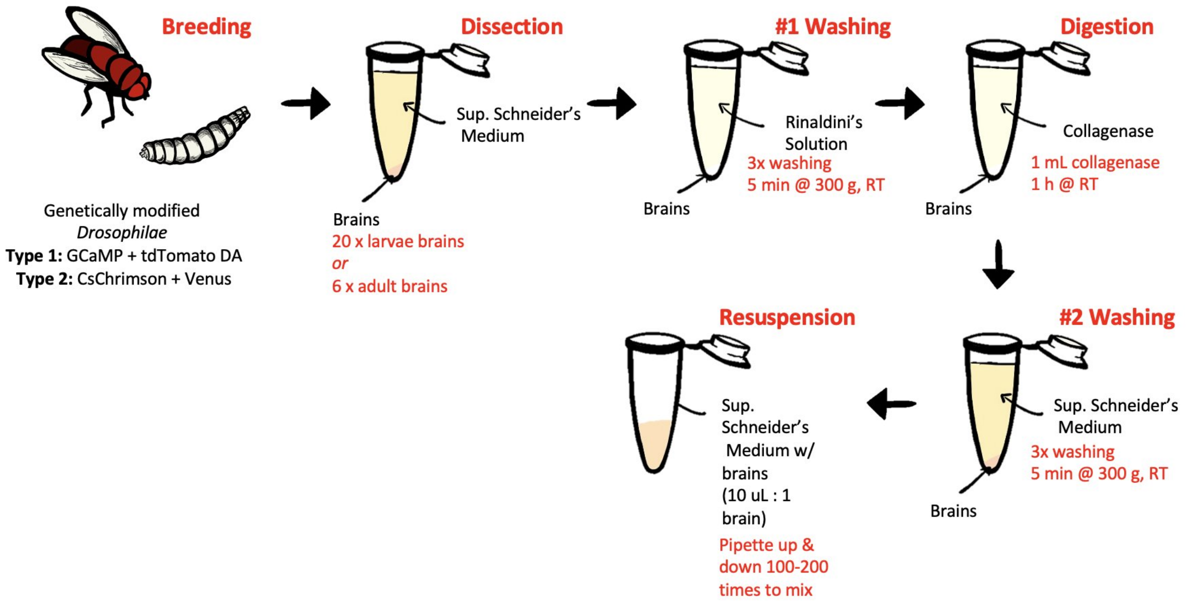


Figure 42: Schematic of the protocol used to extract *Drosophila* neurons from adult or larvae brains. Neurons resuspended in supplemented Schneider's medium could then be plated in either 96-well plates or glass-bottom petri dishes.
